# Comparative cryo-EM studies reveal structural aspects of host specificity for tobamoviruses

**DOI:** 10.64898/2026.11.02.763805

**Authors:** Ananya Chatterjee, Aravind Venkatasubramanian, A. Abdul Kader Jailani, Prantik Mazumder, Sandeep Singh, Somnath Basu, Uddipan Das, Anirban Roy, Bikash Mandal, Partha Pratim Datta

## Abstract

Tobamoviruses are a group of economically important plant viruses exhibiting extreme stability and persistence. Despite high morphological similarity, they show extensive host-family specificity, induce a wide range of symptoms, and inflict significant global yield losses in fruit and vegetable crops. Composed of a rod-shaped helical assembly of identical coat proteins encasing a single-stranded RNA, tobamoviruses have long served as fundamental models for understanding helical assemblies. Here we report the cryo-EM structures of two viruses from this genus, cucumber green mottle mosaic virus (CGMMV) and tomato mottle mosaic virus (ToMMV), at 2.86 Å and 2.71 Å resolution, respectively, revealing divergent packing strategies within the virion assembly. Whereas the solanaceous-infecting ToMMV closely mirrors genetically-similar canonical tobacco mosaic virus (TMV), CGMMV employs a specialized hydrophobic packing strategy based at the solvent-exposed inner loop lining the solvent-exposed central canal. Comparative biophysical and molecular dynamics analyses from the reconstructed structures reveal more stable and compact protein packing for CGMMV at both the subunit and assembly levels. Furthermore, we identify a conserved electrostatic pentapeptide motif (residues 113–117) on the lower radial alpha helix that serves as an additional RNA-stabilizing region, harbouring a clade-defining Lys113 synapomorphy in cucurbit-infecting tobamoviruses. Together, our findings elucidate how subtle electrostatic tuning and divergent hydrophobic packing strategies drive differential virion stability and host-family adaptation within a strictly conserved helical scaffold.

**Significance statement:** Understanding how plant viruses adapt to diverse host species while maintaining physical stability is crucial for combating crop loss. Tobamoviruses are hyper-stable, rod-shaped plant viruses with highly conserved structures, yet they infect vastly different host families. By solving high-resolution cryo-EM structures of two representative tobamoviruses and comparing them with existing structural information, we show that despite near-identical overall shapes, these viruses employ distinct internal packing mechanisms and electrostatic strategies to stabilize their viral RNA. This work reveals how subtle sequence variations tune internal hydrophobic and electrostatic networks, providing fundamental insights into the evolutionary principles that allow these viruses to drive host diversification without altering their overarching structural architecture.

## Introduction

Macromolecular stability in supramolecular assemblies is governed by the geometry, extent, and energetics of intermolecular interfaces. Within large protein complexes, closely packed interfaces maximize cooperative non-covalent interactions, including hydrogen bonding, salt bridges, hydrophobic contacts, and van der Waals forces, thereby minimizing solvent exposure and stabilizing the quaternary architecture(1, 2). In repetitive helical systems such as viral capsids, subtle alterations in subunit packing efficiency, buried surface area, or local hydrophobic clustering propagate across the entire lattice(3, 4). Consequently, fine-scale structural variations at individual coat protein interfaces can significantly influence intrinsic thermodynamic resilience, dynamic stability, and uncoating behaviours without necessarily altering the global virion architecture.

Members of the plant virus genus *Tobamovirus* serve as classical models for exploring structural stability in helical assemblies(5, 6). These rod-shaped virions consist of approximately 2,000-2,200 identical coat protein (CP) subunits arranged helically around a single-stranded genomic RNA(7). *Tobacco mosaic virus* (TMV) has long served as the canonical benchmark for this architectural class. Decades of structural and biochemical studies on TMV have revealed a tightly cooperative network of protein-protein and protein-RNA interactions, demonstrating how localized carboxylate clusters, electrostatic environments, and bound divalent cations modulate capsid integrity and genome uncoating(8–14). However, because these structural insights stem predominantly from TMV, the extent to which these specific interaction networks are conserved across divergent *Tobamovirus* lineages remains underrepresented.

To evaluate structural variation across the genus, we performed a comprehensive phylogenetic analysis of 33 ICTV-recognized *Tobamovirus* species using coat protein sequences (Fig. 1A). The resulting tree topology reveals an evolutionary bifurcation, where tobamoviruses cluster into distinct host-family-associated clades (Fig. 1A). Occupying one major clade are the cucurbit-infecting tobamoviruses, represented prototypically by cucumber green mottle mosaic virus (CGMMV)(15). CGMMV is a high-consequence agricultural pathogen known for its exceptional environmental durability. It retains infectivity in irrigation water networks, agricultural soils, and seed coats for extended periods, exhibiting thermal inactivation points between 85°-100°C, temperatures that destroy seed endosperm health and prevent heat-based sterilization(16, 17). CGMMV outbreaks cause yield losses exceeding 15% in cucumber and up to 50% in watermelon crops worldwide(18–21).

**Figure 1:**
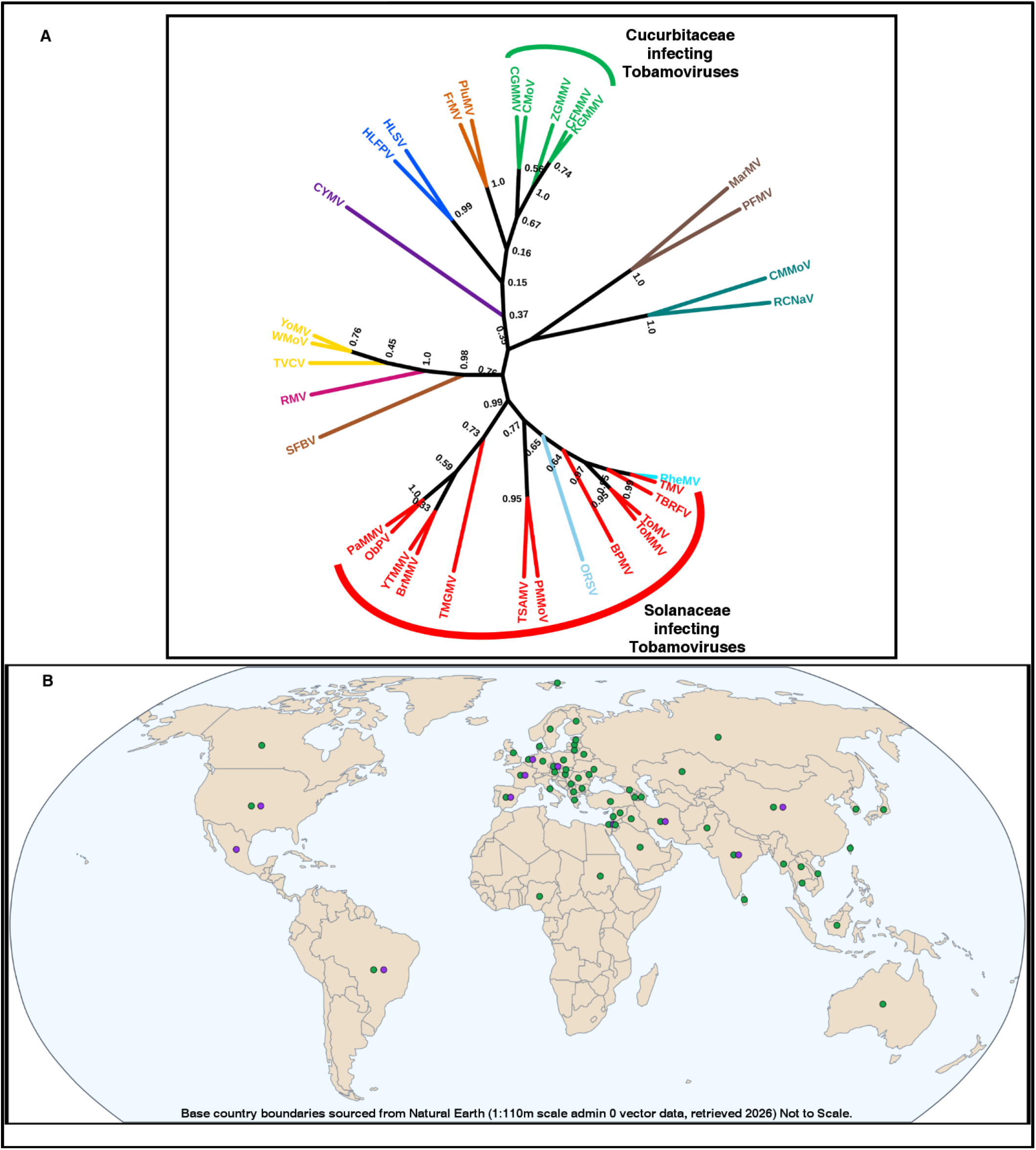
Evolutionary classification and global distribution of tobamoviruses. The maximum-likelihood evolutionary tree inferred from coat protein amino acid sequences in (A) shows tobamoviruses infecting the Solanaceae and Cucurbitaceae families of plants lying at either end of the tree, making them suitable candidates to study and compare host-specific structural adaptations. CGMMV and ToMMV, the two viruses in this study have a global occurrence as shown in the world map in (B) with green dots representing countries with reported CGMMV cases and purple dots representing ToMMV cases. Global distribution of documented CGMMV and ToMMV occurrence (including active, localized, and historical outbreak records) compiled from EPPO Global database (accessed 2026)(22, 23) and CABI Compendium data(24, 25). **Note:** *The base map (dataset source: Natural Earth, 2026, 1:110m scale; Kelso and Patterson, 2010(26)) is not to scale and subject to data availability. The designations employed and the presentation of material on this map do not imply the expression of any opinion whatsoever on the part of the authors or publisher concerning the legal status of any country, territory, city, or area, or concerning the delimitation of its frontiers or boundaries.*

Anchoring the opposite lineage in the phylogeny are the solanaceous-host-infecting tobamoviruses. To conduct a controlled comparative study with CGMMV, we selected tomato mottle mosaic virus (ToMMV) as the representative solanaceous target. ToMMV shares close structural proximity with canonical TMV but represents an urgent, emerging global threat due to its ability to overcome established resistance loci, including the *Tm-2^2^/tm-2* ToMV resistance gene in tomato and *L* resistance alleles in pepper(27, 28). Infection by ToMMV leads to severe leaf deformation, stunted growth, and pale, deformed fruit identified as shoestring disease, resulting in severe commercial loss(29). Both viruses show rapid mechanical transmission and seed-borne persistence and have expanded globally to cause widespread outbreaks, emerging as significant threats to crop production worldwide (Fig. 1B).

Despite infecting different plant families and exhibiting distinct environmental stability, how these divergent viral lineages achieve necessary capsid robustness within a conserved helical architecture remains to be understood. Whether subtle sequence evolution across distinct clades yields altered internal packaging or localized structural variations within this conserved helical fold remains to be systematically explored. To address these questions, we determined high-resolution cryo-electron microscopy (cryo-EM) reconstructions of both CGMMV and ToMMV, establishing atomic models to conduct a systematic comparative analysis. Using CGMMV and ToMMV as representative endpoints of the Cucurbit- and Solanaceous-host-infecting lineages, we investigated how localized variations in inter-subunit packing, internal stabilization networks, and RNA-proximal charge distributions influence capsid integrity. By integrating structural analysis with computational assessments of coat protein behaviour, we reveal candidate lineage-associated strategies that influence localized capsid architecture across a common helical structural scaffold.

## Results

### 1. Global Architecture of CGMMV and ToMMV and Radial Compaction in cucurbit-infecting CGMMV

The structures of the viruses were determined from cryo-EM micrographs using the helical reconstruction method, at resolutions of 2.86 Å for CGMMV (EMD-60696) and 2.71 Å for ToMMV (EMD-63082) (Fig. 2A, B). In both virions, identical coat protein subunits assemble into a right-handed helix intercalated with single-stranded genomic RNA, enclosing a hollow central canal characteristic of tobamoviruses (Fig. 2C, F). Both virions measure ∼186–187 Å in outer diameter, but CGMMV exhibits a narrower central canal (40 Å) than ToMMV (43 Å). Helical symmetry parameters refine to an axial rise of 1.41 Å and an azimuthal twist of 22.0° for CGMMV, compared with an axial rise of 1.44 Å and a twist of 22.0° for ToMMV.

**Figure 2:**
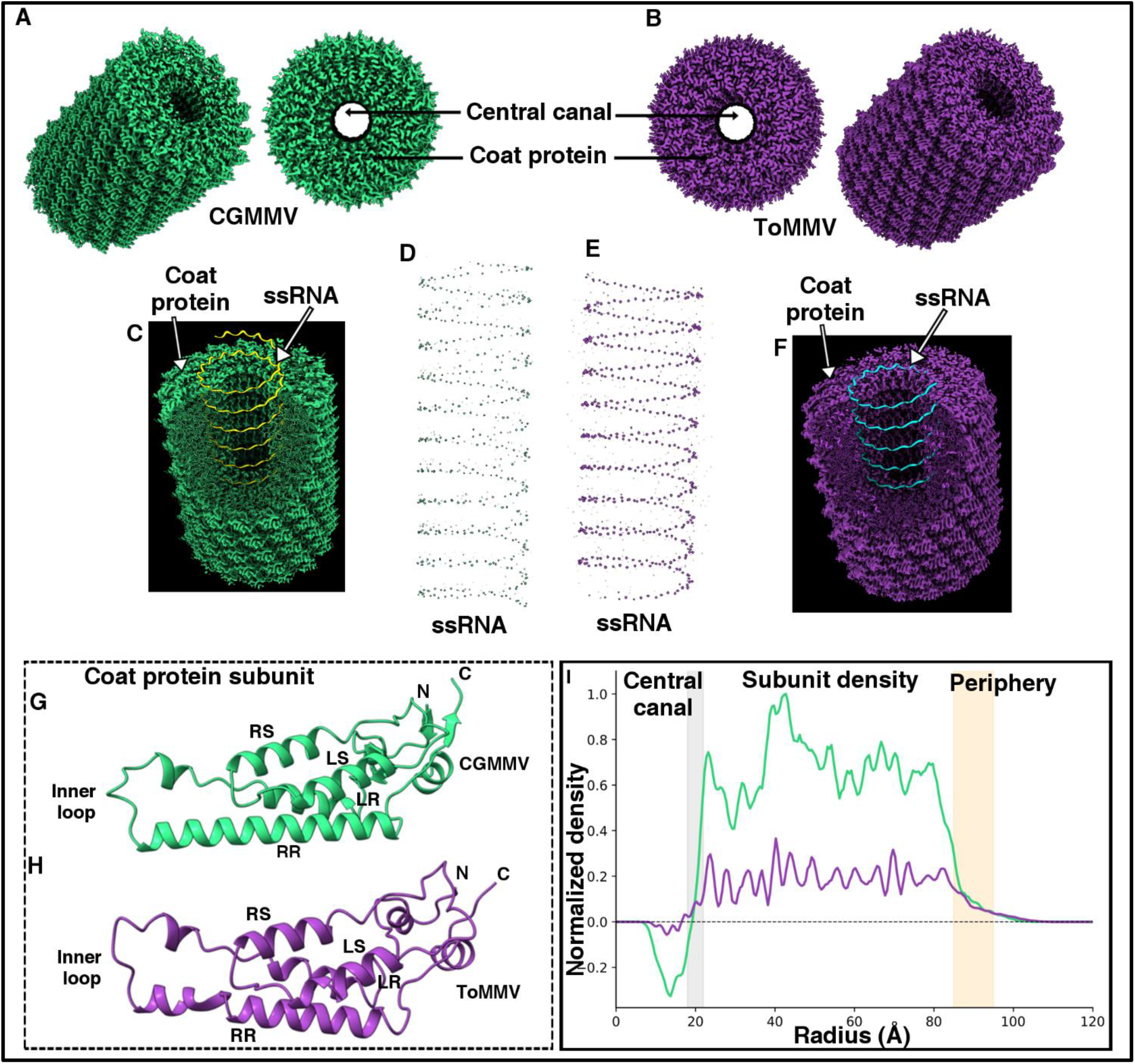
Overall architecture and atomic structure of CGMMV (Green) and ToMMV (Purple). Both virus maps show a helical rod-shaped structure with a central canal at the center (A, B), a characteristic feature of tobamoviruses. The maps are resampled for analysis and visualization purposes. Multiple copies of identical coat protein subunits are arranged in a helical pattern, with the helical ssRNA running throughout the whole virus. The internal architecture of CGMMV has been shown through a partially cut-open structure in C. Increasing the threshold value of the virion map reveals the RNA-associated densities, with prominent densities for the phosphate backbones (D, E). The internal architecture of ToMMV has been shown through a partially cut-open structure in F. Atomic structures of the coat proteins show the characteristic presence of a tobamoviral coat protein chain arrangement with four primary helices (G, H). The radial density map in I resolves the central canal and allows its diameter to be measured in the viruses; the boundary of the central canal is marked by the grey column. CGMMV (green curve) shows a more compact and ordered coat protein arrangement compared to ToMMV (purple curve). The outer capsid boundary is highlighted by the orange shaded column, where radial protein density decays to the solvent baseline. The threshold used for visualization for the CGMMV full map (A&C) is 1.55; for the ToMMV full map (B&F) is 0.55.

At higher map thresholds, density corresponding to the encapsulated RNA becomes distinct. RNA is single-stranded, right-handed, and follows the helical path of the entire virion (Fig. 2D, E). The coat protein subunit for both viruses is a single polypeptide chain forming four primary helices. In accordance with the general nomenclature system, the helices are named, from the N terminus, Left-slewed (LS), Right-slewed (RS), Right-radial (RR) and Left-radial (LR) (Fig. 2G, H). CGMMV showed three additional short helices and two pairs of antiparallel beta strands. We modelled the CGMMV coat protein with 155 residues (PDB-9VJD), while ToMMV coat protein model has 151 residues (PDB-9WQQ) (Fig. 2G, H). Atomic models were built and numbered based on the complete translated coat protein sequences. Note that canonical TMV structural models, including the ones used in this study for comparison, traditionally use a (-1) numbering shift relative to the primary sequence due to the post-translational removal of Met1.

Radial density profiles for CGMMV and ToMMV maps were generated following unified resampling, center alignment, baseline subtraction, and normalisation (Fig. 2I). Both viral reconstructions display a low-density central canal extending to a radius of approximately 20 Å. Beyond this boundary, CGMMV exhibits a sharp surge in radial density, maintaining a high-amplitude plateau across the core subunit region (r = 30-80 Å). In contrast, ToMMV displays a substantially lower mean normalised density (< 0.4) across the equivalent radial span. At the outer capsid periphery (r > 85 Å), CGMMV density declines steeply near r ≈ 95 Å and reaches bulk-solvent baseline by r ≈ 105 Å, whereas ToMMV tapers more gradually between r ≈ 80 Å and r ≈ 100 Å.

### 2. Geometric parameterization of periodicity and pitch-angle relationships across helical lattices of CGMMV and ToMMV

On the basis of the cryo-EM structure, the 3D density was further subjected to mathematical analysis of helical structures. A helix is either right-handed or left-handed, depending on the propagation along its axis of symmetry. The parametrized equation of a right-handed helix, propagating in the positive z-axis direction, is given as(30):

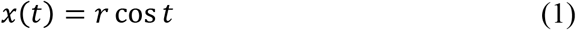

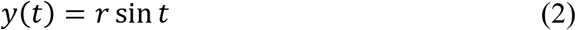

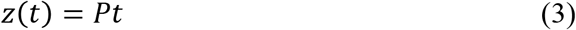

where t is the (time) parameter, r is the radius of the cylinder on which the helix lies, and P is the pitch or height per turn. As a trajectory in the (x,y,z)-space, it is expressed as (r cos t, r sin t, Pt). This helix propagates in the positive z-direction as time t increases while the spiral is moving counterclockwise. For a left-handed helix, we reverse the time parameter, i.e.,

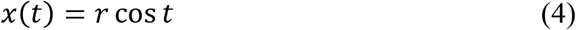

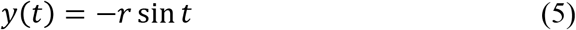

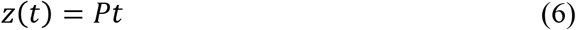

In our context, we consider right-handed helical structures for both viruses.

In the examples of the two viruses (CGMMV and ToMMV), one may imagine placing several units along the helix with parameters (r,P) such that condition 1, where any two adjacent units are equidistant; condition 2, the placement of any two units is equivalent, i.e., any unit can be obtained by appropriate rotational and translational symmetry applied to one base unit; and condition 3, after d turns, the (N+1)^th^ unit aligns vertically above the 1^st^ unit. (We assume that N and d have no common factors). Conditions 1 and 2 are natural biological and geometric assumptions which require no explanation. We shall explore the consequences and meaning of condition 3.

#### Periodicity relation

Condition 3 (Fig. 3A) is equivalent to

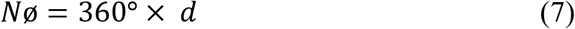

**Figure 3:**
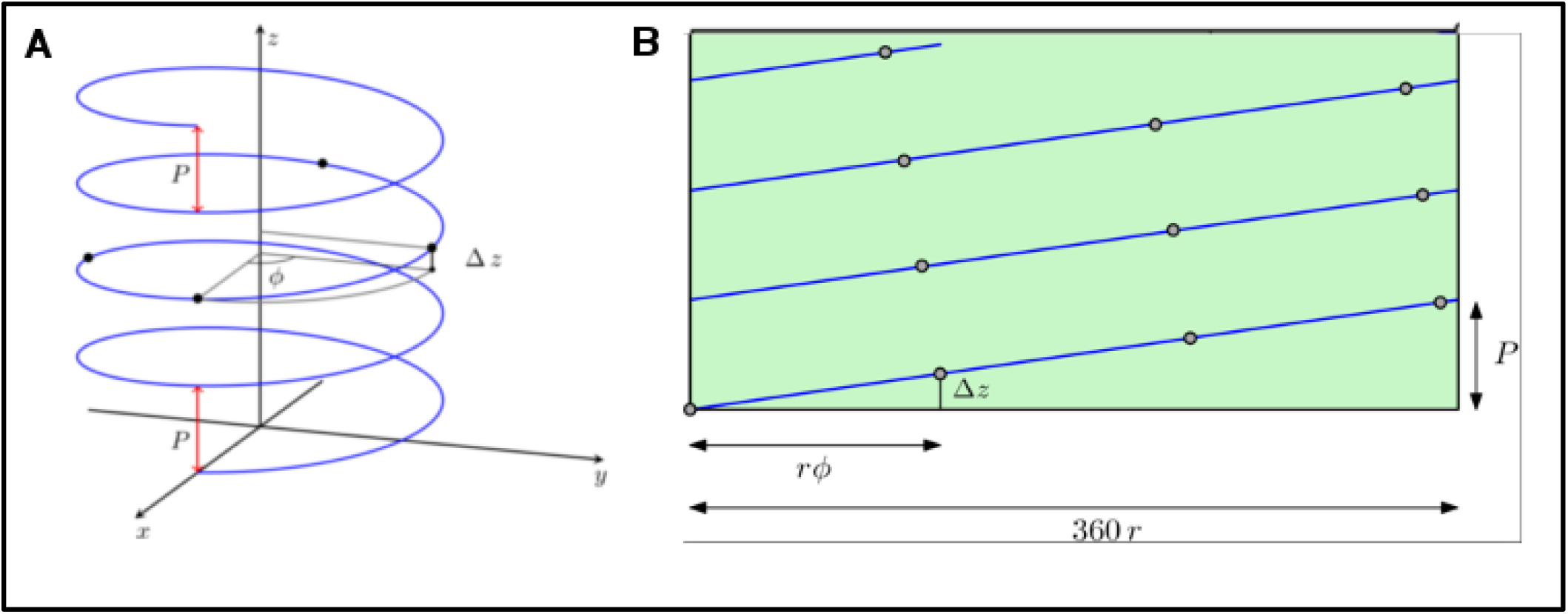
Two equivalent mathematical formulations (and visual depictions) of a helical structure. The marked points on the helix symbolize the nodes present in the helical virus structures being analyzed in this study. In A, the blue curve depicts the trajectory of a circular helix, with the z-axis as the helix axis. The pitch is denoted by P and is the vertical height difference under one full turn of the helix. The angular difference between two adjacent nodes (depicted by thick black dots) is denoted by ø. The height difference between two adjacent nodes is denoted by Δz. B shows the planar rectangular region obtained when the cylinder on which the helix lies is cut open. The helix is denoted by the blue line and the left vertical edge is understood to be identified with the right vertical edge height-wise. The grey bullets represent the nodes and P denotes the pitch of the helix. The radius of the cylinder (or helix) is denoted by r.

This can be seen by measuring the total turning as a particle moves up the helix and does d full turns. This contributes a total angle of 360° × d. The same motion can be alternatively seen as the sum of angle increments (ø) as one moves from one unit to the next. Each such move contributes ø degrees and there are N moves, making the total turn Nø. In particular, the periodicity relation implies that ø has to be a fraction of 360° for condition 3 to hold.

#### Pitch-angle relation

We have the following relation, due to similarity (Fig. 3B)

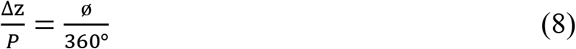

The formulas derived are exact. Using the periodicity relation defined above, minute structural differences can be detected among different helices.

### 1. Interactions and remodelling at the protein-protein interface with hydrophobic clustering and pH-dependent charge profiles

To compare the local contact geometries across virion interfaces, inter- and intrasubunit hydrogen bonding (H-bond) networks were mapped for both CGMMV and ToMMV (Table 1).

**Table 1:** Comparative analysis of inter- and intrasubunit H-bond interactions between CGMMV and ToMMV.

| **Interaction Type** | **Residue Pair** | **CGMMV Donor-Acceptor Distance (Å)** | **ToMMV Donor-Acceptor Distance (Å)** | **Significance / Note** |
|---|---|---|---|---|
| **Lateral (Inter)** | Arg93<sub>*i*</sub> - Glu96<sub>*i+1*</sub> | 3.092 | 2.684 | Conserved salt bridge; more compact in ToMMV |
| **Lateral (Inter)** | Arg123<sub>*i*</sub> – Asp89<sub>*i-1*</sub> | 3.038 | 2.677 / 3.383 | Bidentate in ToMMV |
| **Lateral (Inter)** | Arg114<sub>*i*</sub> - Asp116<sub>*i+1*</sub> | 2.770 | 3.014 | Lateral latch |
| **Axial (Inter)** | Arg93<sub>*i*</sub> - Asp117<sub>*i+16*</sub> | 2.909 | 3.061 | Part of trisubunit coordination |
| **Intrasubunit** | Arg62<sub>*i*</sub> - Glu146<sub>*i*</sub> | 3.160 / 3.320 | Weak electrostatic contact | Bidentate in CGMMV; weak in ToMMV |
| **Intrasubunit** | Arg46<sub>*i*</sub> - Asp43<sub>*i*</sub> | 3.093 | Absent | Right-slewed helix stabilization |

While ToMMV maintains localized high-strength interactions, such as the compact bidentate Asp89-Arg123 bond, CGMMV utilizes unique intrasubunit bracing to pre-stabilize the coat protein fold (Fig. 4A). Stability in the assembly is reinforced by a critical trisubunit coordination hub at Arg93, which simultaneously anchors lateral and axial neighbours to facilitate high-fidelity assembly (Fig. 4B). A detailed comparative list of H-bonds with the donor-hydrogen-acceptor angle is provided in Supplementary Table 1; map supports for the two major intersubunit H-bonds are provided in fig. S1a-d.

**Figure 4:**
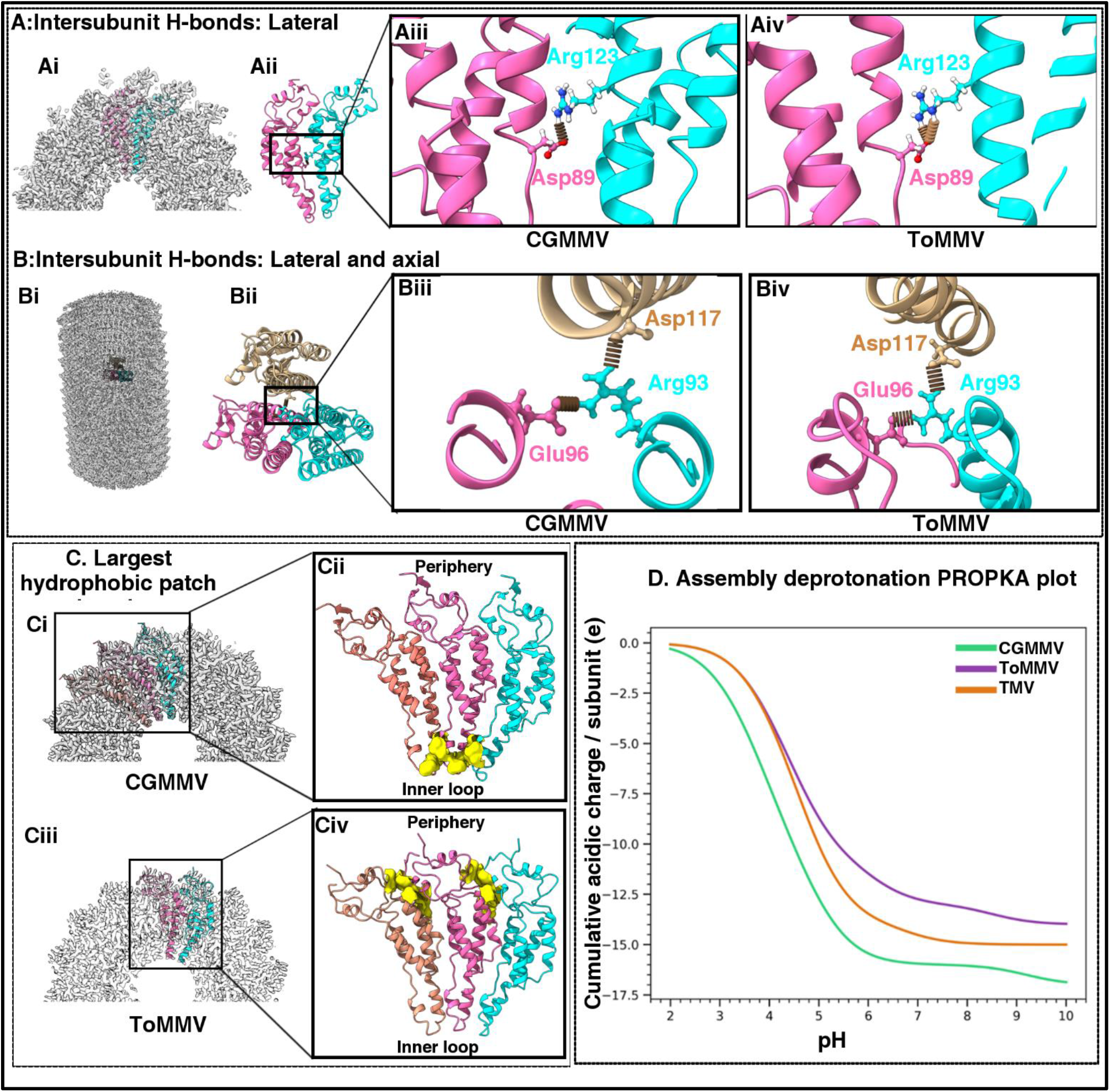
Inter-subunit interactions, surface hydrophobicity, and deprotonation behaviour in CGMMV and ToMMV viral assemblies. (Ai–iv) Lateral intersubunit hydrogen bonding between adjacent subunits (Chain *i* and Chain *i+1*) shown in overview (Ai, Aii) and close-ups for CGMMV (Aiii) and ToMMV (Aiv), highlighting key interactions involving Asp89 and Arg123. In ToMMV, this is a bidentate bond. (Bi-iv) Trisubunit axial and lateral connection networks formed by Chain *i*, Chain *i+1*, and Chain *i+16* in overview (Bi, Bii) and detailed close-ups for CGMMV (Biii) and ToMMV (Biv), showing interactions mediated by Glu96, Arg93, and Asp117 residues. (Ci-iv) Structural location of the largest contiguous hydrophobic patch (marked in yellow). In CGMMV (Ci, Cii), the patch is continuous and is located near the inner loop. In ToMMV (Ciii, Civ), the patch is located towards the periphery section and is intermittent. (D) PROPKA cumulative acidic-residue charge profiles per subunit (Q_acid_) plotted across a pH range of 2.0 to 10.0 for CGMMV (green), TMV (orange), and ToMMV (purple), demonstrating predominantly deprotonated carboxylate states at physiological pH that generate strong inter-subunit electrostatic repulsion (frustration) in CGMMV (≈-16e/subunit) compared to lower negative charge density in ToMMV (≈-12.5 to -13.5e/subunit).

The intersubunit hydrophobicity in canonical Tobamovirus TMV has been shown to influence structural stability, highlighting the role of hydrophobic interactions in capsid integrity (31–33). To evaluate the spatial distribution and extent of non-polar contact surfaces, hydrophobic patch mapping was performed using ΔSASA (Solvent Accessible Surface Area) analysis. Calculating ΔSASA upon assembly provides a direct, unbiased metric of hydrophobic burial that accounts for explicit atomic contact geometry without relying on empirical energy functions or arbitrary distance cutoffs. To prevent edge artifacts and avoid artificial over-counting of contacts from boundary chains, calculations were centered strictly on a central coat protein target subunit (viz. coat protein subunit “Chain c”) situated within a 9-mer local assembly matrix, ensuring that the central subunit experiences both axial and lateral interactions representative of the native helical environment (fig. S2). Surface hydrophobicity of this 9-mer assembly shows that CGMMV carries regular hydrophobic surface patches across the interfaces, whereas ToMMV has smaller scattered patches over the interfaces (fig. S3).

Across the 9-mer assembly (centered around Chain c), CGMMV contained 160 candidate buried hydrophobic residues, compared to 120 in ToMMV. Both viruses exhibited a total of 6 discrete hydrophobic patches (considering the minimum area of the patch >100 Å^2^) but differed in their spatial partition. CGMMV displays higher hydrophobic patch burial overall, with a mean patch buried area of 363.73 Å^2^ versus ToMMV (304.19 Å^2^), corresponding to a total of 36 residues buried in patches for CGMMV and 31 for ToMMV. The largest single hydrophobic patch (Patch 1) covered 558.12 Å^2^ (10 residues across Chains b, c, and d) in CGMMV and 535.48 Å^2^ (9 residues across Chains b and c) in ToMMV. The largest inter-subunit hydrophobic cluster in CGMMV (Patch 1) involves a concentrated proline-and valine-rich network (Val94, Val97, Val98, Pro100, and Pro103) spanning three adjacent subunits (b, c, and d). In contrast, the top two interface patches in ToMMV (Patches 1 and 2) are located primarily at two-chain interfaces (Chains b-c and c-d) involving aromatic and aliphatic residues (Phe11, Val12, Val22, Ile25, Phe68, Val130, Val134, and Phe139). The patch is located on either side of the subunit chains, but due to the spatial separation, it cannot form one large hydrophobic patch, unlike CGMMV that maintains an uninterrupted large patch at a trisubunit interface level. Hydrophobic interactions in CGMMV are organized into fewer but continuous regions rather than multiple discrete clusters. The largest hydrophobic patch in CGMMV is localized to the inner loop region of the coat protein, located in the central canal and exposed to solvent conditions, whereas in ToMMV it is situated towards the outer periphery (Fig. 4C). A detailed comparative ΔSASA analysis of the central chain c in a 9-mer assembly is provided in Supplementary Table 2a; details of the hydrophobic patches from both viruses, with participant chains and amino acids, patch type, and total ΔSASA for each patch are provided in Supplementary Table 2b, c.

In addition to hydrophobic packing, electrostatic interactions play a fundamental role in driving Tobamovirus assembly and stability, a hallmark feature well-characterised in TMV. To evaluate electrostatic stability during assembly in relation to structural accessibility, PROPKA-based deprotonation profiling was performed across a pH range of 2.0 to 10.0 for CGMMV, ToMMV, and TMV (Fig. 4D). Across the entire pH spectrum, CGMMV maintains a consistently more negative net charge than both ToMMV and TMV. Specifically, within the acidic-to-physiological transition (pH 3.0-7.0), all three assemblies undergo rapid deprotonation, but CGMMV exhibits the steepest accumulation of negative charge, reaching near-plateau negative charge values (around -16e per subunit) by pH 6.0-6.5. This sharp downward trajectory reflects its higher total number of acidic residues (17 Asp/Glu per subunit, compared to 15 in TMV and 14 in ToMMV) that readily deprotonate under mildly acidic conditions. In contrast, TMV and ToMMV reach less negative per subunit plateaus (approximately -14.5e and -12.5 to -13.5e at neutral pH, respectively), with ToMMV exhibiting the lowest acidic negative charge density across the physiological range. CGMMV shows significantly higher negative charge repulsion (electrostatic frustration) across physiological pH, requiring extensive hydrophobic burial or counterion screening to stabilize the helical assembly. Solvent accessibility and percentage burial (% Buried) analysis (Supplementary Table 3) reveals several Glu and Asp side chains that experience zero burial upon assembly, remaining fully solvent exposed with unperturbed p*Ka* values (≈ 3.5-4.5) that deprotonate readily under mildly acidic conditions. In CGMMV, a large area of solvent-exposed Glu155 (140.3 Å^2^) and Asp65 (112.9 Å^2^), alongside Asp43, Asp138, and Glu146, provides an excess of unshielded carboxyl groups compared to canonical carboxylates of TMV and ToMMV. This higher density of solvent-accessible acidic side chains largely accounts for the earlier and steeper plunge of the CGMMV curve, pulling its charge rapidly towards -16e per subunit by pH 6.0.

### 1. Synergistic stabilization of the ribonucleoprotein interface via electrostatic motifs at two different alpha-helix bundles

In the helically averaged cryo-EM maps of both viruses, the single-stranded RNA-associated density is resolved as a continuous, well-defined helical strand situated at a radial distance of approximately 40 Å from the central filament axis. The RNA density is clearly visible in the full reconstruction primarily at high threshold levels alongside regions of the coat protein possessing high negative charge; therefore, the phenix.map_box tool was utilized to extract and isolate the RNA density region for optimized visualization (fig. S4). The radial angle between the RNA and individual coat protein subunits is 72.2° in CGMMV and 77.9° in ToMMV (Fig. 5A, B). RNA-associated density strictly adheres to the viral helical symmetry, placed securely within the inter-subunit binding pocket between adjacent coat protein monomers, corresponding to three nucleotides per coat protein subunit. As helical averaging imposes strict structural symmetry along the length of the virion, this density represents a sequence-averaged ensemble of the genomic RNA. Individual nucleotide bases appear as consensus volumes instead of sequence-specific base conformations, and these RNA densities are modelled as 3-nucleotide bases (GAA) per coat protein subunit as per standard practice (Fig. 5C, D). The map clearly shows the spatial occurrence of the phosphodiester backbone for both viruses, residing within a specialized positively charged groove of the coat protein, i.e., the “RNA groove” coordinated by a pair of Arginine residues (Arg91 and Arg93), which is consistent with TMV (equivalent positions are Arg90 and Arg92) (Fig. 5E-H).

**Figure 5:**
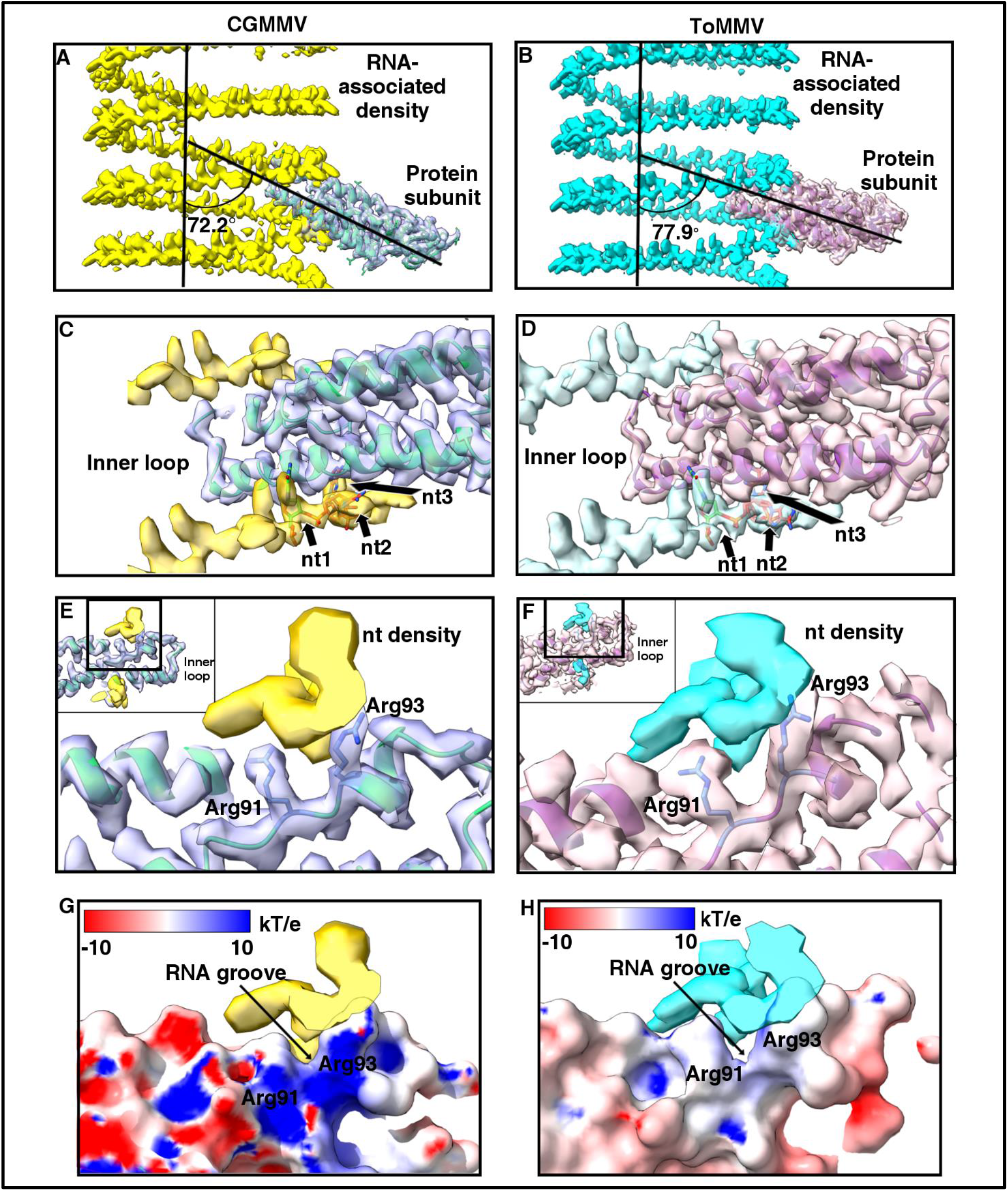
RNA-coat protein contact zones. The RNA-associated density was cut out from the whole map using phenix.map_box, with a clear phosphate backbone and averaged nucleotide densities. The coat proteins are placed at a wider angle with RNA in the case of ToMMV compared to CGMMV (A, B). 3 nucleotides of RNA are modelled following the density; helical averaging limits the bases to a sequence-averaged ensemble (C, D). RNA passes through a characteristic Arg91 and Arg93 pair in the lower helix region of the coat protein (E, F); insets show the location on the coat protein. The placement of this Arg pair forms a positively charged RNA groove in the area, as seen from the APBS surface electrostatic potential colouring of the coat protein subunits (G, H). Threshold values used for visualization for CGMMV panels are 1.37; for ToMMV panels are 0.237.

Comparative genomic parameter metrics for CGMMV and ToMMV are detailed in Table 2.

**Table 2:** Comparative genomic and capsid protein packing parameters in CGMMV and ToMMV.

| **Parameter** | **CGMMV** | **ToMMV** |
|---|---|---|
| **Genome size (nt)** | 6423 | 6399 |
| **Nucleotide per subunit** | 3 | 3 |
| **Subunits per 3 turns (approx..)** | 49 | 49 |
| **Total number of subunits per virion** | 2141 | 2133 |
| **Total number of turns per virion (approx.)** | 131.1 | 130.6 |

RNA-associated spatial coordination analysis from both viruses revealed that along the lower axial face of each coat protein subunit, the ssRNA strand runs directly adjacent to the N-terminal segment of the Left-Radial (LR) α-helix (residues 112–134; Fig. 6Ai, Ci). At this region, an electrostatically balanced pentapeptide patch (residues 113–117) was observed, comprising two consecutive basic residues followed by a neutral spacer and two acidic residues present in close proximity (Fig. 6Aii, Aiii, Cii, Ciii). In both viruses, positions 116 and 117 are occupied by consecutive Aspartate residues (Asp116 and Asp117); specifically, Asp117 serves as the axial partner in the trisubunit H-bonding hub described in Section 3 (Arg93*_i_*-Glu96i*_+1_*-Asp117*_i+16_*) (Fig. 6Av, Cv). Multiple sequence alignment across the genus *Tobamovirus* shows that this electrostatic patch shows strict conservation at positions 114 (Arg), 116 (Asp), and 117 (Asp), whereas position 113 demonstrates sequence variability (Fig. 6B). In TMV, Arg113 establishes a rigid, direct hydrogen bond with the RNA backbone at a distance of ≈ 3.4 Å(10, 14). In contrast, in our study in CGMMV, static modelled rotamer of Lys113 places its terminal ɛ-amino group at an average distance of 4.5-5 Å from the RNA phosphates. Despite this extended distance, a continuous bridging density envelope at 1.3σ is consistent with an RNA-protein interaction mediated by an ensemble of accessible lysine rotamers (Fig. 6Avi). This dynamic multi-conformer sampling mirrors the conformational disorder observed in real-space refinements of the TMV inner loop (10). In an Arg-Arg pair, both planar guanidinium groups form tight, bidentate interactions with the RNA backbone, whereas substituting Arg113 with Lys113 replaces one of these multivalent anchors with a flexible ammonium group carrying a single point charge. Consequently, replacing the rigid Arg-Arg pair with a Lys-Arg dyad weakens localized RNA coordination, shifting the viral genome anchoring mechanism from a tight electrostatic lock to a more diffuse interface.

**Figure 6:**
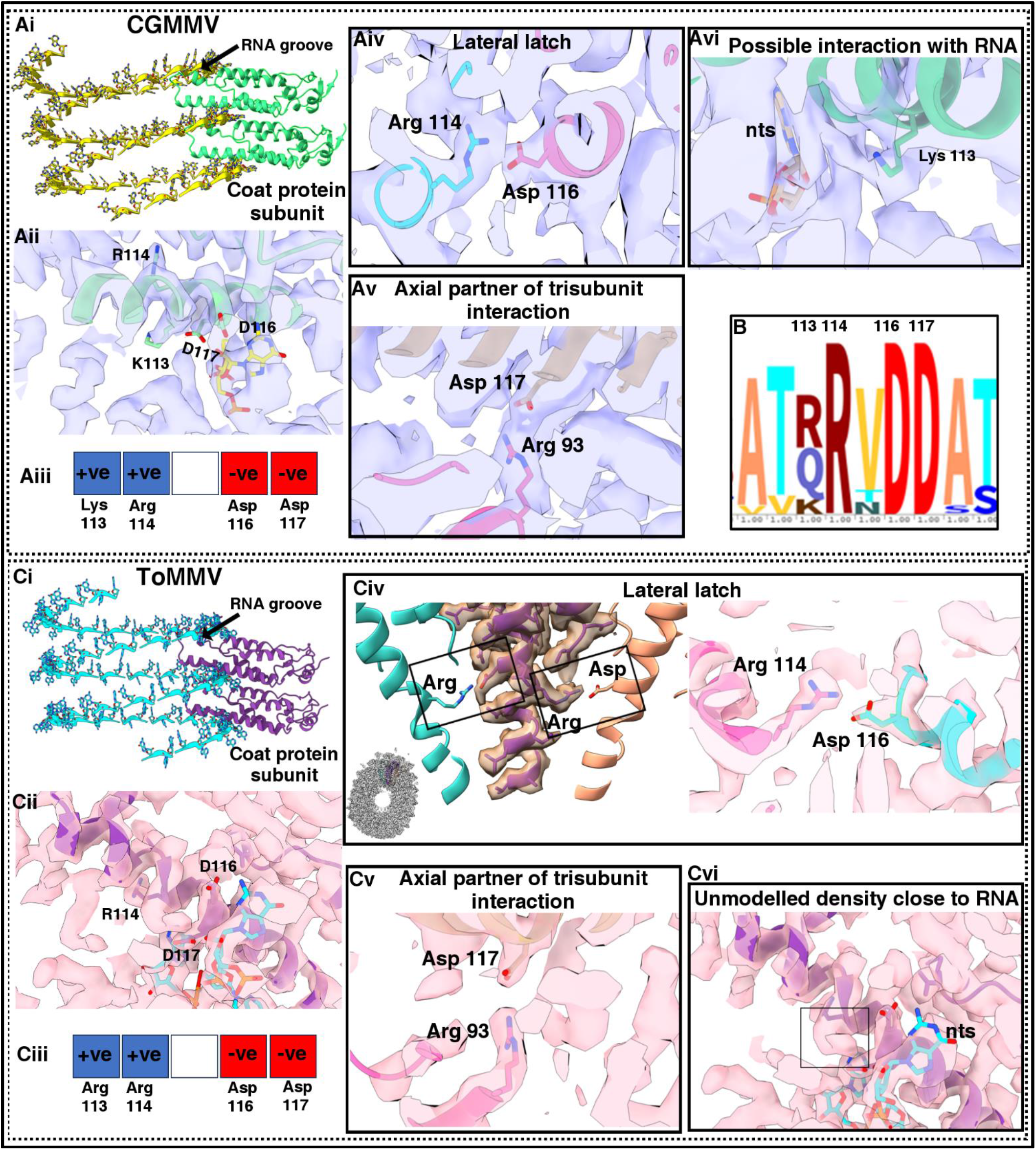
A conserved electrostatic pentapeptide motif at the lower left-radial helix of tobamovirus coat proteins. (Ai, Ci) Coat protein subunits intercalate between adjacent turns of the ssRNA helix, contacting the genome along both the upper and lower helical faces. (Aii–iii, Cii–iii) Whereas the upper RNA groove is coordinated by the conserved Arg91/Arg93 pair, the lower interface features a densely charged pentapeptide patch (residues 113–117) comprising Lys113 (CGMMV) or Arg113 (ToMMV) followed by invariant Arg114, Asp116, and Asp117. (B) CGMMV and ToMMV have Lys/Arg at the 113^th^ position, and the rest are made up of Arg114, Asp116, and Asp117. (Aiv) In CGMMV, Arg114i and Asp116i+1 form a direct inter-subunit lateral salt-bridge latch. (Av, Cv) In both viruses, the Asp117 side chain projects axially to coordinate Arg93 within the tri-subunit hydrogen-bonding hub. (Avi) In CGMMV, continuous bridging density connects Lys113 to the RNA phosphate backbone. (Civ) In ToMMV, Arg114 undergoes rotameric rearrangement to establish the lateral latch with Asp116i+1, accompanied by local unmodelled density near the RNA interface (Cvi) indicative of electrostatic side-chain plasticity. Threshold values for visualization are as follows Aii: 1.5; Aiv-vi: 0.5; Cii-vi:0.85.

In CGMMV, within this interface, Asp116 forms a direct salt-bridge partner with Arg114 from the laterally adjacent coat protein subunit, forming a lateral latch between neighbouring subunits (Arg114*_i_*-Asp116*_i+1_*) (Fig. 6Aiv). In the case of canonical TMV, residue 112 is occupied by an Arginine (10). Along with Arg90, Arg92, and Arg113, Arg112 completes an Arginine-rich basic pocket that neutralizes the negative charge of the RNA phosphate backbone(10, 14) [Residues following mature structure numbering, i.e., (-1) offset]. However, the spatial proximity of two consecutive basic residues with long side chains (Arg112 and Arg113) generates a localized electrostatic bottleneck. In our ToMMV cryo-EM map, this locus exhibits coalesced side-chain density in which discrete rotamers for Arg113 and Arg114 cannot be uniquely resolved [Residues following full sequence numbering]. Instead, we observed an alternately rotated Arg rotamer conformation (assigned as Arg114) which was locking the lateral latch with adjacent neighbour Asp116 (Arg114*_i_*-Asp116*_i+1_*) (Fig. 6Civ); and another undefined density that we could not model with any Arg rotamer (Fig. 6Cvi). The unmodelled density and alternative Arg rotamers point to conformational plasticity driven by local electrostatic repulsion between adjacent basic side chains. In this flexible state, both Arginine residues remain capable of establishing dynamic electrostatic contacts with the negatively charged viral RNA backbone. Rather than representing static disorder, this helically averaged density likely captures a dynamic conformational equilibrium driven by electrostatic repulsion between the adjacent guanidinium groups. This plasticity allows the Arg113-Arg114 dyad to alternate between securing the lateral intersubunit latch (Arg114*_i_*-Asp116*_i+1_*) and locking the RNA backbone from the lower radial helix face.

### 1. Evolutionary conservation and host-family associated partitioning of the 113-117 electrostatic motif

Evolutionary conservation profiling was performed using the empirical Bayesian Rate4Site algorithm implemented in ConSurf with canonical TMV single coat protein (PDB-4UDV) as the target sequence (Fig. 7A) (11). The analysis revealed that the surface-exposed 113-117 patch represents a evolutionarily constrained motif characterised by invariant electrostatic anchors. Specifically, positions Arg114, Asp116, and Asp117 achieved the maximum conservation score (Grade 9, normalised rate -1.299 to -1.303), designating them as strictly conserved, exposed functional residues across all analyzed viral taxa (Fig. 7B, Supplementary Table 4). In contrast, the internal positions 113 and 115 exhibit sequence plasticity where position 113 is classified as moderately conserved (Grade 7, score -0.652), partitioned primarily between basic and polar uncharged residues (45% Gln, 42% Arg, and 12% Lys), while position 115 serves as a largely conserved hydrophobic spacer (Grade 8, score -1.021; 81% Val, 9% Thr, 9% Asn) (Supplementary Table 4). Mapping these conservation tiers onto the TMV 4UDV monomeric structure illustrates that the invariant residues Arg114, Asp116, and Asp117 form an invariable and constrained local environment, whereas the chemical identity at positions 113 and 115 dictates local electrostatic potential and steric packing without perturbing the underlying four-helix bundle fold.

**Figure 7:**
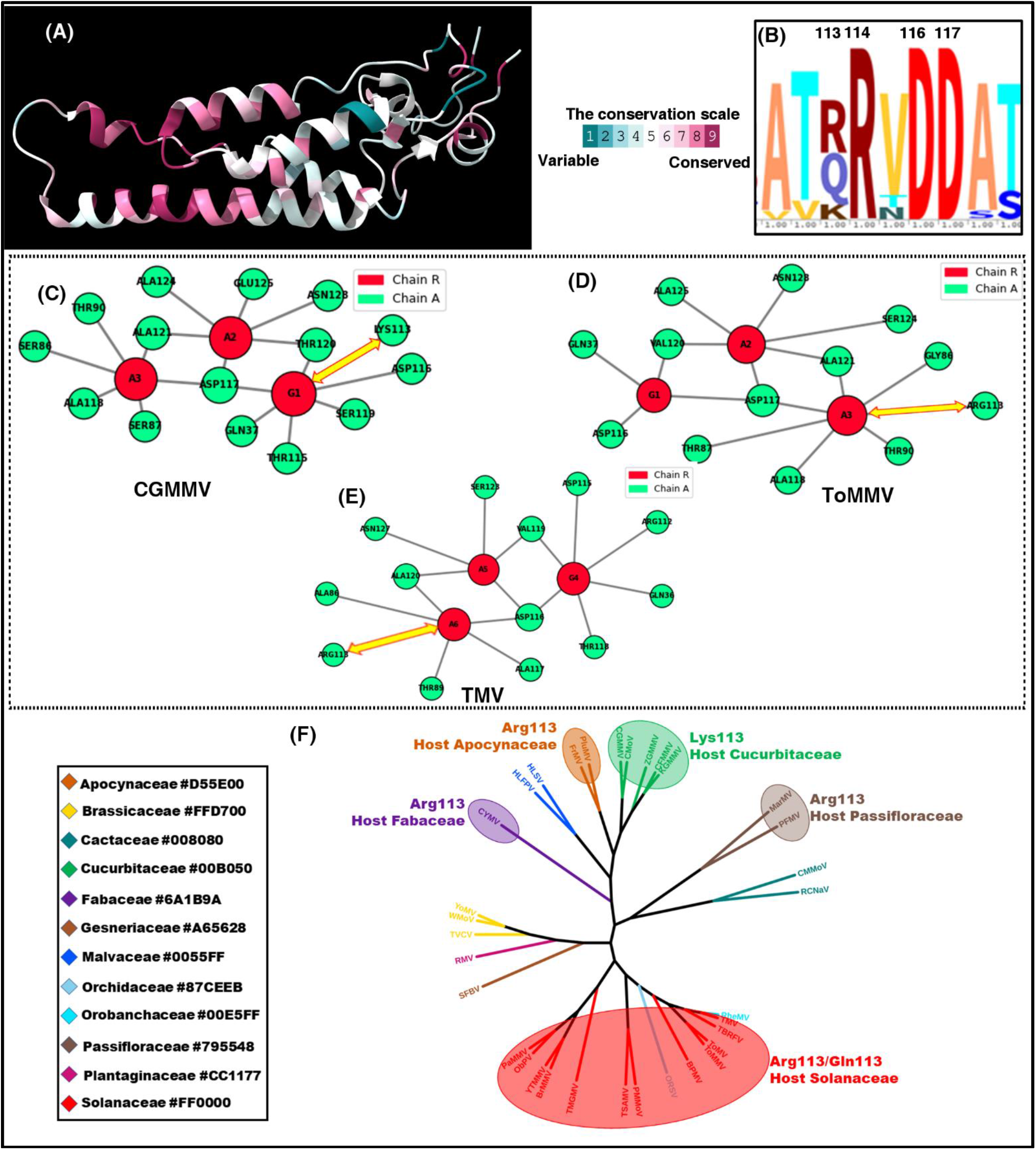
Conservation rate profiling and phylogenetic host-infection mapping of the coat protein 113–117 surface patch. (A) Ribbon structure of the coat protein monomer of TMV (PDB: 4UDV, Chain A) coloured by ConSurf evolutionary conservation scores from cyan (variable, Grade 1) to maroon (conserved, Grade 9), as shown in the inset. (B) Sequence-level alignment showing an invariant electrostatic patch formed by Arg114 (100% R), Asp116 (100% D), and Asp117 (100% D), flanked by the variable position 113. C, D and E show 2D interaction graphs depicting residue-level contacts between the viral coat protein (Chain A) and ribonucleotides (Chain R), highlighting coordinated RNA-binding pockets centered around nucleotides G1-A3 in CGMMV as shown in C, and G64-A66 in ToMMV as shown in D, and G4-A6 in TMV (E). Black outlines denote spatially distinct sub-interfaces, the key contact of Lys/Arg113-RNA is highlighted by the yellow double-headed arrow. F represents the Maximum-Likelihood tree (500 bootstrap replicates) showing phylogenetic segregation of residue 113/115 motifs by host family. Arg113 is widely distributed among Apocynaceae-, Fabaceae-, Passifloraceae-, and Solanaceae-infecting clades, whereas Lys113 is strictly restricted to a monophyletic Cucurbitaceae-infecting clade. Taxa representing tobamoviruses infecting the same host plant family are marked with the same colour. The colour scheme according to host family is provided in the inset.

To evaluate the interchain interactions between the viral coat protein and the ssRNA around this conserved patch, two-dimensional interface network graphs were generated using ProNA3D for CGMMV and ToMMV single coat protein (chain A) and GAA (chain R)(34). For comparison, TMV single coat protein was also taken into the study (PDB-4UDV). These network graphs mapped the contact landscapes into defined topological clusters, illustrating how individual nucleotides organize surrounding amino acid side chains across the macromolecular boundary. Within both viral assemblies, the protein segment spanning residues 113 to 117 forms an essential electrostatic and polar core directly engaging the ribonucleotide pocket (Fig. 7C-E). In CGMMV, this region is prominently featured at the G1 contact center, where Lys113, Thr115, and Asp116 coordinate the nucleic acid. Lys113 and Asp116, together with Thr120, delineate a distinct sub-interface cluster around G1 and adjacent nucleotide A2, providing positive charge complementarity and directed hydrogen-bonding capacity to stabilize the base and phosphate backbone. Additionally, Asp112 borders this segment and serves as a bridging node directly linking both A2 and G1 to the protein framework. In ToMMV, this binding segment exhibits clear functional homology paired with key residue substitutions. At position 113, basic coordination is maintained by Arg113, which forms a core sub-interface with Ala118 around nucleotide A66. Directly downstream in the loop, Asp116 is fully conserved and engages flanking nucleotide G64. While Thr115 in CGMMV is not retained as a direct contact in ToMMV, the adjacent acidic anchor shifts slightly in sequence register to Asp117, which acts as a major interaction hub coordinating G64, A65, and A66 simultaneously. Across both structures, the 113-117 motif operates as a conserved structural module combining a strongly basic anchor (Lys113 in CGMMV; Arg113 in ToMMV and TMV) and conserved carboxylate groups (Asp116 and Asp112/117) to lock the RNA backbone into the viral coat protein interface. In CGMMV, Lys113 participates in the coordination of guanine G1 alongside Thr120 and Asp116. In contrast, Arg113 of ToMMV forms a core sub-interface with Ala118 focused exclusively on A66, a contact mode mirrored in TMV where Arg113 similarly targets the 3′-terminal adenine A6. While all three viruses preserve the essential positive charge required for ribonucleic acid interaction, the Lys-to-Arg substitution pairs with an altered side-chain geometry and shifts the direct contact from a 5′-flanking guanine in CGMMV to a 3′-terminal adenine in ToMMV and TMV.

Projecting the discrete amino acid combinations at positions 113 and 115 onto the maximum likelihood phylogenetic tree demonstrated that surface charge remodelling correlates strictly with host family colonization and intra-host mobility (Fig. 7F). At the variable 113th position, the coat protein accommodates three distinct amino acid residues, Arg, Gln, and Lys, which correlate closely with the host plant family. The ancestral basic patch harbouring Arg113 (R113, V115 or R113, T115) is widely conserved across incongruent host families, including Apocynaceae (virus: PluMV, FrMV), Fabaceae (virus: CYMV), Passifloraceae (virus: MarMV, PFMV), and various solanaceous tobamoviruses. Interestingly, the Lys113 substitution is strictly restricted to a distinct, monophyletic Cucurbitaceae-infecting clade (including CGMMV, ZGMMV, CFMMV, with CMoV as the sole Gln113 exception), indicating that Lys113 serves as a defining synapomorphy of the major tobamoviruses infecting Cucurbitaceae hosts. Furthermore, the charge neutralization via a transition to Gln113 would reduce the local positive charge, which may limit non-specific electrostatic interactions along vascular cell walls.

### 1. Quantitative analysis of individual subunit stability and capsid packing efficiency

To evaluate the structural integrity and thermodynamic stability of the viral assemblies, a comparative analysis was performed among CGMMV, ToMMV, and TMV for an isolated coat protein subunit (Table 3).

**Table 3:** Various stability markers compared for the two experimental and the canonical Tobamovirus-TMV.

| **Stability markers** | **CGMMV** | **ToMMV** | **TMV** |
|---|---|---|---|
| **Volume (Å<sup>3</sup>)** | 20108.02 | 20808.04 | 20598.67 |
| **Solvent Accessible Surface Area (SASA) (Å<sup>2</sup>)** | 8803.78 | 9277.39 | 9223.71 |
| **Fractional hydrophobic SASA** | 0.36 | 0.28 | 0.26 |
| **Void volume (Å<sup>3</sup>)** | 5071.5 | 5606.2 | 5288.53 |
| **Packing Density** | 0.748 | 0.731 | 0.743 |
| **Compactness coefficient** | 2.46 | 2.53 | 2.53 |
| **Total polar surface area (Å<sup>2</sup>)** | 3552.34 | 4112.73 | 4128.4 |
| **Occurrence of Proline (%)** | 5.59 | 5.03 | 5.06 |
| **Alpha helix content (%)** | 55.27 | 52.83 | 56.96 |
| **Disulfide bridge formation** | Absent | Absent | Absent |

Quantitative comparison indicates that the isolated coat protein subunit of CGMMV possesses the smallest total volume (20,108.02 Å³) and the lowest solvent accessible surface area (SASA) (8,803.78 Å²), indicating reduced exposure of the subunit polypeptide chain to the surrounding aqueous environment. This compact structure is accompanied by a minimal void volume (5,071.5 Å³) and the highest packing density (0.748), signifying an efficient internal spatial arrangement of amino acid residues with minimal internal cavity space. A slight variation is observed in the compactness coefficient, where CGMMV displays a value of 2.46 compared to 2.53 for both ToMMV and TMV; however, this parameter reflects deviation from a perfect spherical geometry rather than internal packing efficiency, likely reflecting local variations in subunit elongation or curvature rather than structural instability. Furthermore, CGMMV exhibits the lowest total polar surface area (3,552.34 Å²), indicating greater burial of polar residues within the structural interior. Structural rigidity is supported by a higher proportion of proline residues (5.59%) in CGMMV relative to ToMMV (5.03%) and TMV (5.06%), which restricts main-chain conformational flexibility in CGMMV. Secondary structure composition remains predominantly helical across all three viruses, with TMV exhibiting the highest alpha-helix content (56.96%), followed by CGMMV (55.27%) and ToMMV (52.83%), while disulfide bonds are uniformly absent across all assemblies. Together, these structural metrics highlight the CGMMV isolated coat protein subunit as possessing a highly dense, rigid subunit architecture characterised by minimized solvent exposure, optimized internal spatial packing, and minimized cavity formation.

To evaluate the inter-subunit stability of the coat protein assemblies, the Buried Surface Area (BSA) was calculated for 3-turn structural assemblies of CGMMV and ToMMV, each comprising 49 coat protein subunits. BSA quantifies the surface area rendered inaccessible to solvent molecules upon macromolecular assembly, where higher values correspond to expanded inter-subunit contact interfaces, enhanced hydrophobic interactions, and favourable van der Waals contacts. For the complete 49-subunit assembly, CGMMV demonstrated a total BSA of 177,089.58 Å², compared to 175,689.94 Å² for ToMMV. On a per-subunit basis calculation, CGMMV has an average buried surface area of 3,614 Å², and ToMMV shows 3,585 Å², indicating that approximately 40-45% of the total monomer surface area becomes sequestered from the aqueous environment upon capsid formation. These substantial values reflect extensive inter-subunit interfaces characteristic of stable viral macromolecular architectures.

### 2. Structural plasticity at the inner loop revealed by comparative temperature-based molecular dynamics among three tobamoviruses

To evaluate the conformational stability and thermal response of individual coat protein subunits, molecular dynamics (MD) simulations were conducted for CGMMV, ToMMV, and TMV at 25 °C and 85 °C across a 200 ns trajectory. Analysis of backbone root-mean-square deviation (RMSD) demonstrates a clear temperature-dependent shift in conformational flexibility among the three tobamoviruses. At ambient temperature (25 °C) (Fig. 8A), CGMMV displays the lowest conformational drift, maintaining a tightly constrained backbone RMSD within 0.20-0.35 nm throughout the simulation run. ToMMV exhibits moderate baseline flexibility, converging to an RMSD plateau around 0.35-0.45 nm. TMV exhibits a slightly higher degree of initial structural deviation, maintaining an RMSD plateau between 0.4-0.5 nm after 100 ns, indicating higher intrinsic backbone flexibility in its monomeric state. Upon thermal exposure at 85 °C, distinct profiles of structural plasticity emerge (Fig. 8B). CGMMV retains substantial structural integrity, exhibiting only a slight elevation in backbone trajectory to an RMSD range of 0.30-0.45 nm with minor, constrained fluctuations. In contrast, both ToMMV and TMV exhibit heightened conformational dynamics at elevated temperature. ToMMV demonstrates a steady trajectory increase, reaching RMSD values of 0.50-0.60 nm after 100 ns. TMV displays the highest degree of conformational deviation under thermal stress, progressing toward an RMSD range of 0.55–0.65 nm toward the end of the 200 ns trajectory.

**Figure 8:**
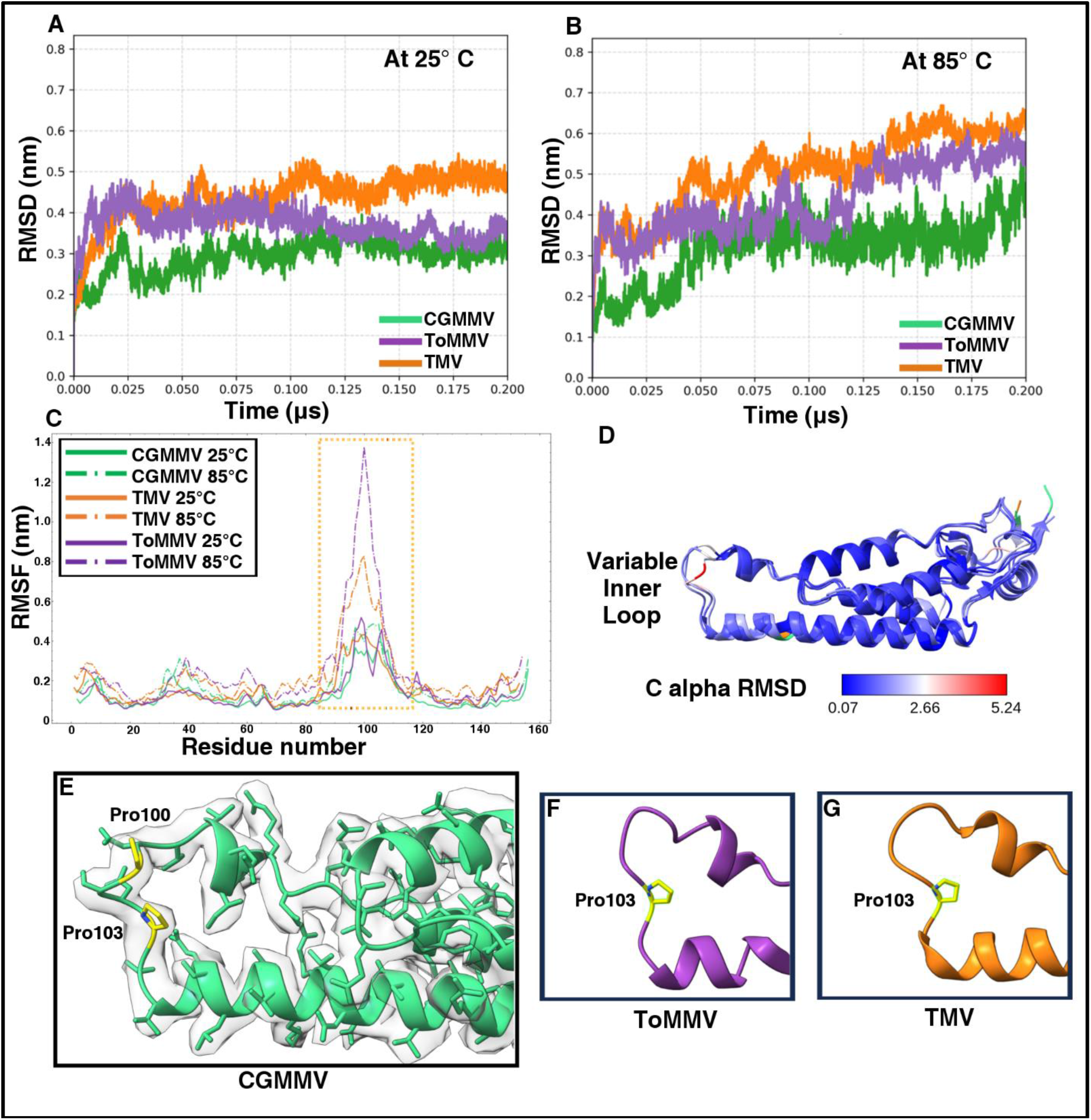
Temperature-variable comparative MD simulations of individual coat proteins demonstrate the stability of tobamoviruses as well as reveal the inner loop as the most dynamic region. Molecular Dynamics studies were performed on the reconstructed individual coat proteins from CGMMV and ToMMV, with TMV as the canonical representative of tobamoviruses. (A) The viruses show a stable trajectory at room temperature. (B) As temperature increases to a stress of 85°C, CGMMV shows the most stability (Green), while ToMMV maintains a median stability (Purple), and TMV shows very high susceptibility to increased temperature stress (Orange). (C) RMSF analysis revealed the inner loop to be the most dynamic region, with the CGMMV inner loop having the least motion. (D) C-alpha RMSD analysis of the three coat protein subunits showed that the primary structural difference comes from the inner loop region. (E) At the inner loop, CGMMV has two Prolines at the 100^th^ and 103^rd^ positions. Unlike CGMMV, both TMV and ToMMV carry a single Proline at the 103^rd^ position (F,G).

To pinpoint specific regions driving the overall conformational variations, residue-level Root Mean Square Fluctuation (RMSF) profiles were analyzed. Across all three viruses, the fundamental helical core regions remain highly rigid, displaying consistently low RMSF values (<0.2 nm). However, a pronounced peak in localized fluctuation is observed across residues ≈ 80-120 (boxed region), corresponding structurally to the variable inner loop, though the elevated-fluctuation window is broader than the loop itself and includes its flanking helices. At 25°C, CGMMV exhibits the lowest RMSF peak (≈ 0.35 nm) within this loop region, whereas ToMMV and TMV exhibit slightly higher fluctuations. Upon thermal perturbation at 85°C, the inner loop exhibits a marked increase in local mobility across all three coat proteins. This thermal response is most pronounced in ToMMV, where inner loop fluctuations surge to a maximum RMSF of approximately 1.35 nm, indicating severe local structural distortion. TMV also exhibits a notable elevation in loop mobility at 85°C, whereas CGMMV maintains the most constrained loop profile (≈ 0.5 nm) (Fig. 8C). The inner loop is a very vulnerable region of the coat protein, as it is situated in the central canal and remains fully solvent-exposed along the inner surface of the central channel. The inner loop is not protected by any secondary structure. C-alpha RMSD studies among the three tobamoviruses show that ToMMV bears more structural similarities with TMV, while CGMMV is divergent from both ToMMV and TMV (Fig. 8D). The main difference in the C-alpha RMSD of the coat protein subunit comes in the inner loop region.

The inner loop (residues 98–104), which projects into the lumen of the central canal, is composed predominantly of polar uncharged residues flanked in CGMMV by two prolines at positions 100 and 103 (Fig. 8E). Multiple sequence alignment (MSA) for tobamoviruses shows that the Pro103 residue is extensively present in many tobamoviruses, presenting itself as an almost conserved residue (fig. S5). In the case of both ToMMV and TMV, there is no Proline at the 100th position, increasing local conformational flexibility within the inner loop; both of these viruses carry Pro103 (Fig. 8F, G). This additional Pro100-induced type II turn renders the CGMMV inner loop markedly more compact than that of TMV or ToMMV (Fig. 8E-G). This alteration is possibly one of the main reasons for forming a more acute angle of attachment of the CGMMV coat protein to its ssRNA helix compared to that of ToMMV, leading to a more tightly packed CGMMV virion with a narrower central canal.

## Discussion

We report the cryo-EM structures of CGMMV and ToMMV at near-atomic resolutions, depicting the helical assembly of the coat protein subunits ensconcing the single-stranded RNA helix. The cryo-EM structure of CGMMV shows similar observable features obtained from previous Transmission Electron Microscopy (TEM) negative staining images and also to the previous 3.4 Å fiber diffraction structure (PDB-1CGM), but with better resolution(16, 35). The structure of ToMMV has a great resemblance to the recently published cryo-EM structure of TMV with 1.90 Å resolution (PDB-6SAE)(36). While both viruses maintain the characteristic helical arrangement of the genus *Tobamovirus*, structural comparison between CGMMV and ToMMV reveals notable local variations in coat protein subunit packing and interface geometry due to optimized amino acid coordination.

In canonical TMV, host-entry disassembly is governed by electrostatic repulsion across intersubunit carboxylate clusters, most notably the Glu95-Glu97 switch (corresponding to Glu96-Glu98 in full-length sequence numbering) near the inner loop, which destabilizes the capsid upon encountering elevated cytoplasmic pH and depleted Ca^2+^ concentrations (PDB ID: 6SAE) (36). In CGMMV, however, the substitution of Glu98 with a nonpolar valine (Val98) ablates this canonical Ca^2+^-binding site, necessitating an alternative stabilization and uncoating regime. Furthermore, the central canal is directly exposed to aqueous solvents, plant sap, and chemical disinfectants; therefore, the unstructured inner loop represents a primary vulnerability for thermal and hydrolytic degradation, as corroborated by our RMSF trajectories. In CGMMV, assembly of a contiguous, three-subunit hydrophobic belt (Val94, Val97, Val98, Pro100, Pro103) around the inner loop lining the central canal ensures a hydrophobic continuous zipper along the central canal, and the presence of the Pro100-Pro103 pair imposes a rigid type-II turn that locks the inner loop conformation and shields the hydrophobic core from water penetration (Fig 9, upper panel). By contrast, TMV-like tobamoviruses carry only a single proline (Pro103), leaving the inner loop conformationally relaxed and reliant on RNA binding for full ordering. Consequently, whereas thermal stress causes TMV coat proteins to dissociate into amorphous globular aggregates upon RNA degradation, heat treatment of CGMMV at 80°C causes the virions to fracture into short, rigid rod-like segments that retain their helical lattice (39). This exceptional hydrothermal and aqueous resilience explains how CGMMV persists in surface water and long distance agricultural irrigation networks (17, 37, 38; Fig. 9). Furthermore, within the host plant, a tightly packed and hydrophobically sealed capsid facilitates long-distance systemic transport through the horizontal cucurbit vascular network (40). Though the primary route of systemic movement of CGMMV is the phloem path, CGMMV frequently leaks out of the phloem into surrounding tissues and then reloads itself into the vascular tissue, with xylem being highly involved (41), facilitating systemic movement within the host vascular network. A highly surface-hydrophobic virion is an excellent advantage for causing this level of systemic infection.

**Figure 9:**
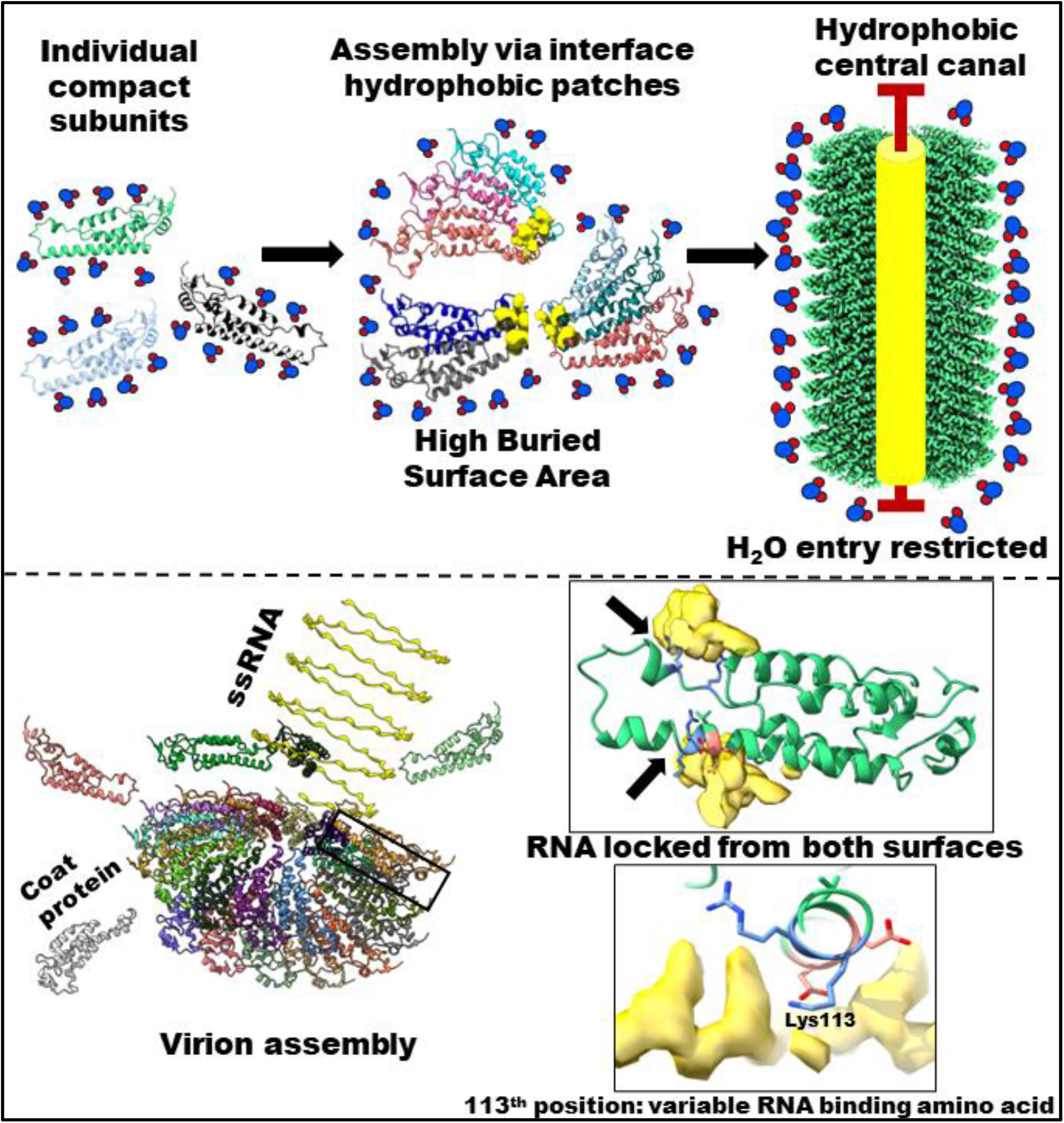
Model for helical virion assembly of CGMMV driven by a hydrophobic core and dual-surface ssRNA clamping. • **Upper panel:** Individual compact subunits initially exist in a solvated state with exposed surface area. Inter-subunit assembly aligns interface hydrophobic patches (yellow), yielding a high buried surface area that drives oligomerization. The resulting cylindrical complex forms a continuous, solvent-excluding longitudinal hydrophobic central canal that prevents water entry. • **Lower panel:** Helical single-stranded RNA (ssRNA) coordinates with the assembled scaffold during virion assembly. The RNA is locked and stabilized between adjacent subunit tiers through dual-surface contacts from both top and bottom surfaces. Inset illustrates the local coordination environment highlighting the residue at the 113th position acting as a variable RNA-binding amino acid among tobamoviruses and the position of conformational flexing of the RNA during RNA-protein binding.

Upon entering a host cell, TMV transitions from an inert virion into an active translation template via co-translational disassembly, wherein host ribosomes engage the 5′ end of the genomic RNA and progressively strip coat protein subunits from the ribonucleoprotein helix (42, 43). Because CGMMV lacks the canonical Glu96-Glu98 carboxylate trigger yet features tighter hydrophobic subunit packing, we hypothesize that its uncoating must be tuned at the protein-RNA interface. In CGMMV, the substitution of Arg113 with Lys113 accompanies a more acute subunit-to-RNA tilt angle (72.2° versus 77.9° in ToMMV), seating the genomic RNA deeper within the inter-subunit groove and likely shielding it from host ribonucleases during long-distance vascular transit. Simultaneously, replacing the bidentate guanidinium group of Arg113 with the smaller, monofunctional ɛ-amino group of Lys113 reduces local electrostatic binding affinity for the phosphodiester backbone. We propose that this Lys113 substitution establishes a mechanical trade-off: the genomic RNA is physically recessed for protection during extracellular and vascular transport, yet electrostatically poised for rapid mechanical extraction by translocating ribosomes during co-translational uncoating (Fig. 9, lower panel). The loose hold on RNA in Cucurbitaceae-infecting tobamoviruses such as CGMMV ensures that though the virus survives long-range transport, it undergoes coordinated capsid disassembly upon necessity. This model aligns with observations that intact CGMMV coat protein subunits persist after long-range phloem transport (40), reflecting their high intrinsic monomeric stability. Nevertheless, because our reconstructions were determined from virions purified in neutral phosphate buffer, future cryo-EM studies across systematically varied Ca^2+^ and pH conditions will be required to definitively exclude alternative cation dependent switches and biochemically validate this mechanical uncoating mechanism.

Taken together, our high-resolution structures and biophysical analyses define specific molecular targets for future mutational and functional studies across the genus *Tobamovirus*. The conserved 113–117 electrostatic motif on the left-radial helix, which integrates axial and lateral inter-subunit latches with direct RNA coordination, and the Pro100-Pro103 inner-loop motif represent prime candidates for site-directed mutagenesis aimed at attenuating CGMMV persistence in greenhouse and field environments. Extending structural characterization to additional cucurbit-infecting tobamoviruses, particularly Cucumber mottle virus (CMoV), an evolutionary outlier that carries Gln113 in place of Lys113, will further clarify how position 113 co-evolves with host range and vascular biology. Ultimately, the conjunction of structural and biophysical data in this article demonstrates that while the external morphology of tobamoviruses remains strictly conserved, fine-scale remodelling of internal hydrophobic and electrostatic networks enables radically different thermodynamic and mechanical strategies, offering design principles applicable to both antiviral crop protection and bio-inspired helical nanomaterials.

## Materials and Methods

### Virus isolation and purification

For this study, the CGMMV isolate designated as CGMMV-BgDel was used, which was originally obtained from a bottle gourd displaying mild mottling symptoms at the IARI vegetable field in New Delhi(44). Virions were partially purified following preestablished method by Gooding and Hebert, 1967(45). The ToMMV-Tss isolate was isolated and purified according to the methods of Mazumder et al. (2024)(29).

### Cryo-EM data collection

Cryo-EM grids were prepared using a Vitrobot Mark IV (Thermo Scientific, USA) operated at 18°C and 100% relative humidity. Before sample application, holey carbon Quantifoil grids (Au R 0.6/1 for CGMMV; Au R 1.2/1.3 for ToMMV) were rendered hydrophilic via glow discharge with a Quorum GloQube for 1 minute and 25 mA for CGMMV; and in a Quorum Q150TS Plus for 1 minute and 20 mA for ToMMV. The sample was applied to the grid, blotted to remove excess fluid (4 s blotting time for CGMMV; 3 s blotting time and a blot force of 1 for ToMMV), and vitrified by plunging into liquid ethane; storage was done in liquid nitrogen until further use. Data collection was performed at 300 kV with an amplitude contrast of 0.04 and a spherical aberration of 2.7 mm. For CGMMV, 1,132 gain-corrected movies were captured on a Titan Krios microscope equipped with a Falcon III direct detector, using an initial pixel size of 1.38 Å, 30 frames per movie, with an electron dose of 35.82 e^-^/Å^2^. For ToMMV, 4,884 gain-corrected movies were acquired on a ThermoFisher Scientific Krios G3i microscope equipped with a Gatan K3 detector, utilizing an initial pixel size of 1.1 Å, 38 frames per movie, and an electron dose of 39.3 e^-^/Å^2^. All automated data collection was driven by EPU software. Details of data collection, refinement, and validation statistics are provided in Supplementary Table 5.

### Cryo-EM data processing and helical reconstruction

All the movies were imported into RELION 3.0.8(46), followed by motion correction using RELION’s own implementation. The contrast transfer function (CTF) was estimated using CTFFIND4.1(47). Particles were picked as end-to-end helices in RELION(48) using a start-end coordinate convention. A total of 6582 filaments were traced for CGMMV, and 26020 filaments were traced for ToMMV (fig. S6 and S7). Overlapping helical segments were extracted in RELION using a tube diameter of 160 Å, a particle crop box size of 256 pixels, and a calculated helix step (inter-box distance) of 84 Å (corresponding to 60 unique asymmetrical units at a 1.4 Å helical rise), yielding an initial stack of 2,137,255 segments for CGMMV and 15,765,310 segments for ToMMV. These particles were then moved to CryoSparc 4.1.2(49) as a particle stack and were subjected to 2D classification. 3 rounds of 2D classification were performed for both viruses, and finally 44 classes for CGMMV and 25 classes for ToMMV were chosen for further processing, which showed defined helical features and had a resolution better than 15 Å, making the total final segment numbers 2,102,400 for CGMMV and 15,132,861 for ToMMV that were forwarded to helical refinement (fig. S8 and S9). Initial helical symmetry parameters were calculated from the layer-line projections of the best 2D classes using PyHI(50) (fig. S10 and S11). Helical refinement was performed in cryoSPARC utilizing these initial values as estimates. The local symmetry search was optimized across a grid space spanning a helical twist of 19°-25° and an axial rise of 1.26-1.54 Å. Final maps were post-processed using a solvent mask automatically optimized by the refinement workflow. The final global resolutions were determined to be 2.86 Å for CGMMV and 2.71 Å for ToMMV, respectively, based on the gold-standard Fourier Shell Correlation (FSC) threshold of 0.143 between independent half-maps (fig. S12 and S13). Local resolution estimation was completed using phenix.local_resolution (fig. S14-S17). The resulting cryo-EM maps and the atomic coordinates have been submitted to the EMDB and PDB databases subsequently.

### Model building and analysis

Initial model building was done using ModelAngelo (version 1.0.1)(51) with the build command using the flags -pf for protein and -rf for RNA. The model generated from ModelAngelo was produced from the whole map. Individual coat protein subunits from this model were checked manually to find the best-fitting subunit chain, which also best covers the sequence. Once that particular subunit chain was identified, that chain was extracted as a separate pdb file. This single-chain model, the protein sequence FASTA file, and the whole map were taken to Phenix and the map_box option was used to cut out the density of one coat protein subunit by using phenix.map_box, keeping the original absolute spatial coordinates intact (Keep origin = True). The sub-box boundaries were defined by generating an initial atomic mask using an atom mask radius of 2.0 Å, which was subsequently expanded using a selection radius of 3.0 Å to capture surrounding density relevant to the local atomic environment. To eliminate sharp truncation edge artifacts in Fourier space during downstream calculations, a 3.0 Å box cushion was added around the selected region in combination with soft mask extraction, ensuring a smooth density falloff at the box boundaries. Using this map boxed density, two iterative rounds of real-space refinement were executed in Phenix (phenix.real_space_refine) using default secondary structure, rotamer, and C-beta restraints, interwoven with manual rotamer adjustments, backbone fitting, and Ramachandran optimization in Coot (version 0.8.1)(52, 53). Model parameters were validated by MolProbity (54). Model was rigid-fitted with the whole map and boxed single-subunit density (fig. S18 and S19). Fitting of the side chains of the amino acids was checked in the pdb against the map (fig. S20 and S21). Density associated with the ssRNA was cut out using phenix.map_box with the whole map and RNA model generated from ModelAngelo with the same parameters as the boxed subunit density. The boxed RNA density is comparable to the RNA-associated density from the whole map and maintains a comparable local resolution as the map (fig. S4, S22 and S23). To understand RNA-coat protein interactions, a three-nucleotide GAA fragment was modelled against the RNA-associated helical averaged density (fig. S24). Viral assemblies of 3-turns with 49 subunits were built in ChimeraX using the helical rise and twist, with the density map as the guide(55). A pipeline for cryo-EM data processing, helical reconstruction, and atomic model refinement of CGMMV and ToMMV is provided in a schematic format in fig. S25. Full refinement and validation statistics are detailed in Supplementary Table 5.

### Radial density analysis

Radial density analysis was performed using a custom Python script (Supplementary Text S1). As the data collection parameters for the two viruses were different, in order to evaluate spatial mass distribution differences between the reconstructed capsids without introducing artifacts, maps were aligned and resampled to a common coordinate reference frame. Because the reconstructions possessed different initial voxel dimensions, magnifications, and box origins, the CGMMV density volume was resampled onto the spatial grid of the ToMMV map (1.1Å/voxel) using interpolation in UCSF ChimeraX. The central helical symmetry axes of both density maps were aligned, establishing a shared coordinate origin (x_0_, y_0_, z_0_) at the exact geometric center of both boxes. Three-dimensional cylindrical radial density profiles were extracted by integrating voxel density values across full concentric cylindrical shells centered along the aligned helical Z axis. The two-dimensional radial distance r*_i_* from the central Z-axis for any voxel located at (x*_i_*, y*_i_*, z*_i_*) was defined by:

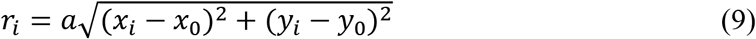

Where *a* represents the isotropic voxel spacing (1.1 Å/voxel).

Concentric shells were evaluated at a radial step size of 0.5 Å, oversampling relative to the 1.1 Å voxel size to avoid radial binning artifacts and capture fine structural features without over-smoothing or introducing discrete grid noise. Baseline correction was applied by calculating the mean density within the exterior bulk solvent shell (105 Å ≤ r ≤ 115 Å) where no macromolecular protein mass is present and subtracting this background offset from the complete radial profile. To facilitate direct, proportional comparison of density amplitudes across reconstructions while preventing scaling artifacts from surrounding volume padding, profiles were normalised relative to the maximum baseline-corrected density peak of the CGMMV reference map. This allows direct spatial comparison of shell features, peak locations, and boundary attenuation between the two viral capsids. The final baseline-corrected radial density profiles were plotted against the radius in Å using Matplotlib(56).

### Hydrogen bonds

To observe the H-bonds, similarly, a 3-turn working unit with 49 subunits of the virus was made as described above. The H-bonds were calculated after adding hydrogen (AddH) using the structural analysis option in ChimeraX with the following parameters: Distance tolerance 0.4 Å, Angle tolerance 30°.

### Assembly deprotonation profiles and Subunit surface electrostatics

To evaluate and compare the pH-dependent carboxylate deprotonation profiles of the three-turn, 49-subunit virus assemblies (CGMMV, ToMMV, and TMV), subunit-normalised titration curves mapping the cumulative acidic-residue charge per subunit (Q_acid_) against pH were generated. Structural p*K*_a_ data for the acidic residues (Asp and Glu) within each virus assembly were modelled within the context of the three-turn assembly using PROPKA(57) to capture intersubunit desolvation and electrostatic coupling. Using a custom Python script (Supplementary Text S2), the ionization state and fractional negative charge of each individual acidic residue (*i*) as a function of pH were determined via the algebraic rearrangement of the Henderson-Hasselbalch equation,

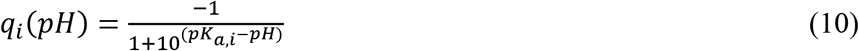

Rather than plotting independent single-residue titration curves, the cumulative acidic-residue charge per subunit (Q_acid_) across a pH range of 2.0 to 10.0 was obtained by summing the fractional negative charges of all Asp and Glu residues per subunit (17 in CGMMV, 15 in TMV, and 14 in ToMMV) at each pH step. All deprotonation mathematical calculations and data visualizations were executed in Python using NumPy for array operations and Matplotlib(58). Complementing the assembly-wide titration calculations, the distribution of surface electrostatics for the individual coat protein subunits was modelled using the Adaptive Poisson-Boltzmann Solver (APBS)(59).

### Solvent Accessible Surface Area (SASA)

SASA measures the surface area of a biomolecule that comes in contact with the solvent. Lee and Richards (1971)(60) defined the solvent molecule as a spherical probe, typically 1.4 Å in radius to represent water. The probe rolled over the surface of a larger molecule. Tracing the path of the center of the probe provides the value of SASA.

In this work, SASA was calculated using the program Freesasa (version 2.1.2)(61). The surface is approximated as a set of slices, which is a classical estimate coined by Lee and Richards (1971)(60). In this context, SASA quantifies static structural exposure and interface shielding; however, it does not directly capture energetic contribution/affinity, electrostatic complementarity, or dynamic, time-dependent surface fluctuations.

### Hydrophobic patch analysis using SASA

To characterise spatial clusters of hydrophobic residues involved in intersubunit interfaces, a nonameric (9-mer) protein assembly consisting of a 3×3 arrangement of coat protein subunits across three adjacent helical turns was chosen as the minimal structural model scale for this characterization, where the target subunit is placed at the center. The 9-mer configuration represents the minimal symmetric unit required to simultaneously model both the horizontal (lateral) and vertical (axial) interfaces of a central coat protein subunit. Positioning the target subunit at the center of a 3×3 grid provides a physiologically relevant and complete local coordination sphere while eliminating boundary artifacts and avoiding the computational burden of full viral assemblies (fig. S2).

Within this 9-mer assembly, the change in solvent-accessible surface area (ΔSASA) upon oligomerization was calculated. Measuring ΔSASA upon molecular assembly directly quantifies how much nonpolar surface becomes shielded from water based purely on exact atomic geometry. This metric assesses hydrophobic burial objectively through physical contact surfaces, avoiding the need for estimated energy equations or arbitrary distance thresholds. Freesasa (version 2.1.2) outputs(61) for both monomeric and nonameric states were analyzed by a custom Python script (Supplementary Text S3) utilizing the SciPy and NumPy libraries (58, 62).

ΔSASA is calculated as the difference between the monomeric and nonameric assembly states for the target subunit.

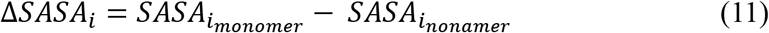

Where SASA*_i_* represents the SASA of the center subunit (*i*th subunit) in monomeric or nonameric form.

Hydrophobic residues (defined as Ala, Val, Leu, Ile, Met, Phe, Trp, Tyr, and Pro) exhibiting a ΔSASA greater than 25.0 Å^2^ were isolated as core interface participants. Spatial clustering of these target residues was subsequently conducted using their C alpha spatial coordinates gathered from the local assembly PDB file. A Euclidean distance threshold (d≤7.0 Å) was applied and contiguous networks containing a minimum of two residues were isolated via a depth-first search graph traversal strategy. Clustered spatial networks were classified into interface patches if they spanned across multiple interacting coat protein subunit chains or single-chain patches if confined to an individual monomeric subunit.

The change in SASA upon assembly (ΔSASA_i_) for the acidic residues (Asp and Glu) was calculated similarly as the difference between the monomeric and assembled states. The percentage burial (% Buried) upon assembly was subsequently determined as the percentage of the monomer’s accessible area buried at the interface:

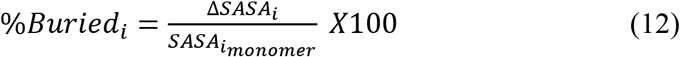

### Phylogenetic analysis

Coat protein sequences of all the tobamoviruses were obtained using NCBI. The sequences were first aligned using ClustalW embedded within MEGAX software(63, 64). A comprehensive Gonnet weight matrix was used with a Gap Opening Penalty of 10.00 and Gap Extension Penalties of 0.10 for pairwise and 0.20 for multiple alignments. These parameters allowed the algorithm to lock the matching parts of the conserved secondary structures in place while letting the more flexible loop regions stretch or gap as needed. The negative matrix option was turned off to prevent over-penalization and premature termination of alignments for the significant sequence diversity. With the aligned sequences, the Maximum Likelihood method was chosen to build the trees (ML trees) with 500 bootstrap replicates to minimize errors using MEGAX(64). Trees are represented using the TreeViewer software(65). The list of all the tobamoviruses from ICTV taken for this phylogenetic study is provided in Supplementary Table 6 with accession number, isolate, abbreviations, host family, and 113^th^ place amino acid composition. The final ML phylogenetic tree with support values is provided in Newick format in Supplementary Text S4.

### Evolutionary conservation analysis

Position-specific evolutionary conservation across the viral coat protein was determined using the ConSurf web server(66). To prevent sampling bias and reflect accurate phylogenetic relationships, the analysis was executed using user-provided datasets comprising the reference atomic model, multiple sequence alignment (MSA), and phylogenetic tree. The experimental high-resolution structure of the coat protein monomer (PDB ID: 4UDV, Chain A) was utilized as the 3D structural anchor. The coat protein MSA generated via ClustalW was supplied as a custom input. Evolutionary rates per amino acid position were calculated using the empirical Bayesian inference method implemented within the Rate4Site algorithm(67). Amino acid substitutions were evaluated under the Best-fit empirical evolutionary substitution model. Continuous evolutionary rates were normalised and partitioned into discrete conservation grades ranging from 1 (most variable) through 5 (intermediate) to 9 (strictly conserved). In addition, solvent accessibility and residue status, designated as buried structural or exposed functional, were predicted using the neural network algorithm embedded within the ConSurf pipeline. The resulting normalised conservation scores and confidence intervals were projected onto the 3D atomic coordinates of the reference structure (PDB: 4UDV, Chain A).

### Protein–RNA interface analysis using ProNA3D

To characterise the interaction between the viral coat protein (chain A) and the RNA molecule (chain R), the structural coordinates from PDB files of both viruses (with GAA trinucleotide as RNA) were imported into UCSF ChimeraX and evaluated using the ProNA3D extension(34). Pairwise interface detection and sub-interface clustering between chain A and chain R were conducted using ProNA3D’s default spatial parameters, applying a heavy-atom contact distance threshold of 5.0 Å and an interface clustering cutoff of 5.0 Å to calculate contacting residue-nucleotide pairs and group them into distinct interaction hubs. 2D interface network graphs were subsequently generated directly via the plugin to map residue-level connectivity, delineate spatially separated sub-interfaces, and identify critical contact partners across the protein-RNA boundary.

### Buried Surface Area (BSA)

Buried Surface Area is quantified as the previously accessible surface area that becomes inaccessible to the solvent upon binding of two molecules(68). BSA is used for measuring the geometric interface in bound conformations of a protein complex. Unlike closed point-group oligomers, helical assemblies feature extensive overlapping interfaces from both lateral (*i* + 1) and axial (*i* +16 or *i* +17) subunit interactions. To account for boundary effects from solvent-exposed terminal subunits, calculations were performed using an intact helical segment, making BSA (Å^2^) for a protein complex with repetitive subunits expressed by:

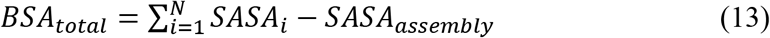

Where N is the total number of repetitive subunits in the assembly, and SASA*_i_* is the surface area of subunit *i* calculated in isolation (unbound state). We have used a 3-turn 49-subunit assembly to calculate the parameters, which were calculated using Freesasa (version 2.1.2)(61).

### Compactness

Compactness quantifies how closely the shape of a protein is compared to that of an ideal sphere, which is the most compact geometric form(69).

Compactness is defined as:

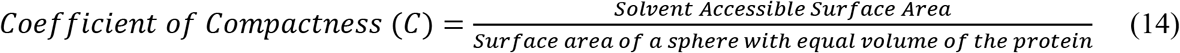

While compactness effectively measures overall geometric elongation and surface-to-volume efficiency, it does not assess internal packing quality, atomic density, or the presence of internal cavities.

### Void volume and packing density

Void volume (V_void_) of a protein represents the internal cavities and pockets caused by imperfect packing of amino acid side chains. These voids are crucial for protein flexibility, ligand binding, and the protein’s response to high pressure(70).

V_void_ (Å^3^) is defined as:

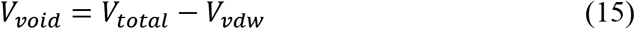

Where V_total_ is the total volume, and V_vdw_ is the Van der Waals volume. However, V_void_ measures purely geometric empty space and does not account for the hydration state, chemical nature, or energetic stability of these pockets.

Packing density (PD) measures how tightly the atoms of a protein are packed together(70). PD is defined as:

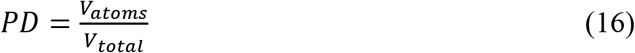

Where V_atoms_ is the volume the atoms occupy, and V_total_ is the total volume.

In this work, Vvoid and PD were calculated using the ProteinVolume 1.3 server(70). Although PD provides an overall index of packing efficiency, it represents a global average and does not detect localized packing defects, nor does it directly measure thermodynamic stability or dynamic conformational entropy.

### Molecular dynamics studies

The modelled PDB files and PDB-4UDV for TMV(11) were subjected to all-atom molecular dynamics (MD) simulations using GROMACS 2023.1 with the OPLS-AA/L force field(71, 72). To evaluate the intrinsic structural flexibility and thermal stability of individual coat protein subunits, simulations were conducted on isolated monomeric models. Each protein monomer was centered in a cubic box with a minimum distance of 1.2 nm between the protein and the box edges. Systems were solvated with explicit TIP3P water molecules, and background ionic strength (0.15 M) was established by adding Na^+^ and Cl^-^ions as counterions using the gmx_genion module.

Energy minimization was performed using the Steepest Descent algorithm until the maximum force fell below 1000 kJ mol^-1^ nm^-1^. Short-range non-bonded electrostatic and van der Waals interactions were calculated using a cutoff distance of 1.0 nm. Long-range electrostatics were treated using the Particle Mesh Ewald (PME) method with a grid spacing of 0.12 nm. All covalent bonds involving hydrogen atoms were constrained using the LINCS algorithm, enabling an integration time step of 2.0 fs.

Temperature equilibration was performed in an NVT ensemble for 100 ps using the V-rescale thermostat (τ_t_ = 0.1 ps). Subsequent pressure equilibration was conducted in an NPT ensemble for 100 ps at 1.0 bar using the isotropic Parrinello-Rahman barostat (τ_p_ = 2.0 ps, compressibility = 4.5 × 10^-5^ bar^-1^). During both equilibration phases, harmonic position restraints (*k* = 1000 kJ mol^-1^ nm^-2^) were applied to all protein heavy atoms. Production MD runs were carried out for 200 ns under NPT conditions across two target temperatures: 298.15 K (25°C) and 358.15 K (85°C). To ensure statistical reliability, all production trajectories were performed in triplicate (n = 3) for each virus and temperature condition.

### Root Mean Square Deviation and Root Mean Square Fluctuations

Root Mean Square Deviation (RMSD) is a numerical measure to quantify the similarity between two protein structures. It calculates the average distance between equivalent atoms between two superimposed molecules and is calculated for the protein backbone to check for the overall fold(73). To quantify the time-averaged local flexibility and structural mobility of individual amino acid residues across the isolated coat protein subunits during the molecular dynamics simulations, Root-Mean-Square Fluctuation (RMSF) profiles were calculated.

**Note:** Salient features of the entire study have been visually represented as a movie file named “Movie S1.mp4”.

## Acknowledgments

We acknowledge Dr. Partha Ray Choudhury (Principal Scientist, Crop Science Division, ICAR Head Quarters, New Delhi), Dr. Santanu Sankar Aich (Senior Scientist, Field Crop Research Station, Burdwan, WB) and Dr. Anita Roy Aich (Associate Professor, BCKV, Mohanpur, Nadia, WB), for helping PPD find the sources of the plant virus isolates. We acknowledge Prof. Jose Maria Carazo (Professor, Spanish National Center for Biotechnology, CNB-CSIC, Spain) and his research team, especially Mikel Iceta Tena, for their help in configuring the architecture of the computer workstations used in this study, and, for the same reason, we acknowledge Dr. Somnath Dutta (Associate Professor, IISc, Bengaluru), and Dr. Vinothkumar Kutti Ragunath (Scientist, NCBS, Bengaluru) as well. We acknowledge Dr. Vinothkumar Kutti Ragunath and the National Electron Cryo-Microscopy Facility at the Bangalore Life Science Cluster (inStem, NCBS, and C-CAMP), supported by grants from the Department of Biotechnology (DBT), Government of India, for collecting the CGMMV cryo-EM micrographs. We acknowledge Prof. Manidipa Banerjee (KSBS, IIT Delhi), Dr. Vijay Singh, and the technical support provided by the Cryo-EM facility at the Sophisticated Analytical & Technical Help Institute (SATHI) and the Central Research Facility (CRF) of the Indian Institute of Technology Delhi (IIT Delhi) for ToMMV cryo-EM data collection using the 300 kV Titan Krios G3i Cryo-TEM, established via the SATHI initiative of the Department of Science and Technology (DST), Govt. of India. We acknowledge Prof. Subhash Chandra Yadav and Dr. Shikha Chaudhary of the Electron Microscope Facility, Department of Anatomy, AIIMS, New Delhi, for help in the initial testing of some of our samples. We thank UD lab students in the Biophysics Department, AIIMS, New Delhi, for storing ToMMV in their facility. PPD is grateful to his former Professor, Dr. R. Usha of MKU, Madurai, for introducing him to plant virus research. AC received a PhD fellowship from the Department of Science and Technology, Govt. of India. AV is provided with a PhD fellowship from IISER Kolkata.

## Funding

All the cryo-EM image processing, analysis, and MD simulations work has been done using both the DBS computational facility funded by IISER Kolkata and the PPD Lab workstation obtained through a grant from the Ministry of Education, Govt. of India (previously known as MHRD-STARS), Grant number: STARS/APR2019/BS/581/FS.

## Author contributions

AC: Overall execution of the project including Cryo-EM reconstructions, model building, MD simulations, various computations, data analysis, map and model deposition, figure generation and manuscript writing, AV: Various software installation, executions and running of those softwares, MD simulations, managing of the workstations and cryo-EM image processing, model building and data analysis; AAKJ: Isolation of CGMMV; PM: Isolation of ToMMV; SS: cryo-EM data collection of ToMMV; SB: Mathematical topology study; UD: Advice on the optimization of the workstations through remote installations of appropriate OS, suggestions on various software installation, advice on executions and running of those software for cryo-EM image processing and computations, various problem solving and manuscript editing. AR and BM supervised the isolation of CGMMV and ToMMV and provided the isolated virus samples to PPD; PPD: Inception and concept of the project, planning and roadmap, finding the resource persons to obtain the virus samples, obtaining all other resources, including financial and instrumental resources, data collection, building up the entire team, development of the entire workflow, execution, data and figure analysis, problem solving, management and overall supervision of the project, figure and manuscript editing.

## Competing interests

The authors report no competing interests.

## Data, code, and materials availability

CGMMV density map is available in EMD-60696; the coat protein atomic model is available in PDB-9VJD. ToMMV density map is available in EMD-63082, coat protein atomic model is available in PDB-9WQQ. All Python scripts to evaluate and reproduce the results in the paper are present in the Supplementary Materials.

## Abbreviations

CGMMV: cucumber green mottle mosaic virus,
ToMMV: tomato mottle mosaic virus,
TMV: tobacco mosaic virus,
Cryo-EM: Cryo-Electron Microscopy,
SASA: Solvent Accessible Surface Area,
BSA: Buried Surface Area,
MD: Molecular Dynamics

## Notes

### Competing Interest Statement

The authors have declared no competing interest.

### Summary of Updates

Cryo-EM data collection details, Abstract, Discussion, Supplementary data

