## Supplementary for "Comparative cryo-EM studies reveal structural aspects of host specificity for tobamoviruses"

*Ananya Chatterjee et al.*

**This file includes:**

Supplementary
Figs. S1 to S25
Tables S1 to S6
Text S1 to S4

**Other Supplementary Materials for this manuscript include the following:**
Movie S1

**Supplementary Figure S1: Density fit for major H-bonds**

**Supplementary Figure S1a: Map-model fitting for Arg93-Glu96-Asp117 tri-subunit lateral & axial H-bond (CGMMV)**

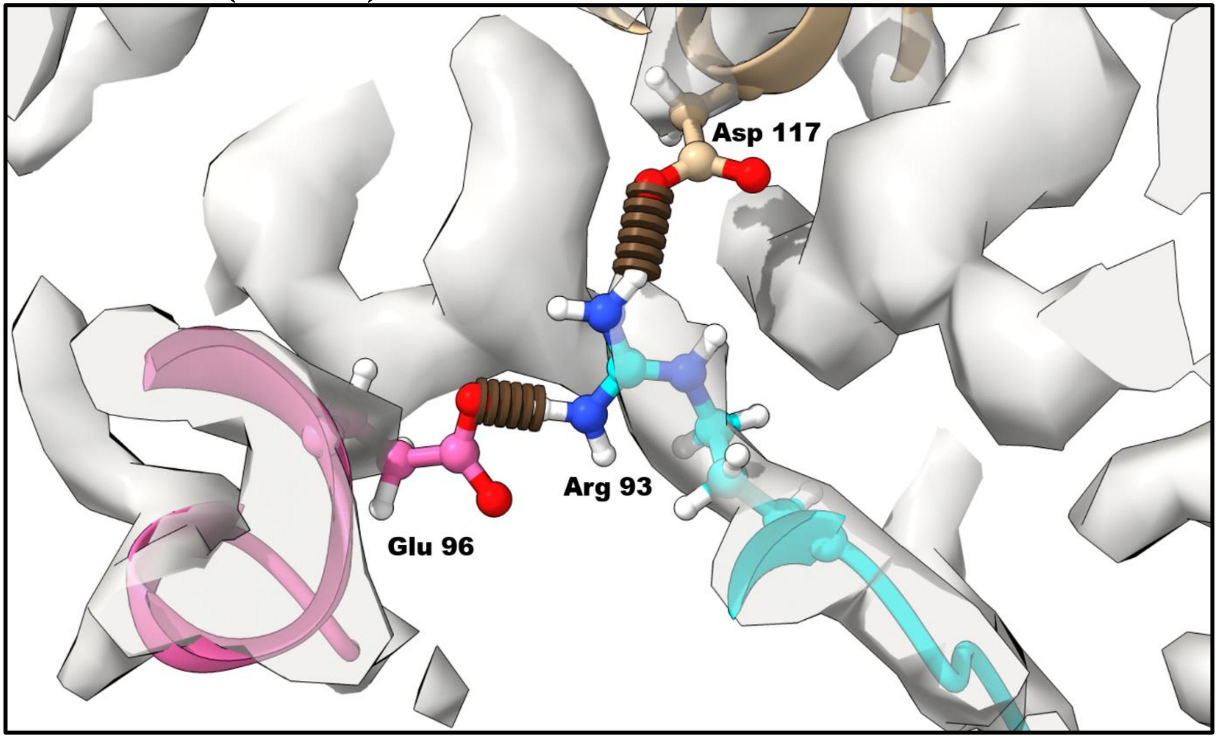


Supplementary Figure (S1a): Map density fit for the Arg93-Glu96-Asp117 tri-subunit lateral and axial interaction network in CGMMV. The threshold value for visualization is 1.85.

**Supplementary Figure S1b: Map-model fitting for Arg93-Glu96-Asp117 tri-subunit lateral & axial H-bond (ToMMV)**

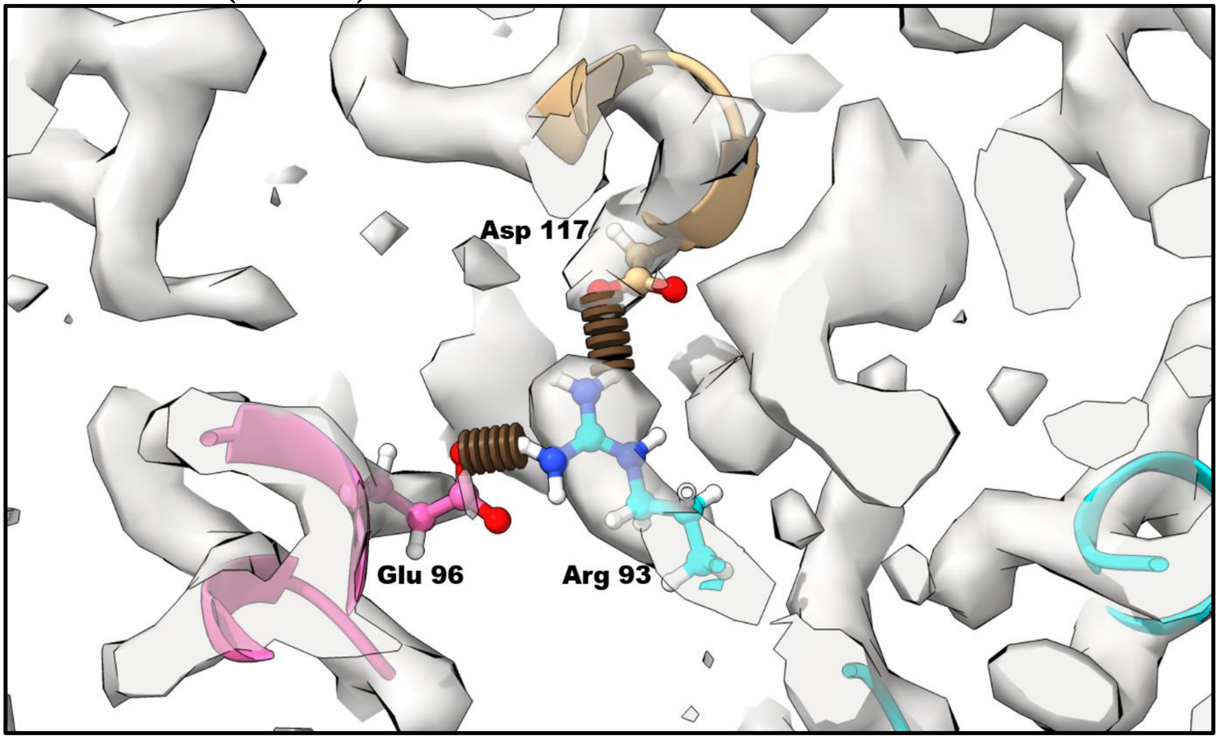


Supplementary Figure (S1b): Map density fit for the Arg93-Glu96-Asp117 tri-subunit lateral and axial interaction network in ToMMV. The threshold value for visualization is 0.85.

**Supplementary Figure S1c: Map-model fitting for Arg123-Asp89 lateral intersubunit H-bond (CGMMV)**

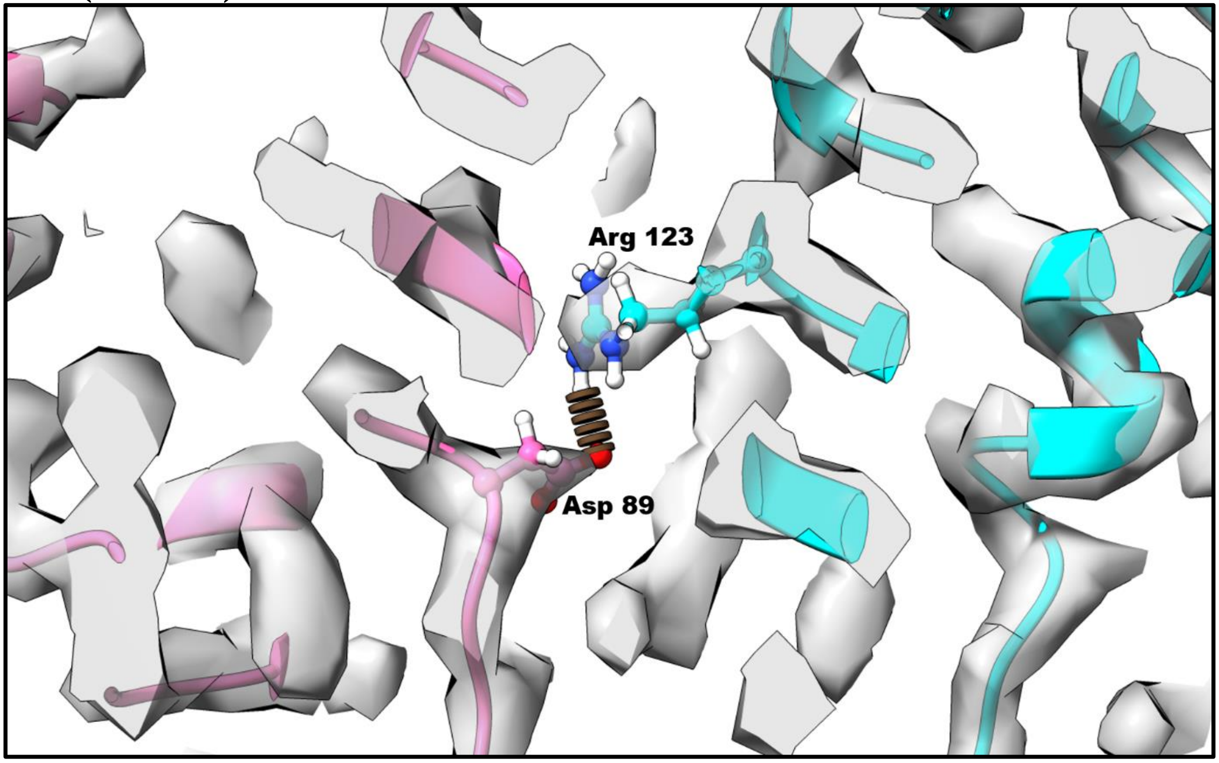


Supplementary Figure (S1c): Map density fit for the Arg123-Asp89 bi-subunit lateral interaction network in CGMMV. The threshold value for visualization is 1.85.

**Supplementary Figure S1d: Map-model fitting for Arg123-Asp89 lateral intersubunit H-bond (ToMMV)**

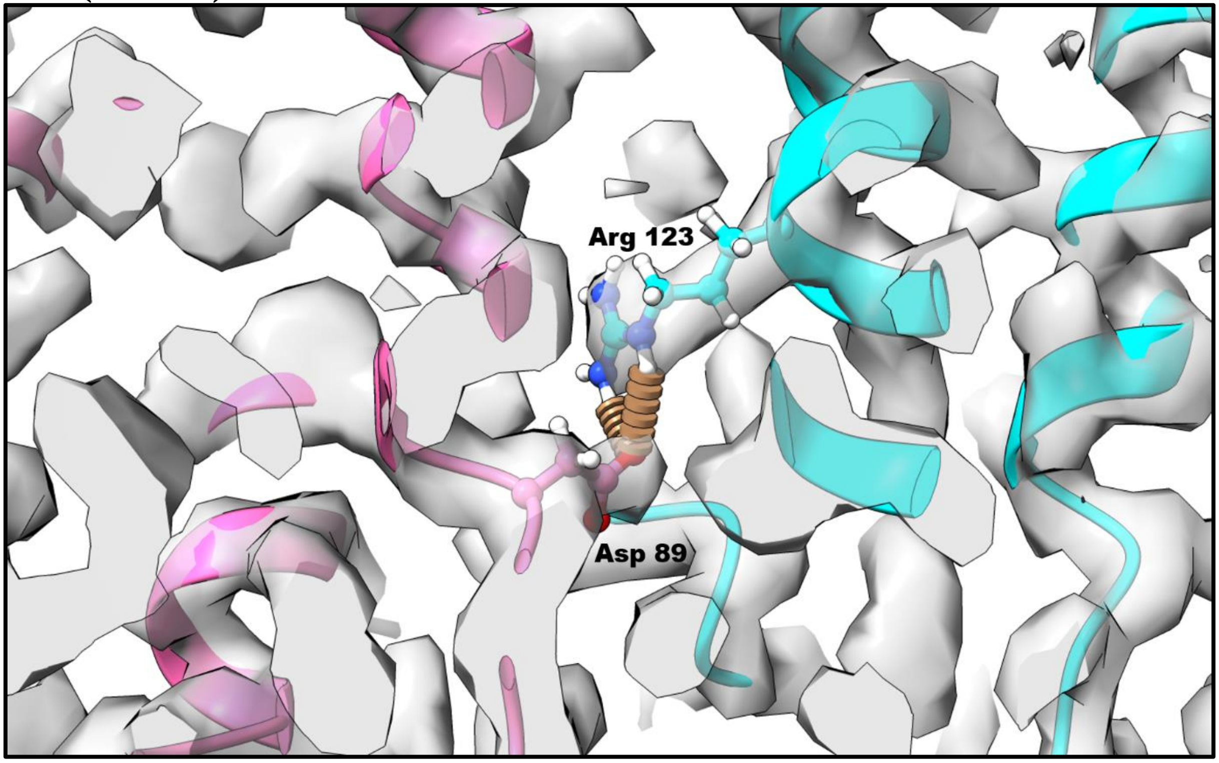


Supplementary Figure (S1d): Map density fit for the Arg123-Asp89 bi-subunit lateral interaction network in ToMMV. The threshold value for visualization is 0.85.

**Supplementary Figure S2: Subunit chain arrangement of 9-mer assembly taken for ΔSASA analysis**

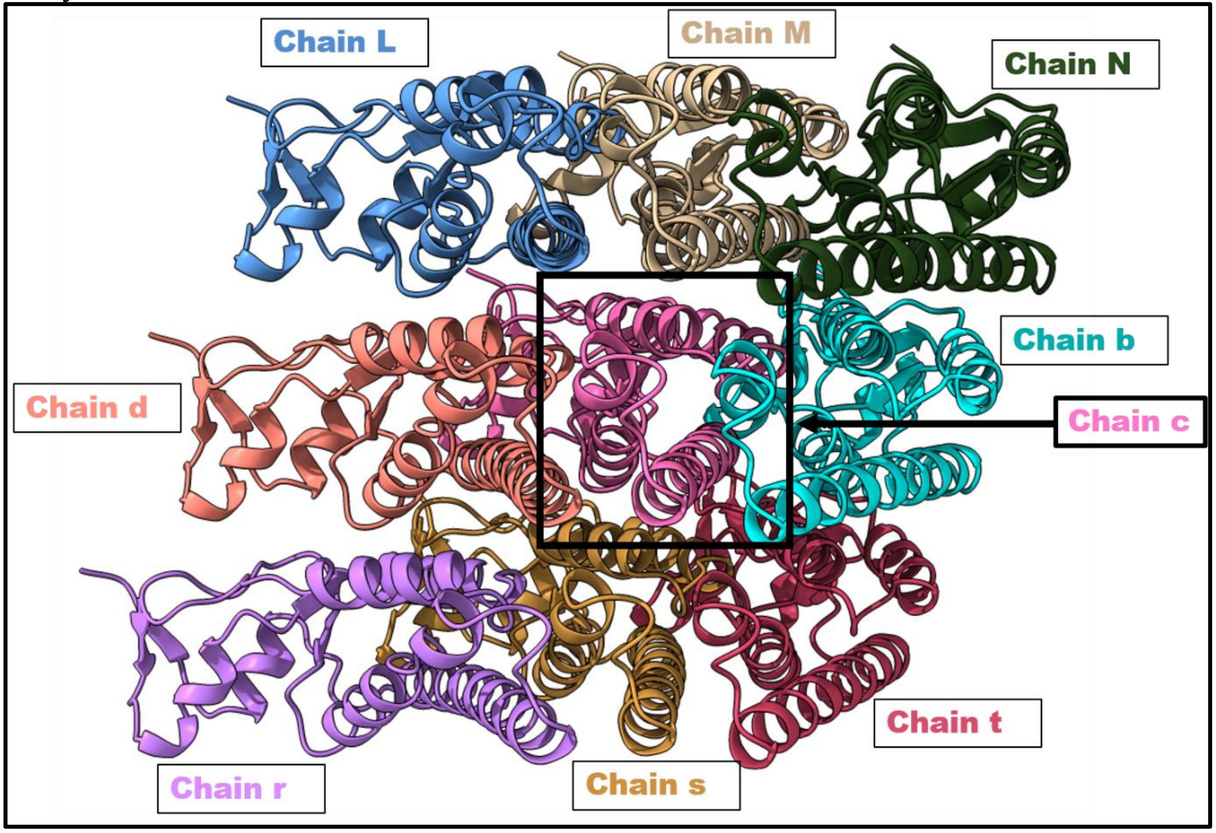


Supplementary Figure S2: Cartoon representation showing the spatial arrangement of surrounding subunits around the central target subunit (Chain c, boxed in black) used for surface area buried calculation. The calculation is done based on this central chain c. Chain numbering is done in ChimeraX by propagating chain A to 49 chains. The numbers are done as chain 1:A, 2:B, ….; when 26 letters are exhausted, the chain numbering shifts to 27:a, 28:b, …. (In lowercase letter).

**Supplementary Figure S3: Surface hydrophobicity of a 9-mer assembly**

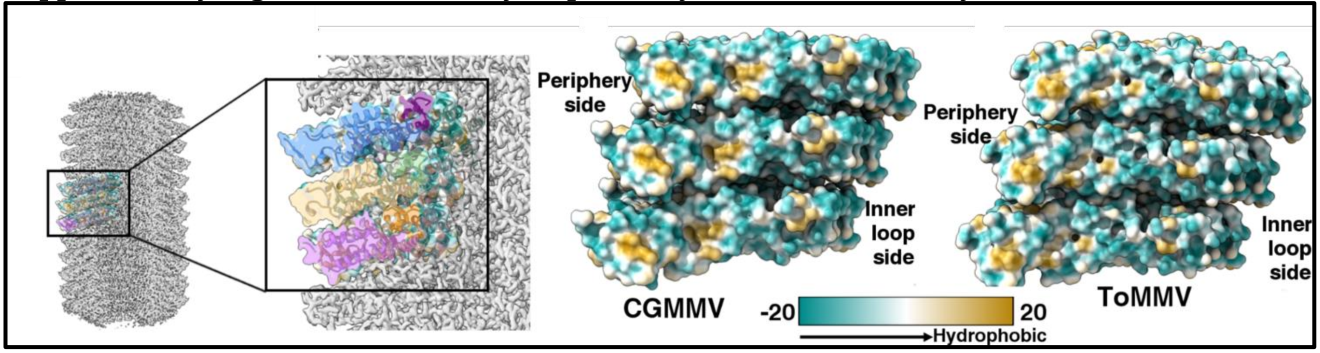


Supplementary Figure S3: Surface hydrophobicity distribution is shown as comparative surface maps coloured by the Kyte-Doolittle scale (cyan: hydrophilic; gold: hydrophobic), where CGMMV exhibits prominent hydrophobic patches at the interfaces, whereas ToMMV displays a dispersed hydrophobic surface profile.

**Supplementary Figure S4: Comparison of RNA map boxed density and whole map**

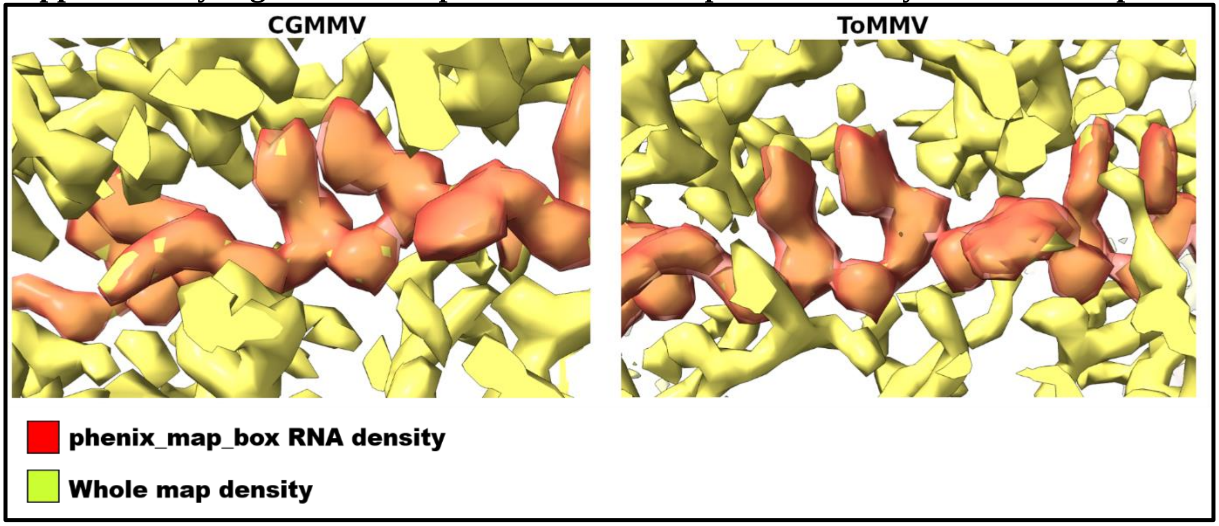


Supplementary Figure S4: Density map visual comparison highlighting the map-boxed single-stranded RNA density fitted within the inner groove of CGMMV (left) and ToMMV (right) capsid whole maps. Threshold level: CGMMV whole map: 2.13, cut-out RNA: 1.75; ToMMV whole map: 1.57, cut-out RNA: 1.08.

**Supplementary Figure S5: Sequence logo for inner loop**

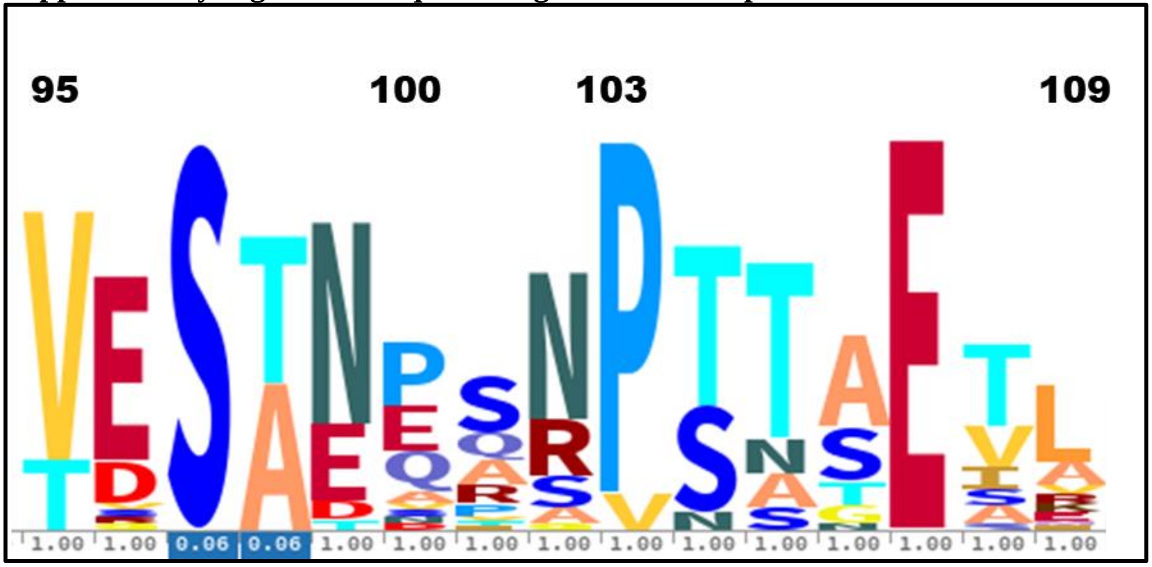


Supplementary Figure S5: Sequence logo of the inner loop of Tobamoviruses showing a majority of the viruses carrying Pro103.

**Supplementary Figure S6: End-to-end helix picking in Relion manual picking for CGMMV**

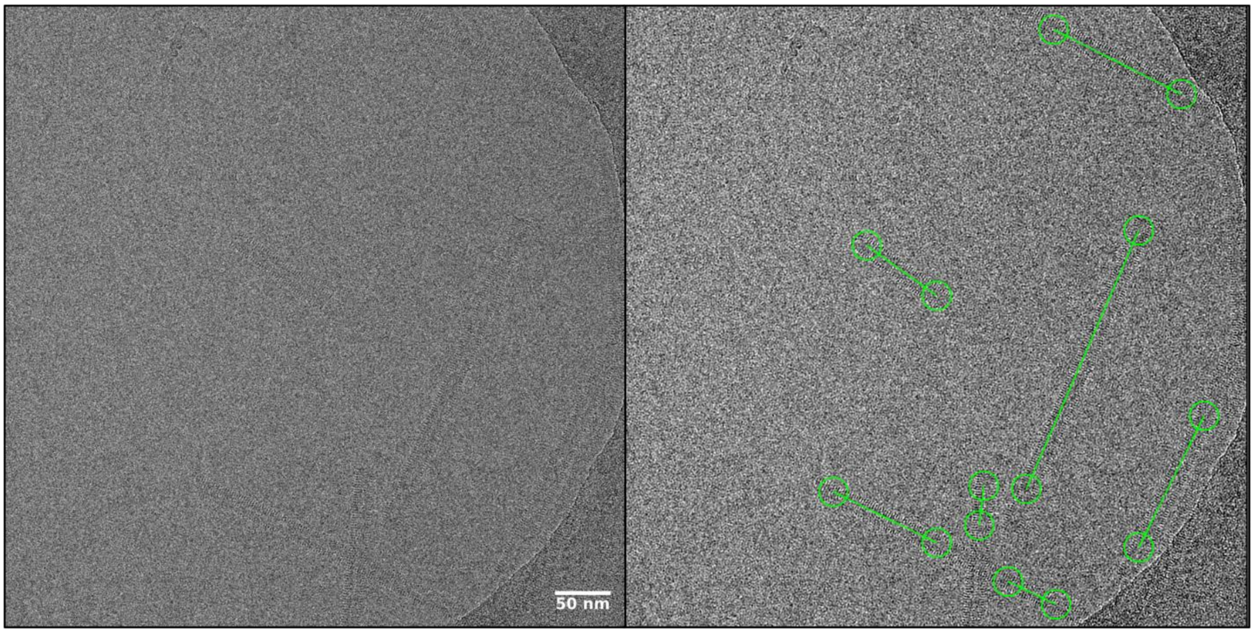


Supplementary Figure (S6): Representative cryo-EM raw micrograph showing end-to-end trace selection (green lines) of helical CGMMV filaments.

**Supplementary Figure S7: End-to-end helix picking in Relion manual picking for ToMMV**

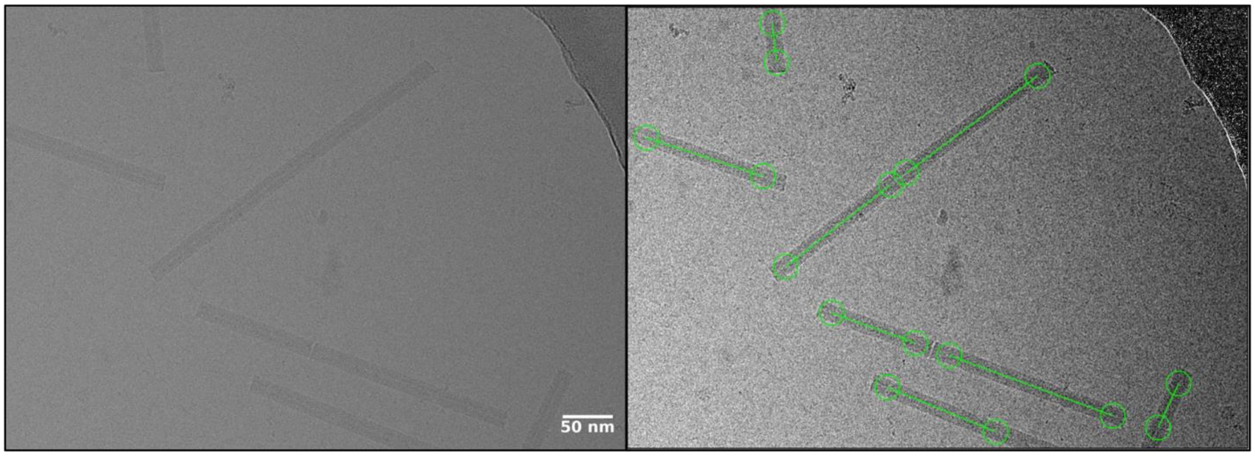


Supplementary Figure (S7): Representative cryo-EM raw micrograph showing manual picking and selection of ToMMV helical assemblies.

**Supplementary Figure S8: 2D class averages selected for helical refinement in CryoSPARC for CGMMV**

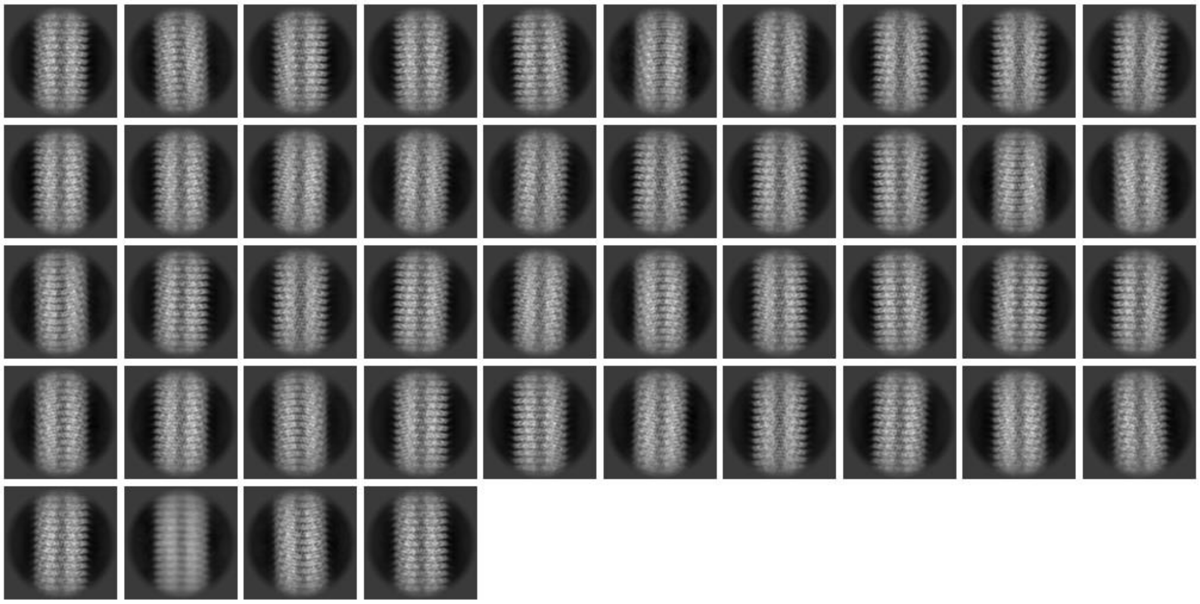

Supplementary Figure (S8): Selected 2D class averages (44) displaying well-defined secondary structure features used for subsequent helical processing.

**Supplementary Figure S9: 2D class averages selected for helical refinement in CryoSPARC for ToMMV**

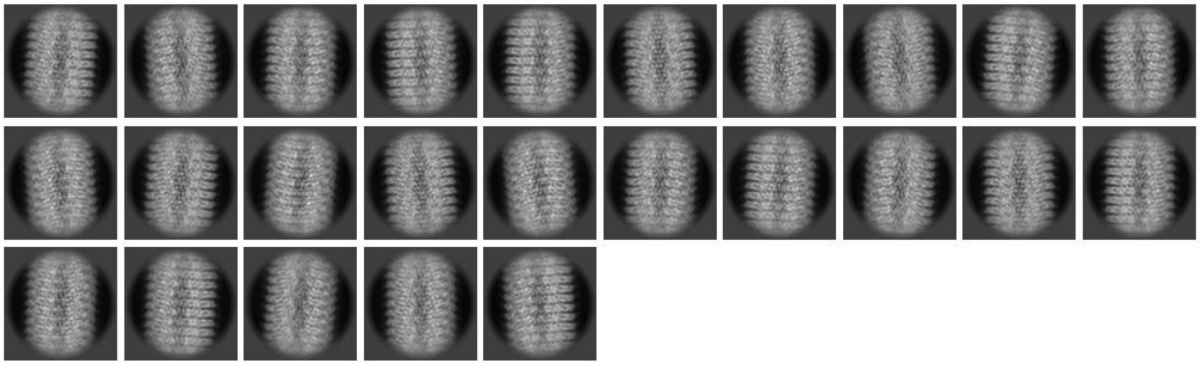

Supplementary Figure (S9): Selected 2D class averages (25) displaying well-defined secondary structure features used for subsequent helical processing.

**Supplementary Figure S10: Determination of helix parameters by indexing the diffraction pattern of 2D class averages for CGMMV using PyHI**

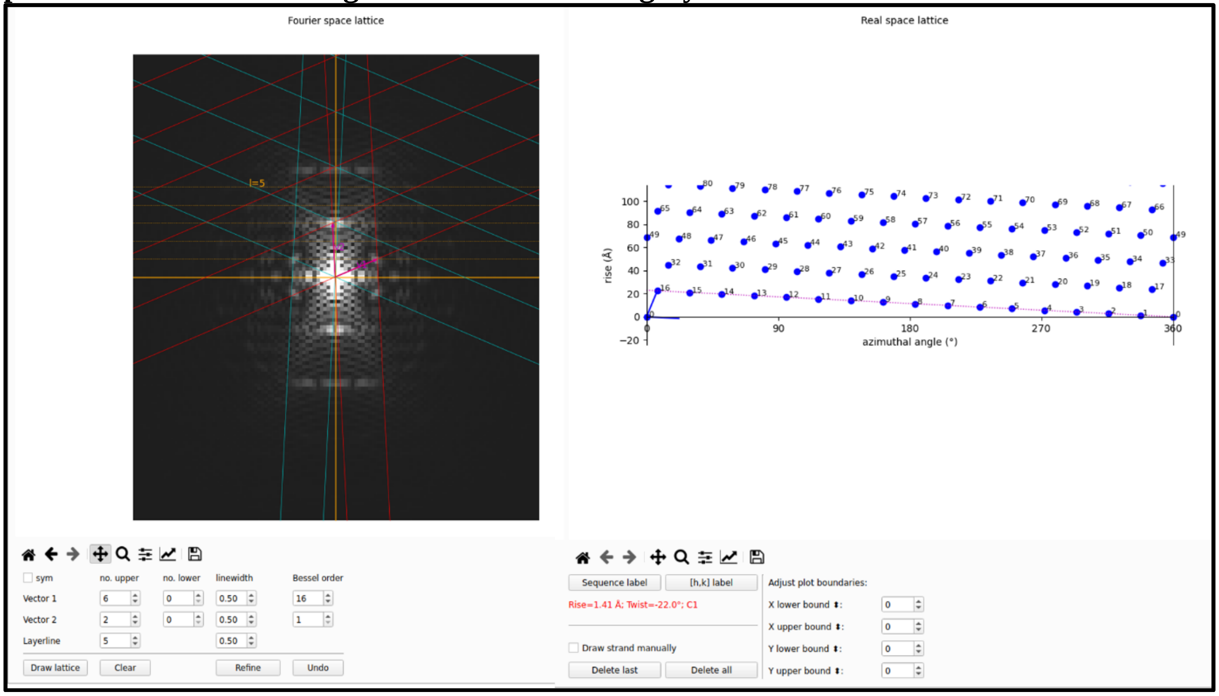


Supplementary Figure (S10): Indexing of the diffraction pattern obtained from 2D class averages in Fourier space (left) and the corresponding real-space lattice determination (right) for CGMMV.

**Supplementary Figure S11: Determination of helix parameters by indexing the diffraction pattern of 2D class averages for ToMMV using PyHI**

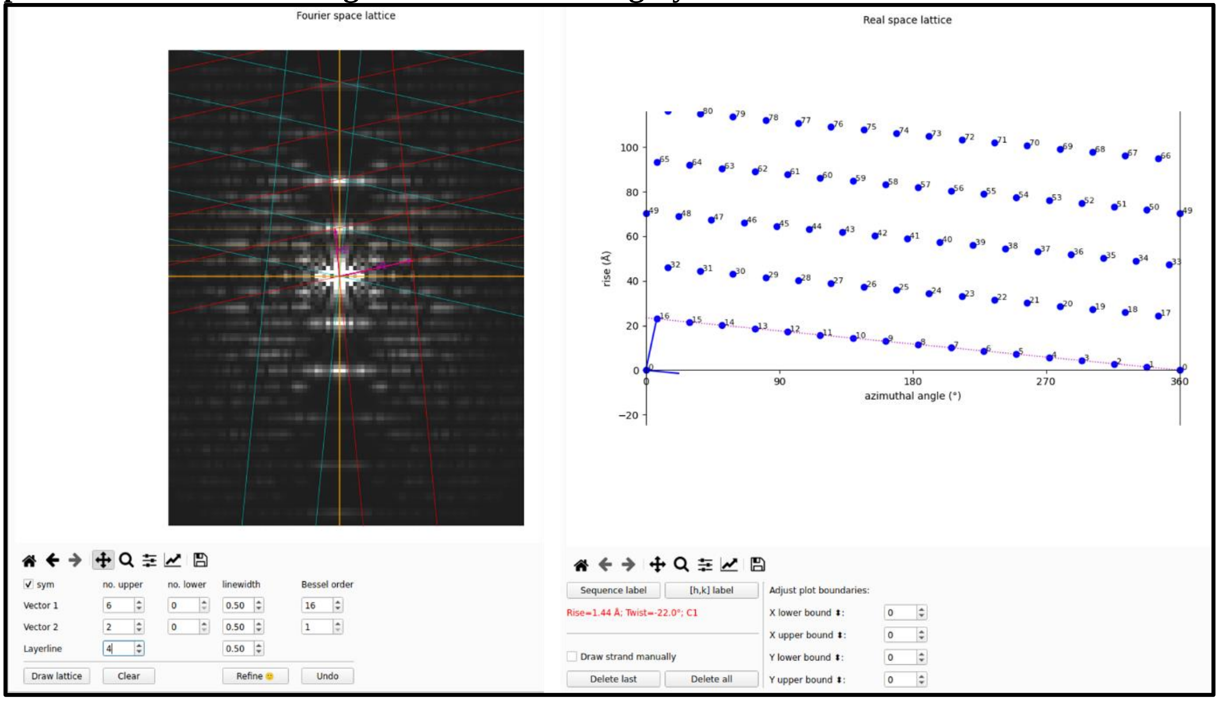


Supplementary Figure (S11): Fourier-space lattice indexing (left) and real-space lattice mapping (right) used to determine rise and twist parameters for ToMMV.

**Supplementary Figure S12: Fourier Shell Correlation (FSC) plot for CGMMV**

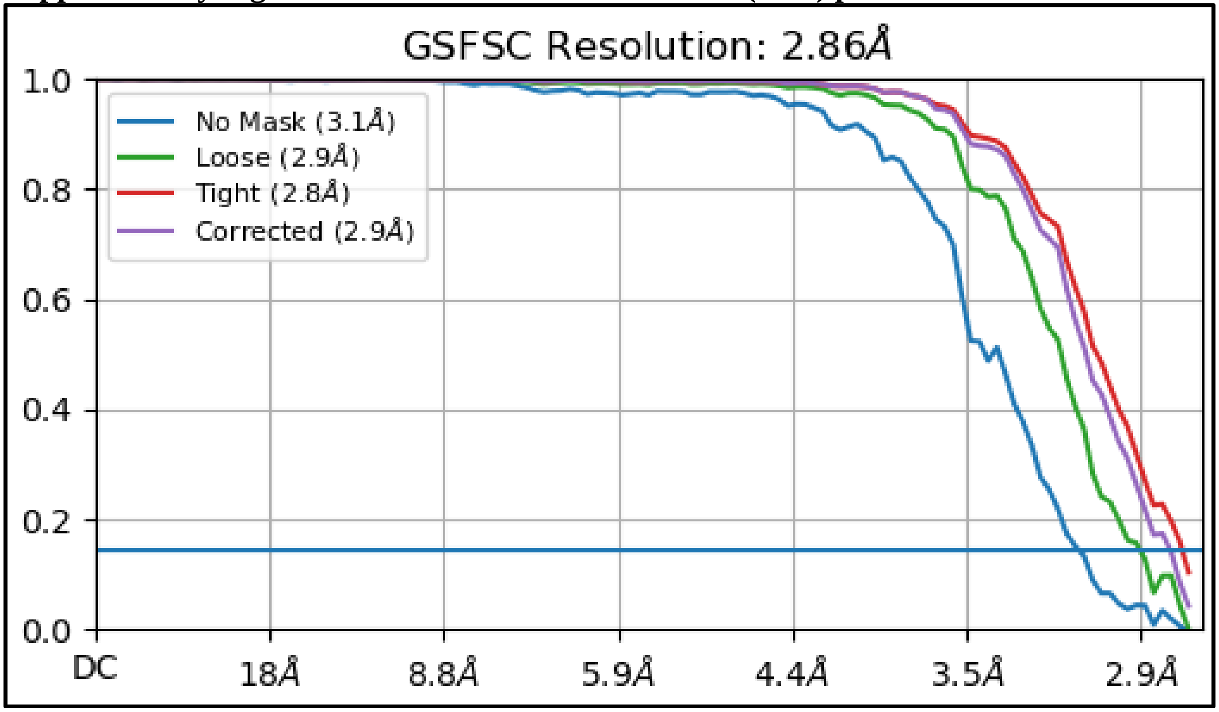


Supplementary Figure (S12): FSC curves calculated at Gold-standard cut-off 0.143, indicating a final global resolution of 2.86 Å.

**Supplementary Figure S13: Fourier shell Correlation (FSC) plot for ToMMV**

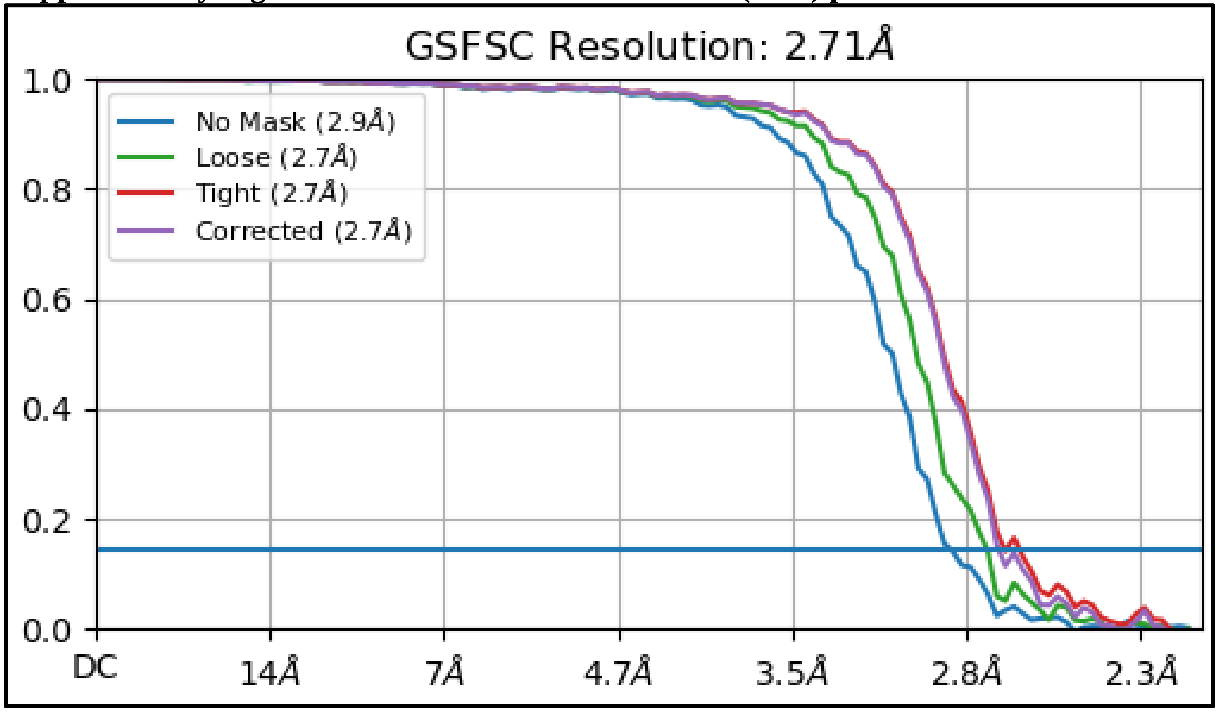


Supplementary Figure (S13): FSC curves calculated at Gold-standard cut-off 0.143, indicating a final global resolution of 2.71 Å.

**Supplementary Figure S14: Local resolution distribution histogram for CGMMV**

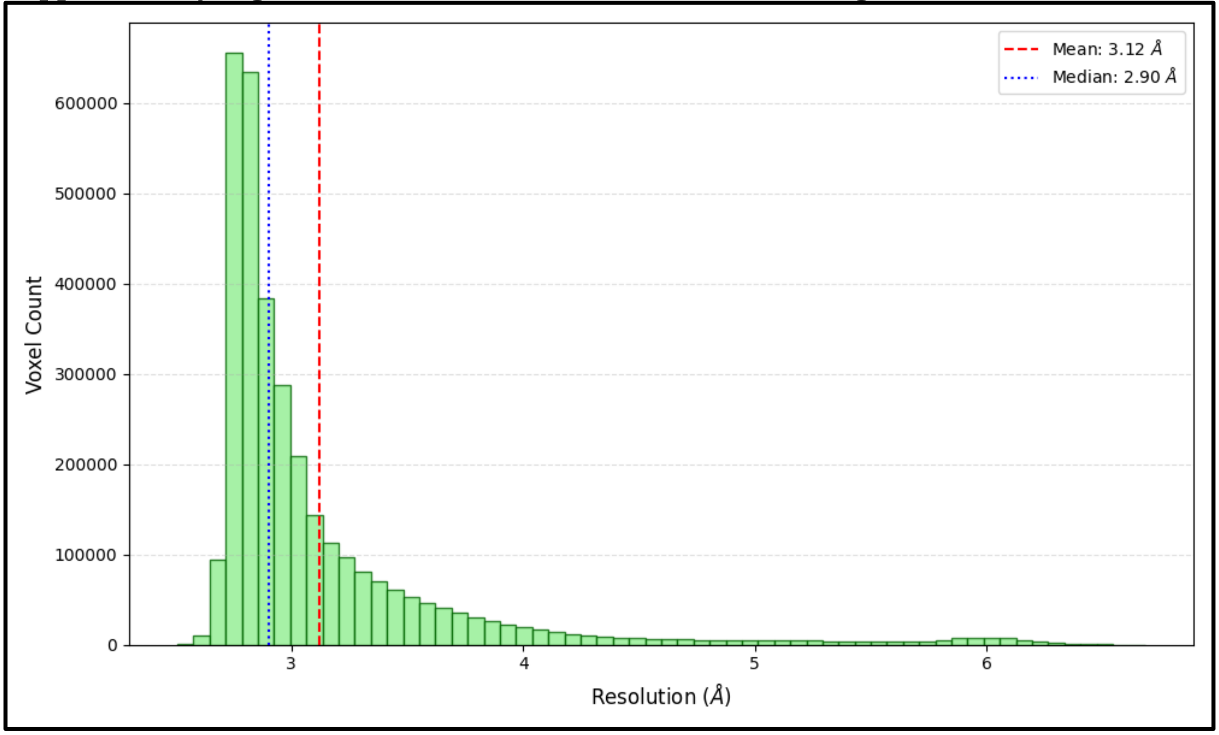


Supplementary Figure (S14): Distribution of local map resolution across overall map voxels, showing a median resolution of 2.90 Å and a mean of 3.12 Å.

**Supplementary Figure S15: Local resolution distribution histogram for ToMMV**

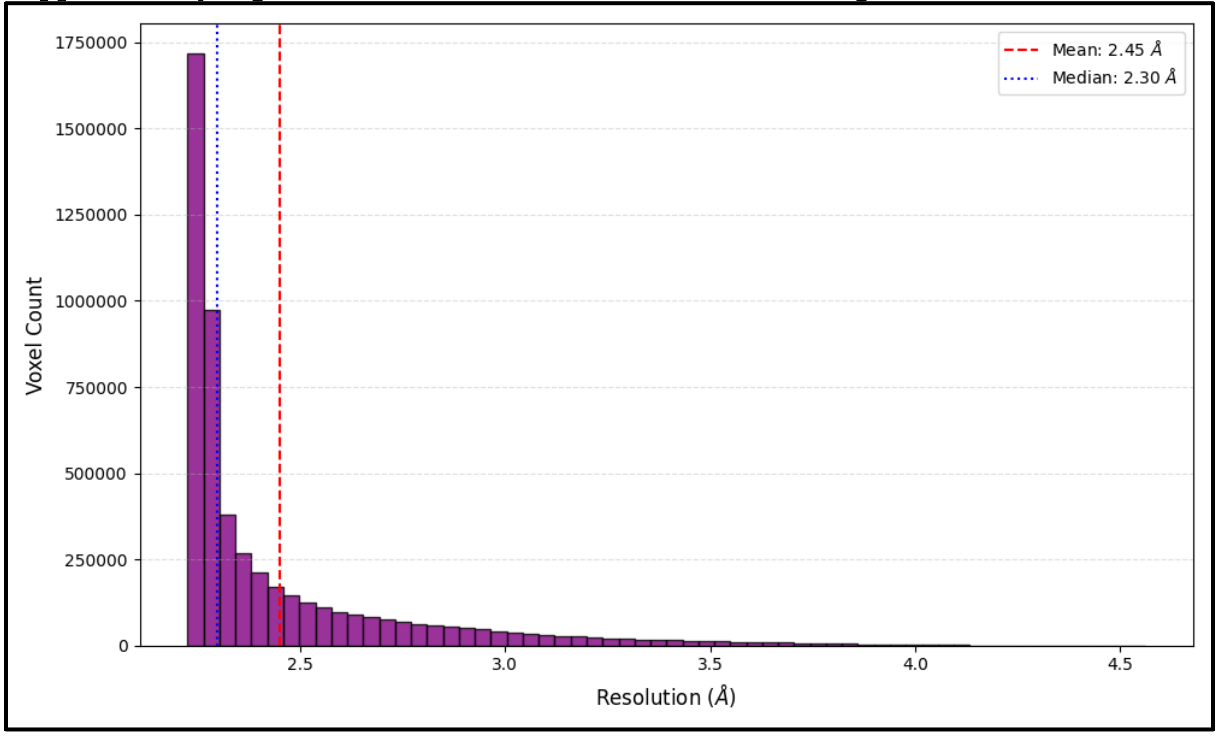


Supplementary Figure (S15): Distribution of voxel counts across local resolution ranges, displaying a median resolution of 2.30 Å and a mean of 2.45 Å.

**Supplementary Figure S16: Local resolution distribution for CGMMV**

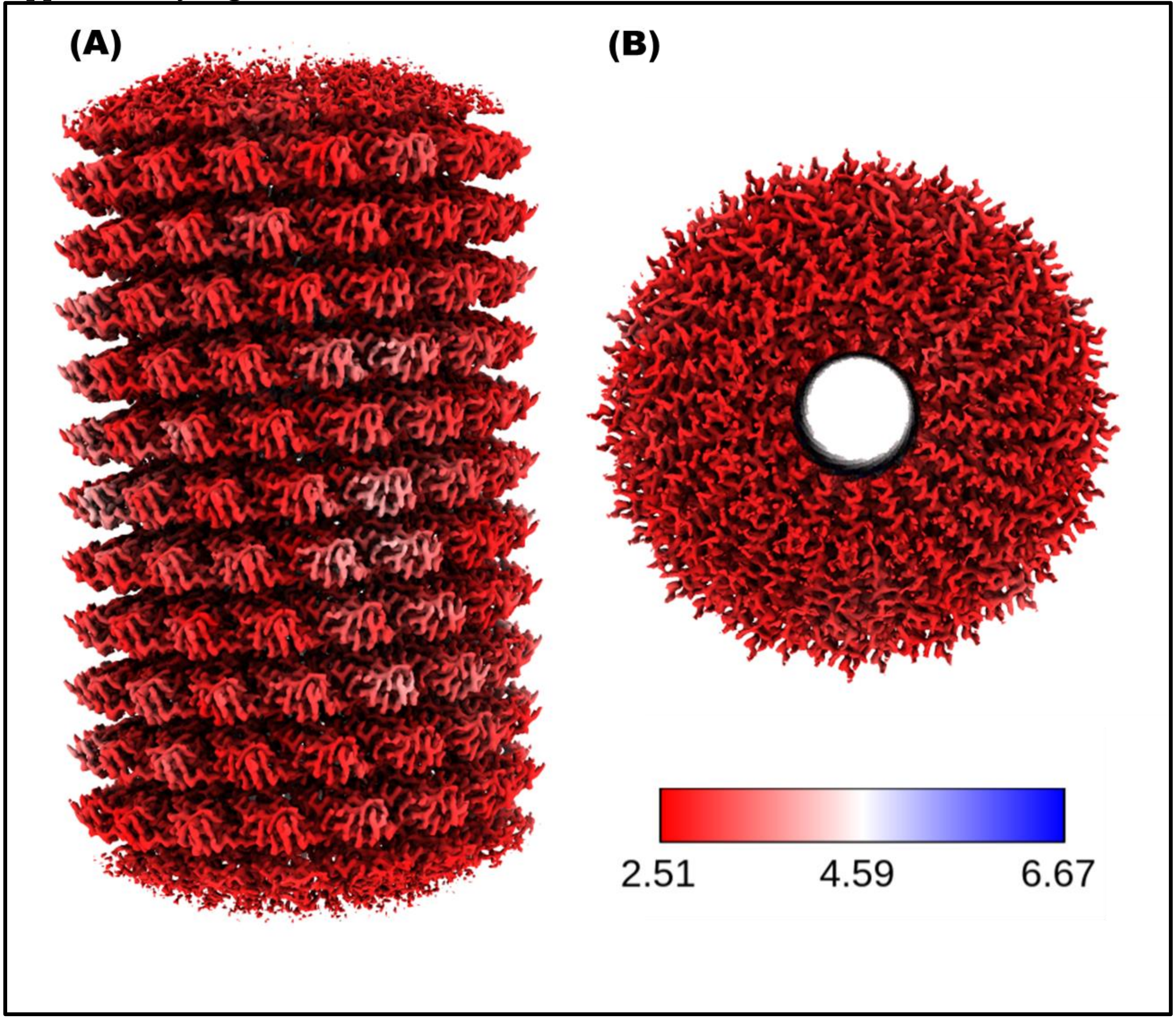


Supplementary Figure (S16): CGMMV map coloured according to local resolution range. The resolution range in Å is provided in the inset. (A) Side view (B) Top view.

**Supplementary Figure S17: Local resolution distribution for ToMMV**

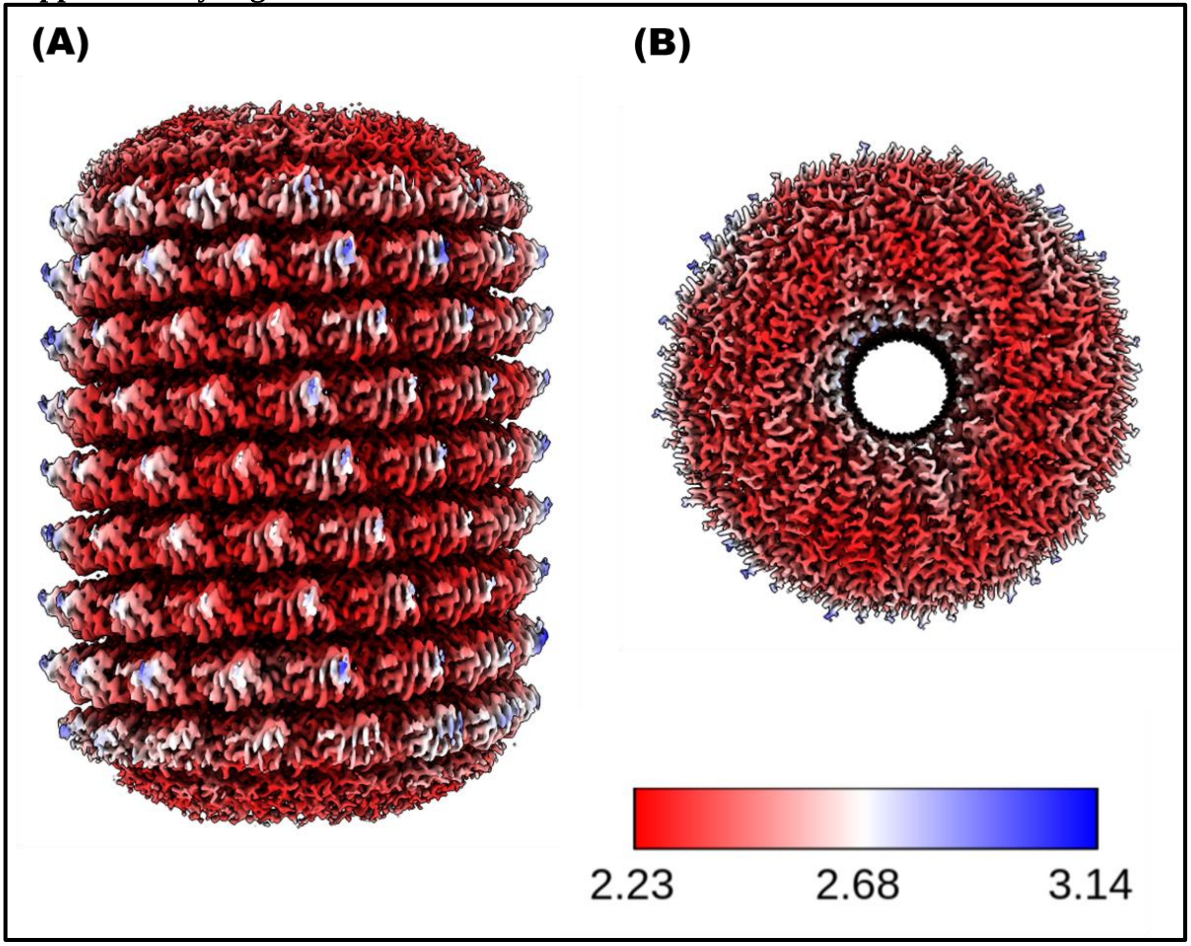


Supplementary Figure (S17): ToMMV map coloured according to local resolution range. The resolution range in Å is provided in the inset. (A) Side view; (B) Top view.

**Supplementary Figure S18: Map vs model comparison of CGMMV**

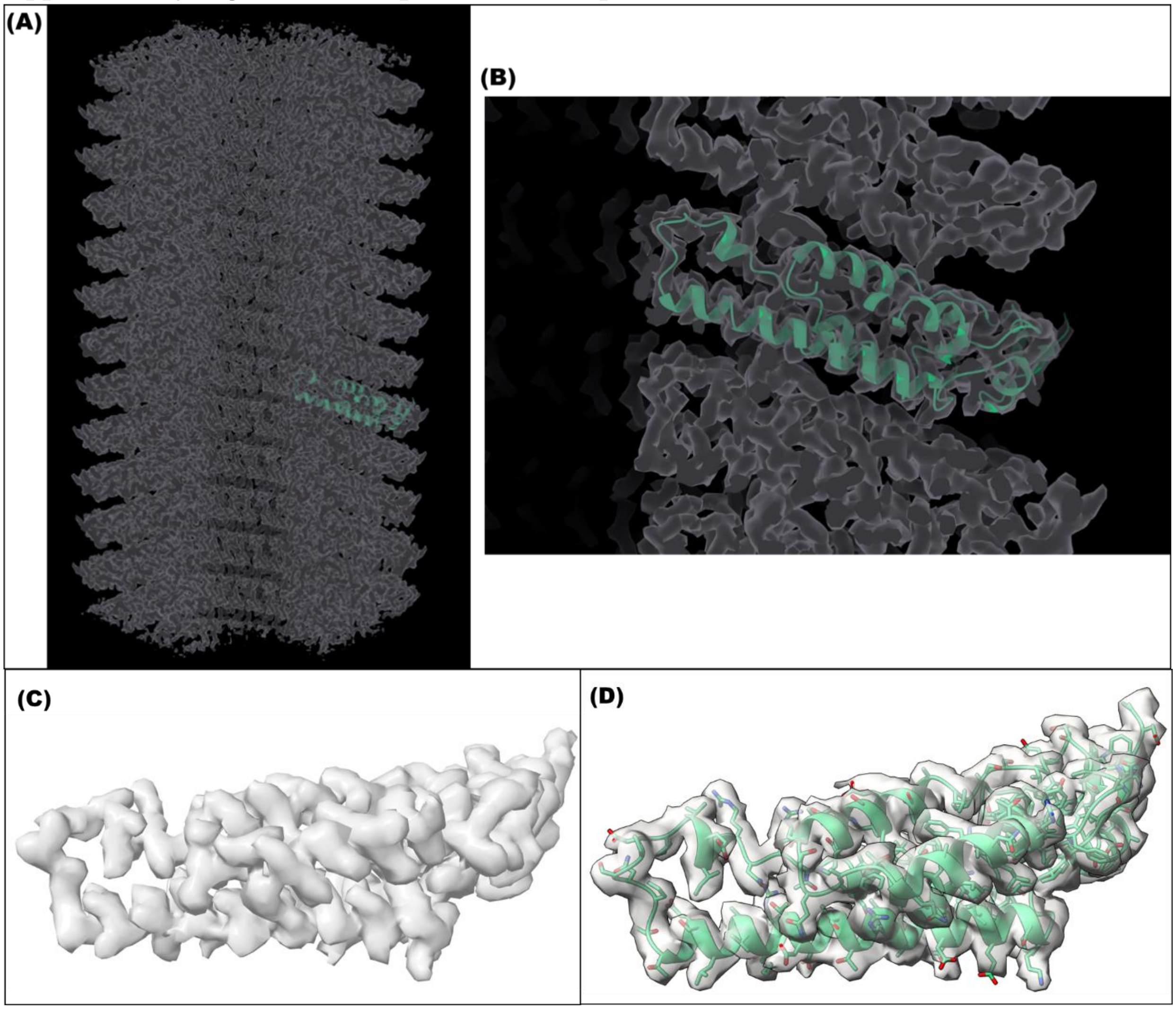


Supplementary Figure S18: For CGMMV (A) Map boxed density from a single subunit fitted with the modelled pdb after phenix.real_space_refinement and COOT manual refinement. Map-model CC=0.9. (B) Zoomed in view of A. (C) Density from a single subunit using phenix.map_box (D) Map boxed density from a single subunit fitted with the final pdb, Map-model CC=0.94. Threshold for visualization for A & B is 1.55; for C & D is 0.431.

**Supplementary Figure S19: Map vs model comparison of ToMMV**

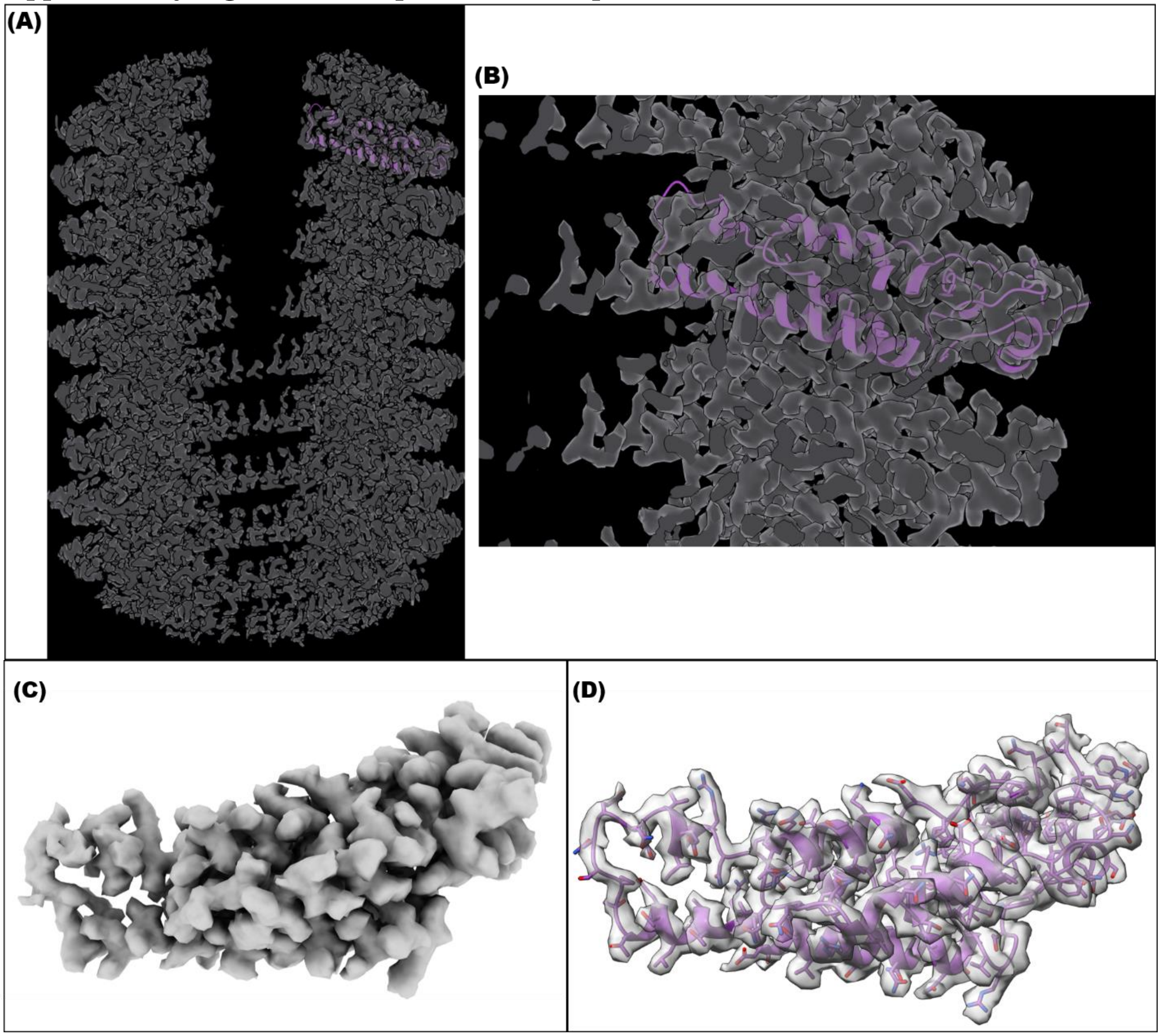


Supplementary Figure S19: For ToMMV (A) Map boxed density from a single subunit fitted with the modelled pdb after phenix.real_space_refinement and COOT manual refinement. Map-model CC=0.87. (B) Zoomed in view of A. (C) Density from a single subunit using phenix.map_box (D) Map boxed density from a single subunit fitted with the final pdb, Map-model CC=0.88. Threshold for visualization for A & B is 0.55; for C & D is 0.368.

**Supplementary Figure S20: Amino acid side chain visualization for CGMMV**

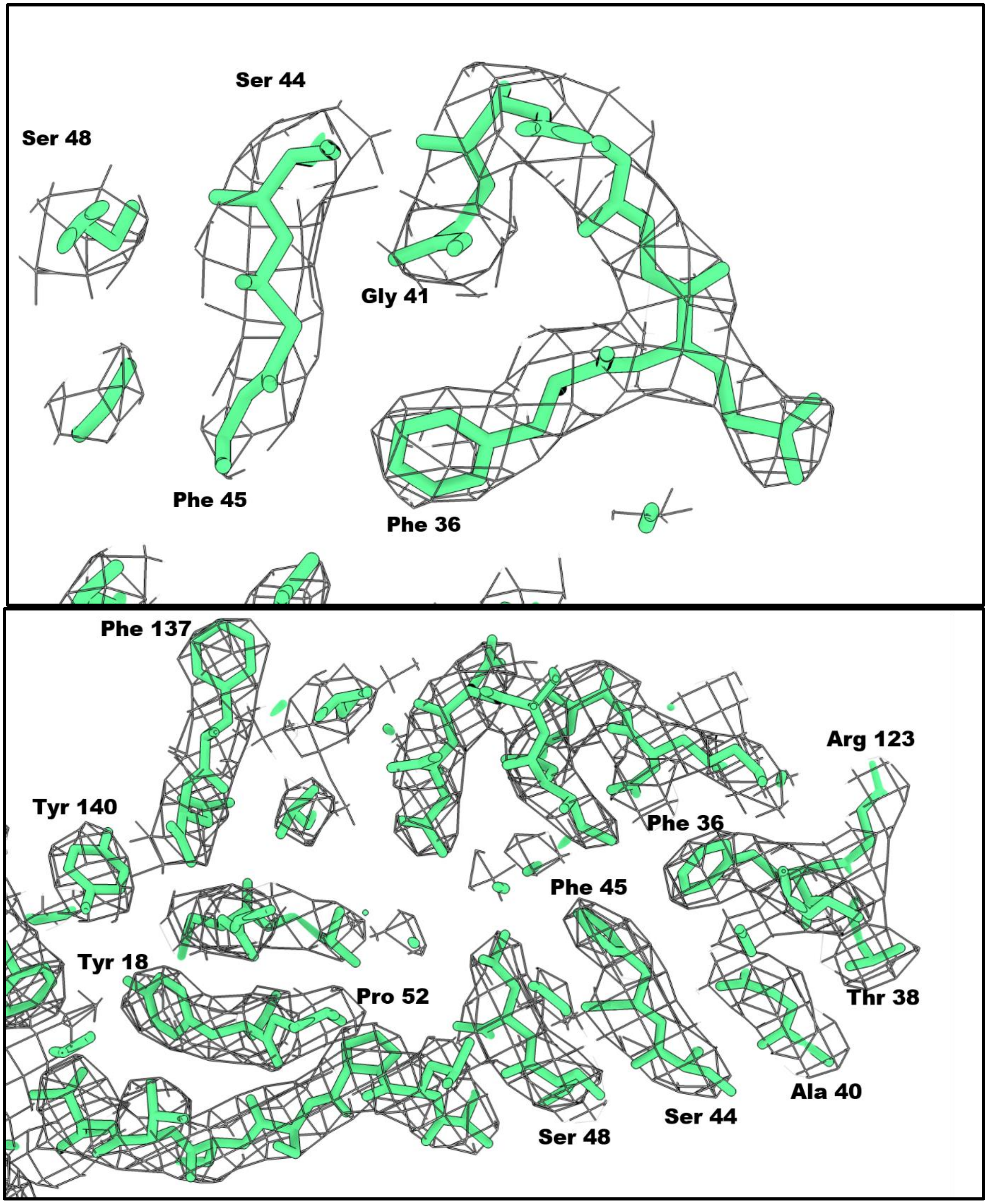


Supplementary Figure (S20): Amino acid side chain visualization in single subunit map-model fitting for CGMMV. Map-model correlation is 0.94.

**Supplementary Figure S21:  Amino acid side chain visualization for ToMMV**

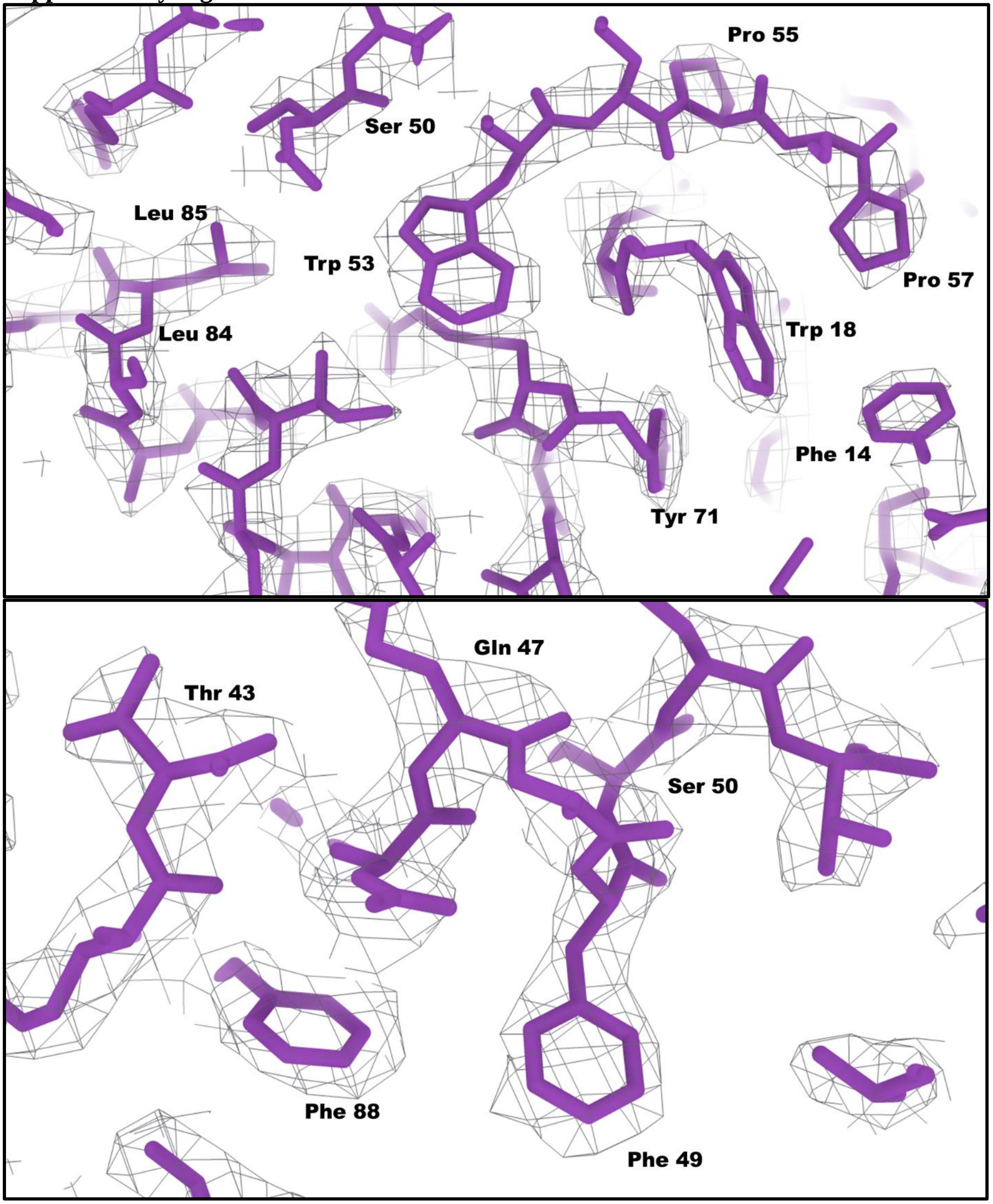


Supplementary Figure (S21): Amino acid side chain visualization in single subunit map-model fitting for ToMMV. Map-model correlation is 0.88.

**Supplementary Figure S22: Map of boxed RNA density local resolution for CGMMV**

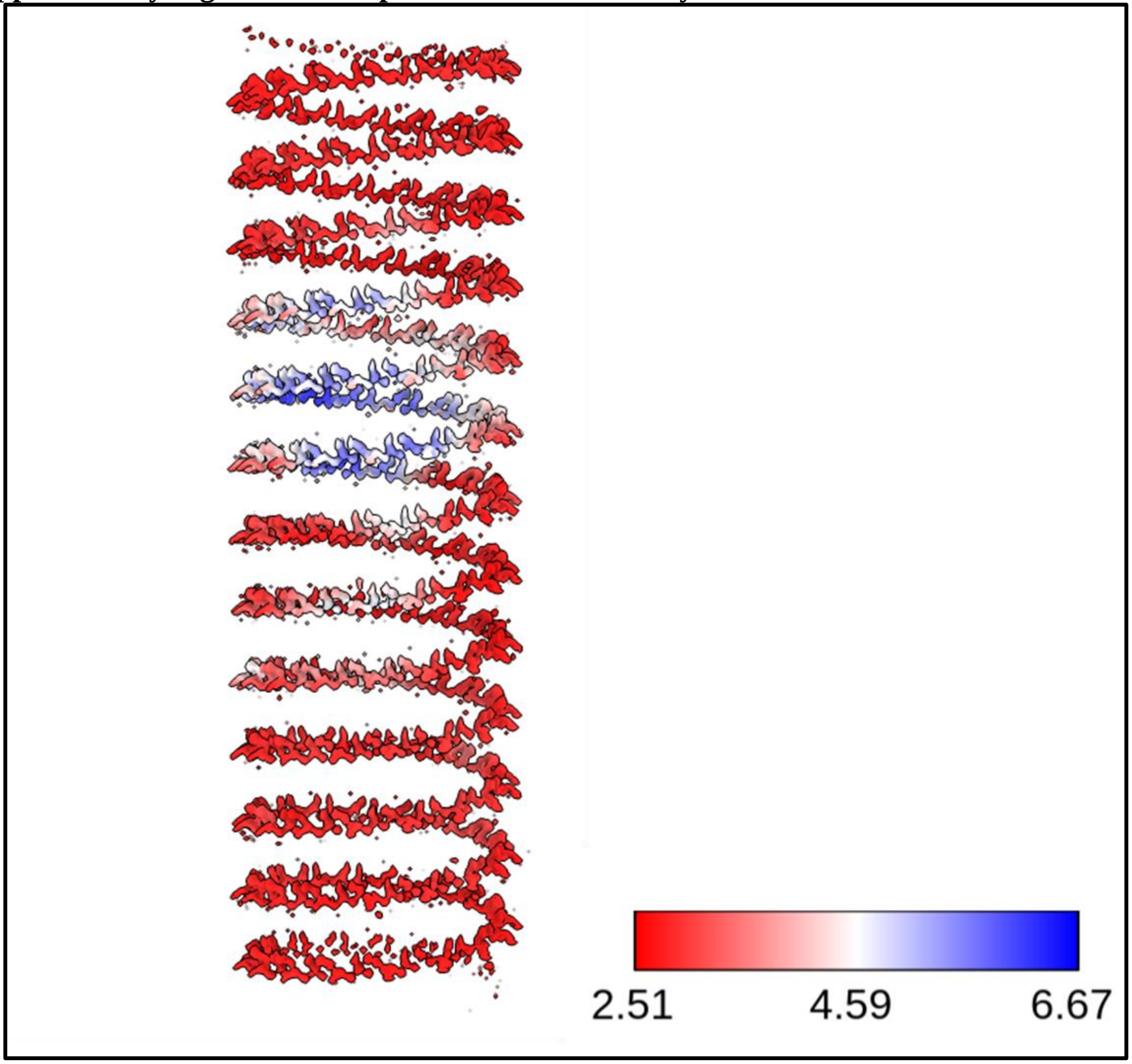


Supplementary Figure (S22): Map of boxed RNA density coloured according to local resolution ranges for CGMMV. The resolution range in Å is provided in the inset.

**Supplementary Figure S23: Map of boxed RNA density local resolution for ToMMV**

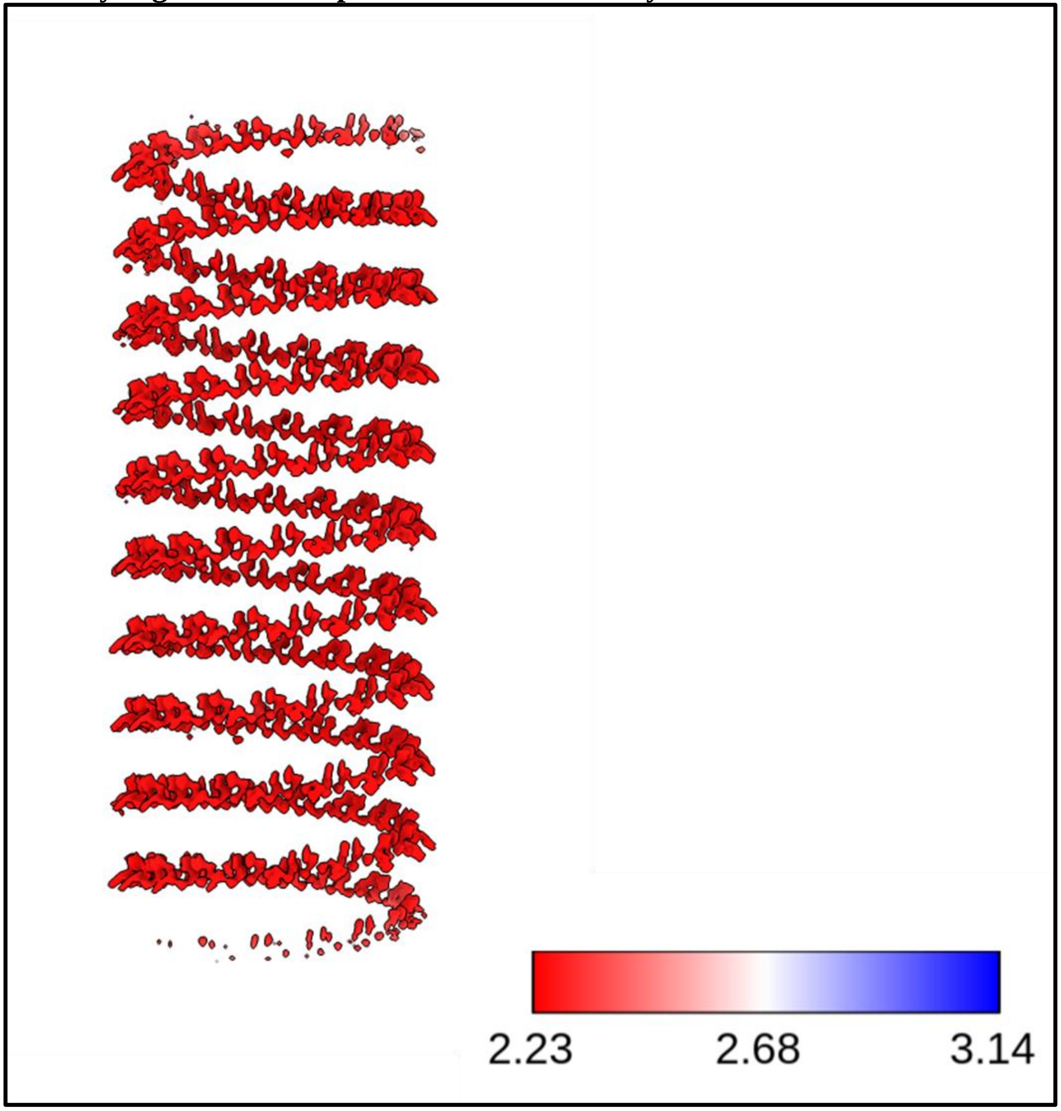


Supplementary Figure (S23): Map of boxed RNA density coloured according to local resolution ranges for ToMMV. The resolution range in Å is provided in the inset.

**Supplementary Figure S24: Coat protein subunit and modelled trinucleotide in the RNA-associated density**

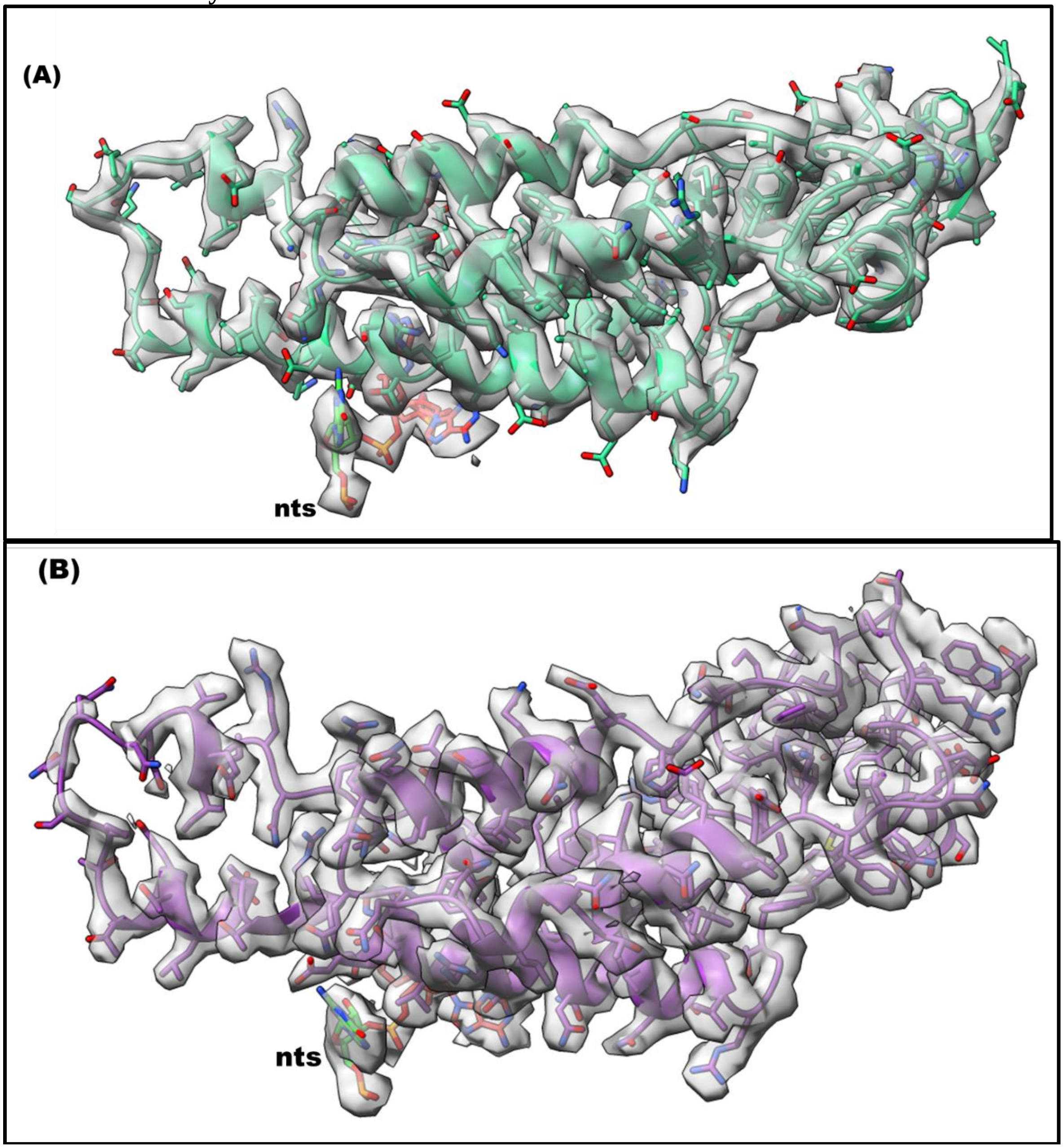


Supplementary Figure (S24): In the helically averaged RNA-associated density, three nucleotides are modelled. The coat protein subunit model and the nts model are fitted together with map box density of a single subunit and associated nucleotides for (A) CGMMV and (B) ToMMV.
Threshold for visualization for CGMMV (A): 1.55; Map-model CC: 0.92
Threshold for visualization for ToMMV (B): 0.55; Map-model CC: 0.86

**Supplementary Figure S25: Workflow for cryo-EM data processing, helical reconstruction, and atomic model refinement of CGMMV and ToMMV**

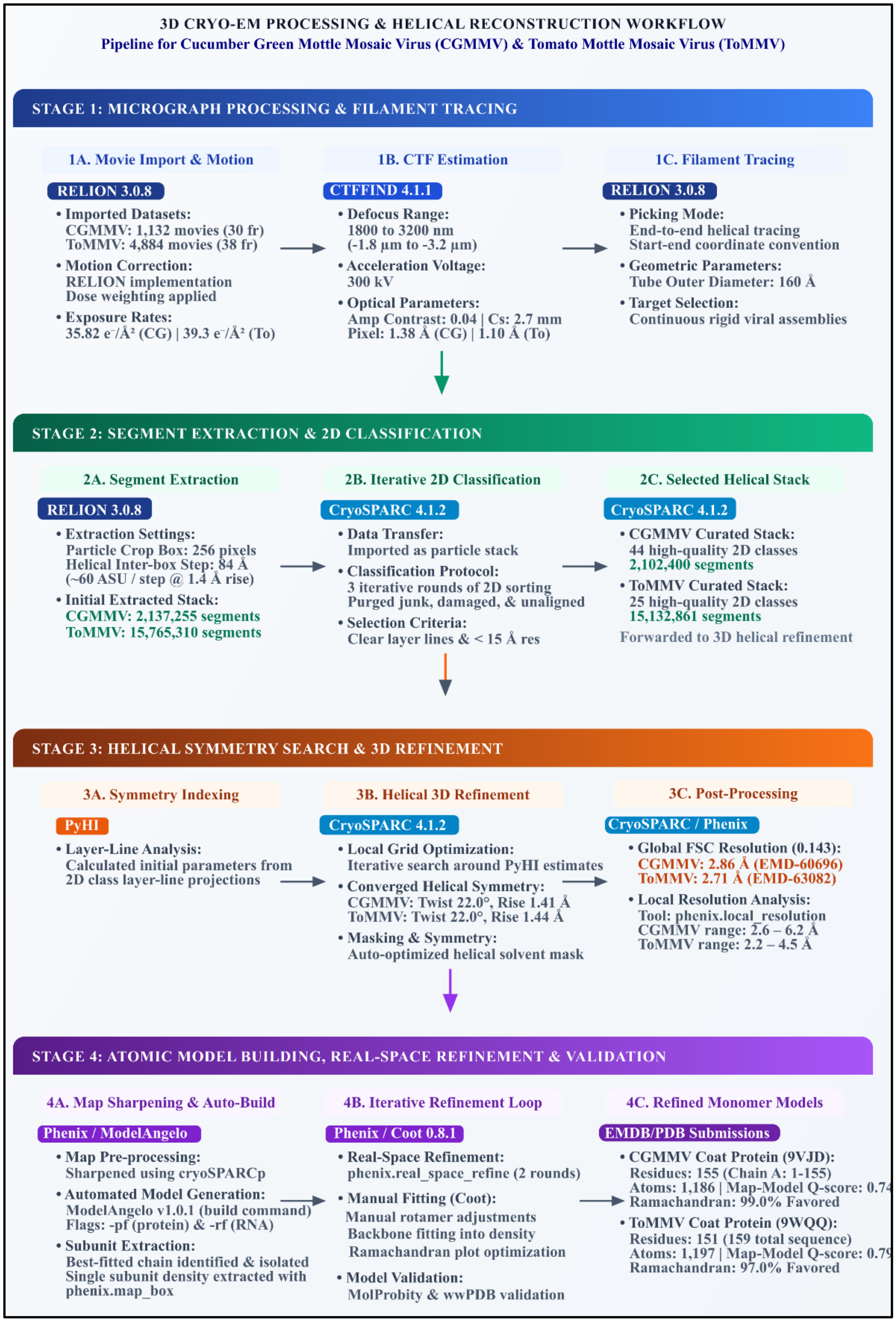


Supplementary Figure S25: Helical processing and refinement pipeline for CGMMV and ToMMV. The stages show filament processing, segment extraction & 2D classification, symmetry indexing & 3D reconstruction, model building & downstream analysis.

**Supplementary Table 1: Detailed Hydrogen bonds in 3-turn 49-mer assembly of CGMMV and ToMMV**

| Interaction Type | CGMMV | | | | ToMMV | | | | Significance / Note |
|---|---|---|---|---|---|---|---|---|---|
| | **Donor** | **Acceptor** | **Donor-Acceptor Distance (Å)** | **Donor-Hydrogen-Acceptor Angle (°)** | **Donor** | **Acceptor** | **Donor-Acceptor Distance (Å)** | **Donor-hydrogen-Acceptor Angle (°)** | |
| Lateral (Inter) | Arg93 NH1 (*i*) | Glu96 OE1 (*i+1*) | 3.09 | 177.68 | Arg93 NH1 (*i*) | Glu96 OE1 (*i+1*) | 2.68 | 140.09 | Conserved salt bridge; more compact in ToMMV |
| Lateral (Inter) | Arg123 NH2 (*i*) | Asp89 OD2 (*i-1*) | 3.03 | 167.2 | Arg123 NE, NH2 (i) | Asp89 OD2 (i-1) | 2.67/ 3.38 | 131.5, 170.59 | Bidentate in ToMMV |
| Lateral (Inter) | Arg114 NH1 (*i*) | Asp116 OD1 (*i*+1) | 2.77 | 148.14 | Arg114 NH2 (i) | Asp116 OD2 (i+1) | 3.01 | 172.18 | Lateral latch |
| Axial (Inter) | Arg93 NH2 (*i*) | Asp117 OD1 (*i+16*) | 2.9 | 142.7 | Arg93 NH2 (*i*) | Asp117 OD2 (*i+16*) | 3.06 | 115.67 | Part of tri-subunit coordination |
| Intra-subunit | Arg62 NE, NH2 (*i*) | Glu146 OE1 (*i*) | 3.16 / 3.32 | 159.1, 146.09 | Arg62 NE, NH2 (*i*) | Glu146 OE2 (*i*) | 3.71 | 149.6 | Bidentate in CGMMV; weak in ToMMV |
| Intra-subunit | Arg46 NH1 (*i*) | Asp43 OD1 (*i*) | 3.09 | 160.13 | | | Absent | | Right-slewed helix stabilization |

**Supplementary Table 2a: Comparative ΔSASA analysis of central chain c in a 9-mer assembly**

| Target Chain: | Chain c (Layer 2) | | CGMMV | ToMMV |
|---|---|---|---|---|
| Layer 1 Neighbours: | Chains L, M, N | Candidate Residues: | 160 residues | 120 residues |
| Layer 2 Neighbours: | Chains b, d | Mean Patch ΔSASA: | 363.73 Å² | 304.19 Å² |
| Layer 3 Neighbours: | Chains r, s, t | Max Patch ΔSASA: | 558.12 Å² (Patch 1) | 535.48 Å² (Patch 1) |
| SASA Cutoff: | ΔSASA > 25.0 Å² | | | |

**Supplementary Table 2b: Comparative hydrophobic patch mapping**

| | CGMMV | | | | | ToMMV | | | | | |
|---|---|---|---|---|---|---|---|---|---|---|---|
| Patch ID | Patch Type | Involved Chains | Total ΔSASA (Å²) | Key Residues Involved | Number of residues involved | Patch Type | Involved Chains | Total ΔSASA (Å²) | Key Residues Involved | Number of residues involved | Δ Buried Area (ToMMV – CGMMV) |
| 1 | Inter subunit interface | b, c, d | 558.12 | Val94(b), Val98(b), Val97(b), Val94(c), Val97(c), Val98(c), Pro100(c), Pro103(c), Pro100(d), Pro103(d) | 10 | Inter subunit interface | b, c | 535.48 | Pro8(b), Phe11(b), Val12(b), Val22(c), Ile25(c), Phe68(c), Val130(c), Val134(c), Phe139(c) | 9 | -22.64 |
| 2 | Inter subunit interface | c, s | 508.49 | Ala51(c), Leu52(c), Pro75(s), Val76(s), Pro79(s), Val82(s), Leu85(s), Phe137(s) | 8 | Inter subunit interface | c, d | 535.46 | Pro8(c), Val12(c), Phe11(c), Val22(d), Ile25(d), Phe68(d), Val130(d), Val134(d), Phe139(d) | 9 | +26.97 |
| 3 | Inter subunit interface | M, c | 508.46 | Ala51(M), Leu52(M), Pro75(c), Val76(c), Pro79(c), Val82(c), Leu85(c), Phe137(c) | 8 | Single-chain (Intrachain) | c | 227.56 | Tyr73(c), Ala75(c), Val76(c) | 3 | NA |
| 4 | Single-chain (Intrachain) | c | 249.80 | Leu25(c), Phe27(c), Val29(c), Ala30(c), Ile130(c) | 5 | Inter subunit interface | b, c | 209.12 | Leu32(c), Pro79(b), Ala83(b), Leu85(b) | 4 | NA |
| 5 | Single-chain (Intrachain) | c | 224.53 | Pro8(c), Leu11(c), Ile12(c) | 3 | Inter subunit interface | c, d | 209.12 | Pro79(c), Ala83(c), Leu85(c), Leu32(d) | 4 | NA |

| | | | | | | | | | | | |
|---|---|---|---|---|---|---|---|---|---|---|---|
| 6 | Single-chain (Intrachain) | c | 132.96 | Leu109(c), Val112(c) | 2 | Single-chain (Intrachain) | c | 108.42 | Ala106(c), Leu109(c) | 2 | -24.54 |

**Supplementary Table 2c: Comparative summary of hydrophobic patches**

| Parameter / Metric | CGMMV | ToMMV | Difference (ToMMV – CGMMV) |
|---|---|---|---|
| **Candidate Interface Residues (ΔSASA > 25 Å²)** | 160 | 120 | -40 |
| **Total Identified Patches** | 6 | 6 | 0 |
| **Inter-subunit Interface Patches** | 3 | 4 | +1 |
| **Intrachain Patches (Chain c)** | 3 | 2 | -1 |
| **Total Residues in Patches** | 36 | 31 | -5 |
| **Mean Patch Buried Area (Å²)** | 363.73 | 304.19 | -59.54 |
| **Largest Patch Buried Area (Å²)** | 558.12 | 535.48 | -22.64 |

**Supplementary Table 3: Summary of ΔSASA (Å$^2$) and percentage burial (% buried) for acidic Glu and Asp residues**

| CGMMV | | | | | TMV | | | | | ToMMV | | | | |
|---|---|---|---|---|---|---|---|---|---|---|---|---|---|---|
| Residue | Monomer SASA ($SASA_m$) | Assembly SASA ($SASA_a$) | ΔSASA= $SASA_m$ - $SASA_a$ | % buried | Residue | Monomer SASA ($SASA_m$) | Assembly SASA ($SASA_a$) | ΔSASA= $SASA_m$ - $SASA_a$ | % buried | Residue | Monomer SASA ($SASA_m$) | Assembly SASA ($SASA_a$) | ΔSASA= $SASA_m$ - $SASA_a$ | % buried |
| Glu47 | 108.44 | 28.46 | 79.98 | 73.7 | Glu22 | 96.36 | 40.49 | 55.87 | 57.9 | Glu23 | 87.36 | 41.55 | 45.81 | 52.4 |
| Glu96 | 132.42 | 79.71 | 52.71 | 39.8 | Glu50 | 106 | 10.46 | 95.54 | 90.1 | Glu51 | 116.55 | 23.87 | 92.68 | 79.5 |
| Glu107 | 89.7 | 15.57 | 74.13 | 82.6 | Glu95 | 139.23 | 70.26 | 68.97 | 49.5 | Glu96 | 116.44 | 69.04 | 47.4 | 40.7 |
| Glu125 | 26.54 | 13.21 | 13.33 | 50.2 | Glu97 | 66.73 | 23.11 | 43.62 | 65.3 | Glu98 | 57.51 | 15.72 | 41.79 | 72.6 |
| Glu131 | 96.2 | 20.36 | 75.84 | 78.8 | Glu106 | 63.07 | 5.7 | 57.37 | 90.9 | Glu107 | 78.14 | 10.83 | 67.31 | 86.1 |
| Glu146 | 33.76 | 33.76 | 0 | 0 | Glu131 | 45.2 | 9.91 | 35.29 | 78 | Glu132 | 44.82 | 20.32 | 24.5 | 54.6 |
| Glu155 | 140.29 | 140.29 | 0 | 0 | Glu145 | 45.06 | 45.06 | 0 | 0 | Glu146 | 39.22 | 39.22 | 0 | 0 |
| Asp43 | 24.44 | 24.44 | 0 | 0 | Asp19 | 55.74 | 58.74 | -3 | -5.3 | Asp20 | 65.25 | 65.25 | 0 | 0 |
| Asp58 | 72.54 | 37.06 | 35.48 | 48.9 | Asp64 | 99.43 | 99.43 | 0 | 0 | Asp65 | 92.32 | 92.32 | 0 | 0 |
| Asp65 | 112.88 | 112.88 | 0 | 0 | Asp66 | 67.39 | 63.01 | 4.38 | 6.4 | Asp78 | 38.09 | 8.12 | 29.97 | 78.6 |
| Asp89 | 87.65 | 3.85 | 83.8 | 95.6 | Asp77 | 44.32 | 5.33 | 38.99 | 87.9 | Asp89 | 115.38 | 10 | 105.38 | 91.3 |
| Asp99 | 77.28 | 45.47 | 31.81 | 41.1 | Asp88 | 111.2 | 5.03 | 106.17 | 95.4 | Asp110 | 98.64 | 62.73 | 35.91 | 36.4 |
| Asp116 | 61.24 | 36.64 | 24.6 | 40.1 | Asp109 | 82.19 | 34.26 | 47.93 | 58.3 | Asp116 | 66.78 | 47.27 | 19.51 | 29.2 |
| Asp117 | 84.55 | 53.29 | 31.26 | 36.9 | Asp115 | 57 | 33.65 | 23.35 | 40.9 | Asp117 | 104.96 | 75.52 | 29.44 | 28 |
| Asp127 | 77.35 | 0.37 | 76.98 | 99.5 | Asp116 | 79.41 | 55.02 | 24.39 | 30.7 | | | | | |
| Asp138 | 15.86 | 15.86 | 0 | 0 | | | | | | | | | | |
| Asp141 | 39.92 | 39.25 | 0.67 | 1.6 | | | | | | | | | | |

**Supplementary Table 4: Amino acid conservation scores calculated in ConSurf based on Multiple Sequence Alignment (MSA) of *Tobamovirus* coat protein and TMV coat protein pdb (PDB 4UDV)**

POS: The position of the amino acid in the sequence.
- SEQ: The one-letter amino acid code.
- ATOM: When there's a model, the ATOM-derived sequence in three-letter code, including the acid's positions as they appear in the PDB file and the chain identifier.
- SCORE: The normalized conservation scores.
- COLOR: The color scale representing the conservation scores (9 - conserved, 1 - variable).
- CONFIDENCE INTERVAL: When using the Bayesian method for calculating rates, a confidence interval is assigned to each of the inferred evolutionary conservation scores; next to it are the colors of the lower and upper bounds of the confidence interval
- B/E: Buried (b) or Exposed (e) residue.
- F/S: functional (f) or structural (s) residue (f - highly conserved and exposed, s - highly conserved and buried).
- MSA DATA: The number of aligned sequences having an amino acid (non-gapped) from the overall number of sequences at each position.
- RESIDUE VARIETY: The residue variety at each position of the multiple sequence alignment.

| POS | SEQ | ATOM | SCORE | COLOR | CONFIDENCE INTERVAL | B/E | F/S | MSA DATA | RESIDUE VARIETY |
|---|---|---|---|---|---|---|---|---|---|
| 1 | M | - | -1.370 | 9 | -1.488, -1.331 9,9 | e | f | 32/33 | M |
| 2 | S | SER:1:A | 0.972 | 4 | 0.269, 1.423 5,3 | e | | 32/33 | S 50%, P 25%, A 21%, V 3% |
| 3 | Y | TYR:2:A | -1.262 | 9 | -1.447, -1.177 9,9 | e | f | 32/33 | Y |
| 4 | S | SER:3:A | 0.399 | 4 | -0.094, 0.808 5,4 | e | | 32/33 | T 40%, S 21%, N 15%, Q 6%, A 6%, H 3%, L 3%, I 3% |
| 5 | I | ILE:4:A | 0.090 | 5 | -0.372, 0.422 6,4 | e | | 32/33 | I 43%, N 18%, V 12%, T 12%, P 12% |
| 6 | T | THR:5:A | 0.810 | 4 | 0.134, 1.072 5,3 | e | | 32/33 | T 34%, S 18%, I 18%, V 9%, N 6%, A 6%, P 3%, L 3% |
| 7 | T | THR:6:A | 0.644 | 4 | 0.134, 1.072 5,3 | e | | 32/33 | S 40%, T 25%, G 9%, N 9%, A 3%, K 3%, R 3%, D 3%, Q 3% |
| 8 | P | PRO:7:A | 1.317 | 3 | 0.422, 1.959 4,2 | e | | 32/33 | P 62%, I 9%, S 9%, L 6%, T 6%, A 6% |
| 9 | S | SER:8:A | 0.854 | 4 | 0.269, 1.423 5,3 | e | | 32/33 | S 31%, N 31%, R 9%, K 9%, T 6%, L 6%, Q 3%, V 3% |
| 10 | Q | GLN:9:A | -0.057 | 5 | -0.452, 0.269 6,5 | e | | 32/33 | Q 59%, N 15%, S 9%, T 6%, K 6%, D 3% |
| 11 | F | PHE:10:A | 0.121 | 5 | -0.372, 0.422 6,4 | b | | 32/33 | L 53%, F 28%, Y 12%, I 6% |
| 12 | V | VAL:11:A | 0.530 | 4 | -0.094, 0.808 5,4 | b | | 32/33 | V 46%, P 15%, Q 15%, I 12%, A 6%, S 3% |
| 13 | Y | PHE:12:A | -0.062 | 5 | -0.527, 0.269 7,5 | b | | 32/33 | Y 65%, A 12%, F 12%, W 6%, L 3% |
| 14 | L | LEU:13:A | 0.783 | 4 | 0.134, 1.072 5,3 | b | | 32/33 | L 46%, F 37%, R 3%, Y 3%, S 3%, T 3%, V 3% |
| 15 | S | SER:14:A | -0.048 | 5 | -0.452, 0.269 6,5 | b | | 32/33 | S 50%, A 21%, T 15%, G 12% |
| 16 | S | SER:15:A | 0.626 | 4 | 0.014, 1.072 5,3 | e | | 32/33 | S 40%, A 21%, K 9%, R 9%, N 6%, V 3%, Q 3%, P 3%, G 3% |
| 17 | A | ALA:16:A | -0.084 | 5 | -0.452, 0.134 6,5 | b | | 32/33 | A 31%, S 25%, V 25%, N 9%, T 6%, M 3% |
| 18 | W | TRP:17:A | 0.418 | 4 | -0.194, 0.808 6,4 | b | | 32/33 | W 59%, Y 21%, F 18% |
| 19 | A | ALA:18:A | -0.457 | 7 | -0.793, -0.194 8,6 | b | | 32/33 | A 71%, V 12%, I 6%, C 3%, Y 3%, F 3% |
| 20 | D | ASP:19:A | -0.116 | 5 | -0.599, 0.269 7,5 | e | | 32/33 | D 53%, P 28%, E 18% |
| 21 | P | PRO:20:A | 0.174 | 5 | -0.372, 0.598 6,4 | b | | 32/33 | P 62%, Y 21%, F 6%, G 6%, V 3% |

| | | | | | | | | | |
|---|---|---|---|---|---|---|---|---|---|
| 22 | I | ILE:21:A | 3.145 | 1 | 1.959, 3.352 2,1 | b | | 32/33 | I 28%, T 15%, V 15%, Y 6%, D 6%, Q 6%, L 6%, E 3%, F 3%, R 3%, N 3%, K 3% |
| 23 | E | GLU:22:A | 2.940 | 1 | 1.423, 3.352 3,1 | e | | 32/33 | E 31%, A 15%, T 12%, D 9%, V 6%, P 6%, L 3%, K 3%, G 3%, S 3%, R 3%, Q 3% |
| 24 | L | LEU:23:A | -0.051 | 5 | -0.527, 0.269 7,5 | b | | 33/33 | L 63%, F 18%, M 15%, V 3% |
| 25 | I | ILE:24:A | 1.334 | 3 | 0.598, 1.959 4,2 | b | | 33/33 | L 36%, I 36%, Y 12%, M 6%, V 6%, Q 3% |
| 26 | N | ASN:25:A | -0.432 | 6 | -0.793, -0.194 8,6 | e | | 33/33 | N 72%, D 12%, E 6%, S 6%, Q 3% |
| 27 | L | LEU:26:A | 0.784 | 4 | 0.134, 1.072 5,3 | b | | 33/33 | L 54%, Q 12%, I 9%, V 9%, Y 6%, F 6%, S 3% |
| 28 | C | CYS:27:A | -0.975 | 8 | -1.228, -0.793 9,8 | b | | 33/33 | C 57%, L 36%, I 6% |
| 29 | T | THR:28:A | 0.554 | 4 | 0.014, 0.808 5,4 | b | | 33/33 | T 42%, V 21%, I 15%, R 6%, S 6%, Q 3%, M 3%, N 3% |
| 30 | N | ASN:29:A | 0.869 | 4 | 0.269, 1.423 5,3 | e | | 33/33 | N 30%, S 30%, T 9%, K 9%, A 9%, V 6%, Q 3%, D 3% |
| 31 | A | ALA:30:A | 0.340 | 5 | -0.194, 0.598 6,4 | b | | 33/33 | A 48%, S 33%, G 6%, T 6%, M 3%, N 3% |
| 32 | L | LEU:31:A | -0.564 | 7 | -0.911, -0.372 8,6 | e | | 33/33 | L 60%, Q 24%, R 9%, S 6% |
| 33 | G | GLY:32:A | 0.091 | 5 | -0.372, 0.422 6,4 | e | | 33/33 | G 63%, S 21%, A 6%, L 3%, V 3%, T 3% |
| 34 | N | ASN:33:A | -0.107 | 5 | -0.527, 0.134 7,5 | e | | 33/33 | N 51%, Q 24%, T 9%, G 6%, L 3%, E 3%, A 3% |
| 35 | Q | GLN:34:A | -0.615 | 7 | -0.911, -0.452 8,6 | e | | 33/33 | Q 42%, S 33%, A 18%, I 6% |
| 36 | F | PHE:35:A | -0.937 | 8 | -1.228, -0.793 9,8 | b | | 33/33 | F 81%, L 9%, Y 9% |
| 37 | Q | GLN:36:A | -1.351 | 9 | -1.488, -1.279 9,9 | e | f | 33/33 | Q |
| 38 | T | THR:37:A | -1.239 | 9 | -1.447, -1.177 9,9 | e | f | 33/33 | T 96%, A 3% |
| 39 | Q | GLN:38:A | -1.351 | 9 | -1.488, -1.279 9,9 | e | f | 33/33 | Q |
| 40 | Q | GLN:39:A | -0.511 | 7 | -0.853, -0.285 8,6 | e | | 33/33 | Q 39%, A 33%, N 21%, S 6% |
| 41 | A | ALA:40:A | -0.816 | 8 | -1.126, -0.666 9,7 | b | | 33/33 | A 63%, G 33%, S 3% |
| 42 | R | ARG:41:A | -1.162 | 9 | -1.385, -1.074 9,9 | e | f | 33/33 | R 90%, K 9% |
| 43 | T | THR:42:A | -1.010 | 8 | -1.279, -0.853 9,8 | e | f | 33/33 | D 51%, T 48% |
| 44 | V | VAL:43:A | 0.006 | 5 | -0.452, 0.269 6,5 | e | | 33/33 | T 51%, I 12%, S 12%, Q 9%, E 6%, V 6%, A 3% |
| 45 | V | VAL:44:A | -0.532 | 7 | -0.911, -0.285 8,6 | b | | 33/33 | V 60%, L 24%, F 9%, I 6% |
| 46 | Q | GLN:45:A | -0.841 | 8 | -1.126, -0.666 9,7 | e | f | 33/33 | R 51%, Q 42%, A 3%, L 3% |
| 47 | R | ARG:46:A | 0.053 | 5 | -0.372, 0.422 6,4 | e | | 33/33 | Q 45%, E 21%, S 9%, R 9%, N 6%, D 6%, A 3% |
| 48 | Q | GLN:47:A | -0.988 | 8 | -1.228, -0.853 9,8 | e | f | 33/33 | Q 75%, S 15%, E 6%, V 3% |
| 49 | F | PHE:48:A | -0.950 | 8 | -1.228, -0.793 9,8 | b | | 33/33 | F 57%, L 39%, I 3% |
| 50 | S | SER:49:A | 1.029 | 4 | 0.422, 1.423 4,3 | e | | 33/33 | S 51%, A 12%, V 9%, I 9%, N 6%, T 6%, Q 3%, G 3% |
| 51 | E | GLU:50:A | 0.078 | 5 | -0.372, 0.422 6,4 | e | | 33/33 | D 27%, N 24%, G 24%, E 15%, A 6%, T 3% |
| 52 | V | VAL:51:A | 1.074 | 3 | 0.422, 1.423 4,3 | b | | 33/33 | L 30%, V 27%, S 12%, I 6%, A 6%, C 6%, Q 3%, F 3%, M 3%, T 3% |
| 53 | W | TRP:52:A | 0.804 | 4 | 0.134, 1.423 5,3 | b | | 33/33 | W 36%, L 33%, F 9%, V 9%, Y 6%, Q 3%, P 3% |
| 54 | K | LYS:53:A | 0.657 | 4 | 0.014, 1.072 5,3 | e | | 33/33 | K 45%, S 24%, Q 9%, R 6%, V 6%, T 6%, D 3% |
| 55 | P | PRO:54:A | 0.897 | 4 | 0.134, 1.423 5,3 | e | | 33/33 | P 24%, S 21%, A 15%, T 15%, Y 6%, V 6%, I 6%, E 6% |
| 56 | S | SER:55:A | 0.513 | 4 | -0.094, 0.808 5,4 | b | | 33/33 | V 54%, I 12%, G 6%, L 6%, S 6%, F 6%, T 3%, P 3%, A 3% |
| 57 | P | PRO:56:A | -0.531 | 7 | -0.911, -0.285 8,6 | e | | 33/33 | P 48%, V 36%, A 15% |
| 58 | Q | GLN:57:A | 0.352 | 5 | -0.094, 0.598 5,4 | e | | 33/33 | T 36%, Q 24%, S 21%, A 9%, R 3%, D 3%, N 3% |
| 59 | V | VAL:58:A | 3.189 | 1 | 1.959, 3.352 2,1 | e | | 33/33 | P 27%, V 18%, L 9%, S 9%, A 6%, K 6%, R 6%, Q 6%, N 3%, I 3%, M 3%, T 3% |
| 60 | T | THR:59:A | 0.621 | 4 | 0.014, 1.072 5,3 | e | | 33/33 | T 51%, N 30%, S 6%, G 3%, D 3%, R 3%, I 3% |
| 61 | V | VAL:60:A | 0.909 | 4 | 0.269, 1.423 5,3 | b | | 33/33 | V 45%, S 18%, Q 12%, I 9%, T 9%, N 3%, D 3% |
| 62 | R | ARG:61:A | -1.161 | 9 | -1.385, -1.074 9,9 | e | f | 33/33 | R 96%, Q 3% |

| | | | | | | | | | |
|---|---|---|---|---|---|---|---|---|---|
| 63 | F | PHE:62:A | -1.134 | 9 | -1.385, -1.020 9,8 | b | s | 33/33 | F 96%, Y 3% |
| 64 | P | PRO:63:A | -0.656 | 7 | -1.020, -0.372 8,6 | e | | 33/33 | P 93%, N 3%, S 3% |
| 65 | D | ASP:64:A | 2.685 | 1 | 1.423, 3.352 3,1 | e | | 32/33 | D 28%, A 25%, E 25%, T 3%, L 3%, I 3%, G 3%, N 3%, V 3%, S 3% |
| 66 | S | SER:65:A | 2.774 | 1 | 1.423, 3.352 3,1 | e | | 32/33 | N 18%, G 18%, S 12%, D 12%, A 12%, T 12%, R 6%, E 3%, L 3% |
| 67 | D | ASP:66:A | 3.002 | 1 | 1.959, 3.352 2,1 | e | | 31/33 | G 38%, E 12%, V 12%, T 9%, D 9%, S 6%, R 3%, A 3%, I 3% |
| 68 | F | PHE:67:A | 0.155 | 5 | -0.372, 0.422 6,4 | b | | 33/33 | F 69%, Y 18%, I 6%, W 3%, S 3% |
| 69 | K | LYS:68:A | 0.840 | 4 | 0.134, 1.423 5,3 | e | | 33/33 | Y 27%, K 24%, R 24%, L 15%, F 6%, Q 3% |
| 70 | V | VAL:69:A | -1.011 | 8 | -1.228, -0.853 9,8 | b | | 33/33 | V 87%, L 6%, A 3%, I 3% |
| 71 | Y | TYR:70:A | 0.738 | 4 | 0.134, 1.072 5,3 | b | | 33/33 | Y 51%, W 21%, F 18%, N 6%, S 3% |
| 72 | R | ARG:71:A | 1.034 | 4 | 0.269, 1.423 5,3 | e | | 33/33 | R 42%, V 15%, S 12%, I 12%, L 6%, A 6%, T 3%, F 3% |
| 73 | Y | TYR:72:A | 0.129 | 5 | -0.372, 0.422 6,4 | e | | 33/33 | Y 39%, N 30%, R 15%, A 3%, V 3%, G 3%, T 3%, F 3% |
| 74 | N | ASN:73:A | 0.290 | 5 | -0.194, 0.598 6,4 | e | | 33/33 | N 51%, D 15%, E 12%, S 12%, V 3%, G 3%, K 3% |
| 75 | A | ALA:74:A | 0.048 | 5 | -0.452, 0.422 6,4 | e | | 33/33 | A 45%, P 24%, S 21%, G 6%, L 3% |
| 76 | V | VAL:75:A | 0.574 | 4 | 0.014, 0.808 5,4 | b | | 33/33 | V 45%, T 24%, R 9%, A 9%, S 6%, I 3%, G 3% |
| 77 | L | LEU:76:A | 0.074 | 5 | -0.372, 0.422 6,4 | b | | 33/33 | L 63%, I 33%, V 3% |
| 78 | D | ASP:77:A | 0.741 | 4 | 0.134, 1.072 5,3 | e | | 33/33 | D 42%, A 18%, K 15%, S 9%, R 9%, G 3%, E 3% |
| 79 | P | PRO:78:A | 0.310 | 5 | -0.285, 0.808 6,4 | e | | 33/33 | P 69%, A 12%, D 6%, S 6%, Q 3%, T 3% |
| 80 | L | LEU:79:A | -0.240 | 6 | -0.666, 0.014 7,5 | b | | 33/33 | L 69%, V 15%, I 9%, A 3%, Y 3% |
| 81 | V | VAL:80:A | 0.834 | 4 | 0.269, 1.423 5,3 | b | | 33/33 | I 33%, V 15%, W 15%, Y 15%, F 12%, L 9% |
| 82 | T | THR:81:A | 1.516 | 3 | 0.598, 1.959 4,2 | e | | 33/33 | T 36%, E 21%, F 9%, D 9%, S 6%, N 3%, V 3%, K 3%, H 3%, G 3%, L 3% |
| 83 | A | ALA:82:A | -0.278 | 6 | -0.666, 0.014 7,5 | b | | 33/33 | A 69%, S 21%, F 3%, G 3%, N 3% |
| 84 | L | LEU:83:A | -0.937 | 8 | -1.228, -0.793 9,8 | b | | 33/33 | L 90%, V 9% |
| 85 | L | LEU:84:A | 0.002 | 5 | -0.452, 0.269 6,5 | b | | 33/33 | L 66%, M 18%, F 9%, A 3%, T 3% |
| 86 | G | GLY:85:A | -0.375 | 6 | -0.731, -0.094 7,5 | e | | 33/33 | G 30%, Q 21%, K 21%, N 15%, S 9%, R 3% |
| 87 | A | ALA:86:A | -0.265 | 6 | -0.599, 0.014 7,5 | b | | 33/33 | S 45%, A 45%, T 9% |
| 88 | F | PHE:87:A | -0.389 | 6 | -0.793, -0.094 8,5 | b | | 33/33 | F 57%, T 18%, L 18%, C 3%, A 3% |
| 89 | D | ASP:88:A | -1.299 | 9 | -1.488, -1.228 9,9 | e | f | 33/33 | D |
| 90 | T | THR:89:A | -1.248 | 9 | -1.447, -1.177 9,9 | e | f | 33/33 | T 90%, S 9% |
| 91 | R | ARG:90:A | -0.556 | 7 | -0.911, -0.372 8,6 | e | | 33/33 | R 75%, K 24% |
| 92 | N | ASN:91:A | -1.229 | 9 | -1.385, -1.126 9,9 | e | f | 33/33 | N 93%, D 6% |
| 93 | R | ARG:92:A | -1.303 | 9 | -1.488, -1.228 9,9 | b | s | 33/33 | R |
| 94 | I | ILE:93:A | -0.919 | 8 | -1.177, -0.793 9,8 | b | | 33/33 | I 78%, V 12%, A 9% |
| 95 | I | ILE:94:A | -1.136 | 9 | -1.331, -1.020 9,8 | b | s | 33/33 | I 90%, F 6%, L 3% |
| 96 | E | GLU:95:A | -1.295 | 9 | -1.488, -1.228 9,9 | e | f | 33/33 | E |
| 97 | V | VAL:96:A | -0.913 | 8 | -1.177, -0.793 9,8 | b | | 33/33 | V 75%, T 24% |
| 98 | E | GLU:97:A | 0.235 | 5 | -0.285, 0.598 6,4 | e | | 33/33 | E 72%, D 15%, G 3%, V 3%, R 3%, S 3% |
| 99 | N | ASN:98:A | -0.472 | 7 | -0.793, -0.285 8,6 | e | | 33/33 | N 66%, E 24%, D 6%, T 3% |
| 100 | Q | GLN:99:A | 1.380 | 3 | 0.598, 1.959 4,2 | e | | 33/33 | P 36%, E 24%, Q 21%, A 6%, N 3%, S 3%, V 3%, D 3% |
| 101 | A | ALA:100:A | 0.021 | 5 | -0.372, 0.269 6,5 | e | | 33/33 | S 36%, Q 18%, A 18%, R 12%, P 6%, V 3%, I 3%, T 3% |
| 102 | N | ASN:101:A | -0.147 | 5 | -0.527, 0.134 7,5 | e | | 33/33 | N 57%, R 21%, S 12%, A 6%, G 3% |

| | | | | | | | | | | |
|---|---|---|---|---|---|---|---|---|---|---|
| 103 | P | PRO:102:A | -0.659 | 7 | -1.074, -0.452 | 9,6 | e | | 33/33 | P 90%, V 9% |
| 104 | T | THR:103:A | -0.568 | 7 | -0.853, -0.372 | 8,6 | e | | 33/33 | T 54%, S 36%, N 9% |
| 105 | T | THR:104:A | -0.641 | 7 | -0.911, -0.452 | 8,6 | e | | 33/33 | T 66%, N 12%, A 12%, S 9% |
| 106 | A | ALA:105:A | 0.451 | 4 | -0.094, 0.808 | 5,4 | e | | 33/33 | A 48%, S 27%, T 12%, G 9%, N 3% |
| 107 | E | GLU:106:A | -1.295 | 9 | -1.488, -1.228 | 9,9 | e | f | 33/33 | E |
| 108 | T | THR:107:A | -0.145 | 5 | -0.527, 0.134 | 7,5 | b | | 33/33 | T 42%, V 21%, I 15%, A 9%, S 9%, Q 3% |
| 109 | L | LEU:108:A | 0.650 | 4 | 0.014, 1.072 | 5,3 | e | | 33/33 | L 51%, A 18%, K 6%, R 6%, E 6%, V 6%, M 3%, I 3% |
| 110 | D | ASP:109:A | -0.439 | 6 | -0.793, -0.194 | 8,6 | e | | 33/33 | N 45%, D 27%, S 15%, L 6%, E 6% |
| 111 | A | ALA:110:A | -1.219 | 9 | -1.385, -1.126 | 9,9 | b | s | 33/33 | A 93%, V 6% |
| 112 | T | THR:111:A | -0.954 | 8 | -1.177, -0.793 | 9,8 | b | | 33/33 | T 87%, V 12% |
| 113 | R | ARG:112:A | -0.652 | 7 | -0.966, -0.452 | 8,6 | e | | 33/33 | Q 45%, R 42%, K 12% |
| 114 | R | ARG:113:A | -1.303 | 9 | -1.488, -1.228 | 9,9 | e | f | 33/33 | R |
| 115 | V | VAL:114:A | -1.021 | 8 | -1.228, -0.911 | 9,8 | b | | 33/33 | V 81%, T 9%, N 9% |
| 116 | D | ASP:115:A | -1.299 | 9 | -1.488, -1.228 | 9,9 | e | f | 33/33 | D |
| 117 | D | ASP:116:A | -1.299 | 9 | -1.488, -1.228 | 9,9 | e | f | 33/33 | D |
| 118 | A | ALA:117:A | -1.092 | 9 | -1.331, -0.966 | 9,8 | b | s | 33/33 | A 93%, S 6% |
| 119 | T | THR:118:A | -1.001 | 8 | -1.228, -0.853 | 9,8 | b | | 33/33 | T 78%, S 21% |
| 120 | V | VAL:119:A | -0.864 | 8 | -1.126, -0.731 | 9,7 | b | | 33/33 | V 78%, T 18%, I 3% |
| 121 | A | ALA:120:A | -0.963 | 8 | -1.177, -0.793 | 9,8 | b | | 33/33 | A 84%, N 9%, S 6% |
| 122 | I | ILE:121:A | -1.010 | 8 | -1.228, -0.853 | 9,8 | b | | 33/33 | I 78%, A 12%, V 6%, S 3% |
| 123 | R | ARG:122:A | -0.989 | 8 | -1.228, -0.853 | 9,8 | b | | 33/33 | R 81%, H 18% |
| 124 | S | SER:123:A | 0.352 | 5 | -0.094, 0.598 | 5,4 | b | | 33/33 | S 45%, A 18%, N 18%, K 6%, I 6%, V 3%, G 3% |
| 125 | A | ALA:124:A | 0.390 | 4 | -0.094, 0.808 | 5,4 | b | | 33/33 | A 30%, Q 18%, S 15%, N 9%, D 9%, E 9%, C 6%, G 3% |
| 126 | I | ILE:125:A | -0.676 | 7 | -0.966, -0.527 | 8,7 | b | | 33/33 | I 75%, L 15%, V 9% |
| 127 | N | ASN:126:A | 0.441 | 4 | -0.094, 0.808 | 5,4 | e | | 33/33 | N 36%, Q 21%, D 12%, P 9%, E 6%, T 3%, R 3%, I 3%, S 3%, K 3% |
| 128 | N | ASN:127:A | 0.125 | 5 | -0.372, 0.422 | 6,4 | e | | 33/33 | N 54%, L 21%, Q 9%, R 9%, T 3%, S 3% |
| 129 | L | LEU:128:A | -0.552 | 7 | -0.911, -0.285 | 8,6 | b | | 33/33 | L 84%, I 9%, C 3%, T 3% |
| 130 | V | ILE:129:A | 1.049 | 4 | 0.422, 1.423 | 4,3 | b | | 33/33 | L 39%, V 18%, M 15%, A 6%, I 6%, F 6%, R 3%, S 3%, C 3% |
| 131 | V | VAL:130:A | 0.693 | 4 | 0.134, 1.072 | 5,3 | e | | 33/33 | N 45%, S 15%, D 9%, V 9%, A 6%, K 6%, E 3%, G 3%, L 3% |
| 132 | E | GLU:131:A | -0.114 | 5 | -0.599, 0.134 | 7,5 | e | | 33/33 | E 57%, A 21%, L 12%, I 3%, S 3%, Y 3% |
| 133 | L | LEU:132:A | -0.970 | 8 | -1.228, -0.793 | 9,8 | b | | 33/33 | L 93%, I 6% |
| 134 | I | ILE:133:A | 0.615 | 4 | 0.134, 1.072 | 5,3 | b | | 33/33 | V 33%, S 21%, N 15%, I 12%, A 6%, Q 6%, T 3%, G 3% |
| 135 | R | ARG:134:A | 1.638 | 3 | 0.808, 1.959 | 4,2 | e | | 33/33 | R 42%, G 12%, D 9%, N 9%, S 6%, A 6%, K 6%, E 6%, Q 3% |
| 136 | G | GLY:135:A | -0.672 | 7 | -1.074, -0.452 | 9,6 | b | | 33/33 | G 90%, V 6%, A 3% |
| 137 | T | THR:136:A | 0.555 | 4 | 0.014, 0.808 | 5,4 | e | | 33/33 | T 51%, H 15%, V 6%, A 6%, N 6%, S 6%, F 3%, Q 3%, E 3% |
| 138 | G | GLY:137:A | -0.548 | 7 | -0.966, -0.285 | 8,6 | e | | 33/33 | G 84%, D 9%, A 6% |
| 139 | S | SER:138:A | 0.091 | 5 | -0.372, 0.422 | 6,4 | b | | 33/33 | V 30%, M 21%, L 18%, F 9%, Y 9%, I 6%, S 6% |
| 140 | Y | TYR:139:A | 0.127 | 5 | -0.372, 0.422 | 6,4 | b | | 33/33 | Y 54%, M 18%, L 12%, F 12%, V 3% |
| 141 | N | ASN:140:A | -0.733 | 7 | -1.020, -0.527 | 8,7 | e | | 33/33 | N 63%, D 33%, S 3% |
| 142 | R | ARG:141:A | -0.433 | 6 | -0.793, -0.194 | 8,6 | e | | 33/33 | R 60%, Q 30%, T 6%, S 3% |
| 143 | S | SER:142:A | 0.543 | 4 | 0.014, 0.808 | 5,4 | b | | 33/33 | A 48%, T 18%, S 18%, K 6%, Q 3%, V 3%, N 3% |
| 144 | S | SER:143:A | 0.627 | 4 | 0.014, 1.072 | 5,3 | e | | 33/33 | S 45%, T 15%, E 12%, A 9%, L 9%, I 3%, G 3%, Q 3% |
| 145 | F | PHE:144:A | -1.290 | 9 | -1.488, -1.228 | 9,9 | b | s | 33/33 | F |
| 146 | E | GLU:145:A | -1.295 | 9 | -1.488, -1.228 | 9,9 | e | f | 33/33 | E |

| | | | | | | | | | | |
|---|---|---|---|---|---|---|---|---|---|---|
| 147 | S | SER:146:A | 0.979 | 4 | 0.269, 1.423 | 5,3 | e | | 33/33 | S 42%, T 15%, A 12%, V 9%, D 6%, Q 6%, E 6%, I 3% |
| 148 | S | SER:147:A | 0.747 | 4 | 0.134, 1.072 | 5,3 | b | | 33/33 | A 36%, V 18%, M 12%, S 9%, L 9%, T 6%, I 6%, Q 3% |
| 149 | S | SER:148:A | -0.672 | 7 | -0.966, -0.452 | 8,6 | b | | 28/33 | S 71%, F 14%, A 3%, T 3%, N 3%, L 3% |
| 150 | G | GLY:149:A | -0.441 | 6 | -0.853, -0.194 | 8,6 | e | | 28/33 | G 85%, N 7%, S 3%, Q 3% |
| 151 | L | LEU:150:A | -0.507 | 7 | -0.911, -0.285 | 8,6 | b | | 33/33 | L 84%, W 9%, V 3%, I 3% |
| 152 | V | VAL:151:A | 0.583 | 4 | 0.014, 1.072 | 5,3 | b | | 33/33 | T 42%, V 27%, P 12%, S 3%, R 3%, I 3%, L 3%, Q 3%, A 3% |
| 153 | W | TRP:152:A | -1.172 | 9 | -1.447, -1.074 | 9,9 | b | s | 33/33 | W |
| 154 | T | - | 0.512 | 4 | 0.014, 0.808 | 5,4 | e | | 33/33 | T 51%, A 21%, S 12%, E 6%, H 3%, V 3%, Q 3% |
| 155 | S | - | 1.080 | 3 | 0.422, 1.423 | 4,3 | e | | 33/33 | T 36%, A 21%, S 18%, E 15%, D 6%, L 3% |
| 156 | G | - | 1.736 | 3 | 0.808, 1.959 | 4,2 | e | | 33/33 | A 45%, T 15%, S 9%, G 9%, R 6%, P 6%, Q 3%, D 3%, N 3% |
| 157 | P | - | 2.462 | 2 | 1.423, 3.352 | 3,1 | e | | 32/33 | P 40%, A 18%, T 15%, G 9%, S 9%, C 3%, L 3% |
| 158 | A | - | 0.584 | 4 | 0.014, 1.072 | 5,3 | e | | 31/33 | A 48%, T 25%, S 9%, P 6%, H 3%, N 3%, Q 3% |
| 159 | T | THR:153:A | 0.483 | 4 | -0.094, 0.808 | 5,4 | e | | 29/33 | T 44%, S 31%, V 6%, P 6%, K 6%, L 3% |

**Supplementary Table 5: Cryo-EM Data collection, refinement, and validation statistics**

| Parameter / Metric | CGMMV | ToMMV |
|---|---|---|
| **Data Collection & Processing** | | |
| **Electron microscope** | ThermoFisher Scientific Titan Krios | ThermoFisher Scientific Krios (G3i) |
| **Voltage (kV)** | 300 | 300 |
| **Detector** | Falcon III direct detector | Gatan K3 direct detector |
| **Pixel size (Å/pixel)** | 1.38 | 1.1 |
| **Defocus range (µm)** | -1.8 to -3.2 | -0.6 to -2.2 |
| **Magnification** | 59000 | 81000 |
| **Total electron exposure ($e^-$/Å²)** | 35.82 | 39.3 |
| **Micrographs collected (n)** | 1132 | 4884 |
| **Initial extracted helical segments (n)** | 2,137,255 | 15,765,310 |
| **Final selected segments for 3D refinement (n)** | 2,102,400 | 15,132,861 |
| **Symmetry imposed** | Helical | Helical |
| **Helical symmetry search range** | Twist 19°-25°, Rise 1.26-1.54 Å | Twist 19°-25°, Rise 1.26-1.54 Å |
| **Determination software** | RELION / CryoSPARC / PyHI | RELION / CryoSPARC / PyHI |
| **Map Reconstruction & Resolution** | | |
| **Overall map resolution, FSC = 0.143 (Å)** | **2.86** | **2.71** |
| **Unmasked FSC resolution (Å)** | 3.10 | 2.90 |
| **Tight masked FSC resolution (Å)** | 2.80 | 2.70 |
| **Corrected FSC resolution (Å)** | 2.90 | 2.70 |
| **Local resolution range (Å)** | 2.6 - 6.2 | 2.2 - 4.5 |
| **Median local resolution (Å)** | 2.90 | 2.30 |
| **Mean local resolution (Å)** | 3.12 | 2.45 |

| | | |
|---|---|---|
| **Map sharpening *B* factor (Å$^2$)** | -150.3 | -133.4 |
| **Refinement & Model Validation** | | |
| **Model composition** | | |
| **Protein residues (per coat protein subunit)** | 155 | 151 |
| **RNA nucleotides (per coat protein subunit)** | 3 | 3 |
| **RMS deviations** | | |
| **Bond lengths (Å)** | 0.004 | 0.004 |
| **Bond angles (°)** | 0.718 | 0.549 |
| **Ramachandran plot (%)** | | |
| **Favored** | 99 | 97 |
| **Allowed** | 1 | 3 |
| **Outliers** | 0.0 | 0.0 |
| **Rotamer outliers (%)** | 2 | 2 |
| **All-atom Clashscore** | 5 | 1.27 |
| **MolProbity overall score** | 1.41 | 1.20 |
| **Map-model overall mean Q Score** | 0.74 | 0.79 |
| **Accession Codes** | | |
| **PDB ID** | 9VJD | 9WQQ |
| **EMDB ID** | EMD-60696 | EMD-63082 |

**Supplementary Table 6: List of Tobamoviruses from ICTV taken for the phylogenetic study.**
Sequence acquisition date: 17th September, 2025.

| Serial No. | Species | Virus name | Isolate | Abbrev | Accession no. for coat protein | Host family | Amino acid composition at 113th position |
|---|---|---|---|---|---|---|---|
| 1 | *Tobamovirus maculacapsici* | Bell pepper mottle virus | Netherlands | BPMV | ABC87814.1 | Solanaceae | R |
| 2 | *Tobamovirus brugmansiae* | Brugmansia mild mottle virus | 2373 | BrMMV | CAL39166.1 | Solanaceae | Q |
| 3 | *Tobamovirus cacti* | Cactus mild mottle virus | Kr | CMMoV | ABX57123.1 | Cactaceae | Q |
| 4 | *Tobamovirus clitoriae* | Clitoria yellow mottle virus | Larrimah | CYMV | AEW50167.1 | Fabaceae | R |
| 5 | *Tobamovirus maculafructi* | Cucumber fruit mottle mosaic virus | Wang | CFMMV | AAG42817.1 | Cucurbitaceae | K |
| 6 | *Tobamovirus viridimaculae* | Cucumber green mottle mosaic virus | BgDel | CGMMV | AAP74202.1 | Cucurbitaceae | K |
| 7 | *Tobamovirus cucumeris* | Cucumber mottle virus | Kubota | CMoV | BAF37647.1 | Cucurbitaceae | Q |
| 8 | *Tobamovirus frangipani* | Frangipani mosaic virus | P | FrMV | ADN93256.1 | Apocynaceae | R |
| 9 | *Tobamovirus fortpiercense* | Hibiscus latent Fort Pierce virus | J | HLFPV | BAP76309.1 | Malvaceae | Q |
| 10 | *Tobamovirus singaporense* | Hibiscus latent Singapore virus | Singapore | HLSV | AAN51974.2 | Malvaceae | Q |
| 11 | *Tobamovirus kyuri* | Kyuri green mottle mosaic virus | C1 | KGMMV | CAC44969.1 | Cucurbitaceae | K |
| 12 | *Tobamovirus maracujae* | Maracuja mosaic virus | Peru | MarMV | ABD94121.1 | Passifloraceae | R |
| 13 | *Tobamovirus obudae* | Obuda pepper virus | Ob | ObPV | BAA02703.1 | Solanaceae | Q |
| 14 | *Tobamovirus odontoglossi* | Odontoglossum ringspot virus | Korean Cy-1 | ORSV | CAA57645.1 | Orchidaceae | R |
| 15 | *Tobamovirus paprikae* | Paprika mild mottle virus | Japanese | PaMMV | BAC07269.1 | Solanaceae | Q |
| 16 | *Tobamovirus passiflorae* | Passion fruit mosaic virus | USA | PFMV | AEF01561.1 | Passifloraceae | R |
| 17 | *Tobamovirus capsici* | Pepper mild mottle virus | S | PMMoV | AAB02337.1 | Solanaceae | R |
| 18 | *Tobamovirus plumeriae* | Plumeria mosaic virus | Plu-Ind-1 | PluMV | AIA57393.1 | Apocynaceae | R |
| 19 | *Tobamovirus muricaudae* | Rattail cactus necrosis-associated virus | South Korea | RCNaV | AEV53623.1 | Cactaceae | Q |
| 20 | *Tobamovirus rehmanniae* | Rehmannia mosaic virus | Henan | RheMV | ABN59377.1 | Orobanchaceae | R |
| 21 | *Tobamovirus plantagonis* | Ribgrass mosaic virus | Kons.1105-R14 | RMV | AEI69662.1 | Plantaginaceae | Q |
| 22 | *Tobamovirus streptocarpi* | Streptocarpus flower break virus | Willingmann | SFBV | CAJ13999.1 | Gesneriaceae | Q |
| 23 | *Tobamovirus mititessellati* | Tobacco mild green mosaic virus | Solis | TMGMV | AAA47937.1 | Solanaceae | Q |
| 24 | *Tobamovirus tabaci* | Tobacco mosaic virus | variant 1 | TMV | QBQ57807.1 | Solanaceae | R |
| 25 | *Tobamovirus fructirugosum* | Tomato brown rugose fruit virus | Tom1-Jo | TBRFV | ALP48479.1 | Solanaceae | R |
| 26 | *Tobamovirus tomatotessellati* | Tomato mosaic virus | Queensland | ToMV | AAK17991.1 | Solanaceae | R |
| 27 | *Tobamovirus maculatessellati* | Tomato mottle mosaic virus | MX5 | ToMMV | AGU13204.1 | Solanaceae | R |

| | | | | | | | |
|---|---|---|---|---|---|---|---|
| 28 | *Tobamovirus tropici* | Tropical soda apple mosaic virus | Okeechobee | TSAMV | ANF29783.1 | Solanaceae | R |
| 29 | *Tobamovirus rapae* | Turnip vein-clearing virus | OSU | TVCV | AAC02785.1 | Brassicaceae | Q |
| 30 | *Tobamovirus wasabi* | Wasabi mottle virus | Shizuoka | WMoV | BAB82443.1 | Brassicaceae | Q |
| 31 | *Tobamovirus anthocercis* | Yellow tailflower mild mottle virus | Cervantes | YTMMV | AGZ63348.1 | Solanaceae | Q |
| 32 | *Tobamovirus youcai* | Youcai mosaic virus | | YoMV | AAB60601.1 | Brassicaceae | Q |
| 33 | *Tobamovirus cucurbitae* | Zucchini green mottle mosaic virus | K | ZGMMV | CAC82485.1 | Cucurbitaceae | K |

### Supplementary Text S1: Python script for radial density plot

```
import mrcfile
import numpy as np
import matplotlib.pyplot as plt
import matplotlib as mpl

# --- GLOBAL PUBLICATION STYLE ---
mpl.rcParams.update({
    "font.family": "Arial",
    "font.size": 11,
    "axes.linewidth": 1,
    "xtick.direction": "out",
    "ytick.direction": "out",
})

def get_3d_cylindrical_profile(file_path, max_radius_ang=120.0, dr_ang=0.5, auto_center=True):
    """
    Computes a 3D cylindrical radial density profile along the Z-axis (helical axis).
    Uses exterior bulk solvent (105-115 Å) for baseline subtraction.
    """
    with mrcfile.open(file_path) as mrc:
        volume = mrc.data.astype(np.float64)
        voxel_size = float(mrc.voxel_size.x)

    nz, ny, nx = volume.shape

    # Check for density sign inversion
    if np.abs(np.min(volume)) > np.max(volume):
        volume = -volume

    # Determine X/Y center along the central helical axis
    if auto_center:
        threshold = np.max(volume) * 0.2
        masked_vol = np.where(volume > threshold, volume, 0)

        y_indices, x_indices = np.indices((ny, nx))
        total_mass = np.sum(masked_vol)

        if total_mass > 0:
            cx = np.sum(x_indices * np.sum(masked_vol, axis=0)) / total_mass
            cy = np.sum(y_indices * np.sum(masked_vol, axis=0)) / total_mass
        else:
            cx, cy = (nx - 1) / 2.0, (ny - 1) / 2.0
    else:
        cx, cy = (nx - 1) / 2.0, (ny - 1) / 2.0

    # 2D distance map from the central Z-axis
    y_idx, x_idx = np.indices((ny, nx))
    r_dist_2d = np.sqrt((x_idx - cx)**2 + (y_idx - cy)**2) * voxel_size

    # Radial Binning Setup
    r_bin_edges = np.arange(0, max_radius_ang + dr_ang, dr_ang)
    radii = (r_bin_edges[:-1] + r_bin_edges[1:]) / 2.0

    # 3D Cylindrical Shell Integration
    densities = []
```

```
    for i in range(len(r_bin_edges) - 1):
        r_min, r_max = r_bin_edges[i], r_bin_edges[i+1]
        mask_2d = (r_dist_2d >= r_min) & (r_dist_2d < r_max)

        if np.any(mask_2d):
            shell_densities = volume[:, mask_2d]
            densities.append(np.mean(shell_densities))
        else:
            densities.append(0.0)

    radii = np.array(radii)
    densities = np.array(densities)

    # Baseline Correction using Exterior Bulk Solvent (105 Å to 115 Å)
    solvent_mask = (radii >= 105.0) & (radii <= 115.0)
    solvent_baseline = np.mean(densities[solvent_mask]) if np.any(solvent_mask) else 0.0
    corrected = densities - solvent_baseline

    return radii, corrected


# --- DATA PROCESSING ---
r_tommv, d_tommv = get_3d_cylindrical_profile('tommv_centered.mrc', max_radius_ang=120.0, dr_ang=0.5,
auto_center=True)
r_cgmmv, d_cgmmv = get_3d_cylindrical_profile('cgmmv_311_resampled.mrc', max_radius_ang=120.0,
dr_ang=0.5, auto_center=True)

# Peak Normalization (Relative to CGMMV Peak Density)
cgmmv_max = np.max(d_cgmmv)
if cgmmv_max > 0:
    d_cgmmv_norm = d_cgmmv / cgmmv_max
    d_tommv_norm = d_tommv / cgmmv_max
else:
    d_cgmmv_norm = d_cgmmv
    d_tommv_norm = d_tommv

# --- PLOTTING ---
fig, ax = plt.subplots(figsize=(6.5, 4.5))

color_cgmmv = "#33d17A"
color_tommv = "#9141ac"

# Trace Plotting (No internal labels/legends)
ax.plot(r_cgmmv, d_cgmmv_norm, color=color_cgmmv, linewidth=2)
ax.plot(r_tommv, d_tommv_norm, color=color_tommv, linewidth=2)

# Zero Baseline Standard Reference Line
ax.axhline(0, color='black', linewidth=0.8, linestyle='--')

# Shaded Region Spans (Clean background accents)
ax.axvspan(18, 22, color='gray', alpha=0.15)
ax.axvspan(85, 95, color='orange', alpha=0.15)

# Dynamic Y-Axis Autoscaling to Prevent Trace Clipping
min_density = min(np.min(d_cgmmv_norm), np.min(d_tommv_norm))
max_density = max(np.max(d_cgmmv_norm), np.max(d_tommv_norm))
```

```
ax.set_xlabel('Radius (Å)')
ax.set_ylabel('Normalized Relative Density')
ax.set_xlim(0, 120)
ax.set_ylim(min_density - 0.05, max_density + 0.1)

# Clean Axis Spines
ax.spines['top'].set_visible(False)
ax.spines['right'].set_visible(False)

plt.tight_layout()

# Save High-Resolution TIFF (600 DPI, LZW Compression)
plt.savefig(
    "radial_density_publication.tiff",
    dpi=600,
    bbox_inches="tight",
    pil_kwargs={"compression": "tiff_lzw"}
)

plt.show()
```

### Supplementary Text S2: Python script for Assembly deprotonation profiles and Subunit surface electrostatics

```
import numpy as np
import matplotlib.pyplot as plt

# ========================
# DATA (Per-subunit pKa values in assembly context)
# ========================

cgmmv = {
    "ASP43":3.54,"ASP58":4.49,"ASP65":3.26,"ASP89":2.88,"ASP99":3.56,
    "ASP116":4.00,"ASP117":3.86,"ASP127":4.05,"ASP138":4.95,"ASP141":3.71,
    "GLU47":9.19,"GLU96":5.27,"GLU107":5.12,"GLU125":4.53,
    "GLU131":4.85,"GLU146":3.85,"GLU155":4.49
}

tommv = {
    "ASP20":4.43,"ASP65":3.72,"ASP78":4.87,"ASP89":4.40,"ASP110":4.05,
    "ASP116":3.57,"ASP117":4.09,
    "GLU23":6.05,"GLU51":8.52,"GLU96":5.10,"GLU98":6.26,
    "GLU107":4.51,"GLU132":4.58,"GLU146":4.50
}

tmv = {
    "ASP19":4.97,"ASP64":3.54,"ASP66":3.99,"ASP77":4.52,"ASP88":3.91,
    "ASP109":4.55,"ASP115":4.16,"ASP116":4.53,
    "GLU22":4.72,"GLU50":4.28,"GLU95":4.94,"GLU97":5.37,
    "GLU106":4.71,"GLU131":6.84,"GLU145":4.77
}

# ========================
# FUNCTIONS
# ========================

def charge_acid(pKa, pH):
    return -1 / (1 + 10**(pKa - pH))

def cumulative_acidic_charge(res_dict, pH):
    return np.array([sum(charge_acid(pKa, ph) for pKa in res_dict.values()) for ph in pH])

# ========================
# CALCULATION
# ========================

pH = np.linspace(2, 10, 500)

cg_charge = cumulative_acidic_charge(cgmmv, pH)
to_charge = cumulative_acidic_charge(tommv, pH)
tmv_charge = cumulative_acidic_charge(tmv, pH)

# ========================
# PLOT
# ========================
```

```
plt.figure(figsize=(6,5))

# Curves
plt.plot(pH, cg_charge, color="#33d17A", linewidth=2, label="CGMMV")
plt.plot(pH, to_charge, color="#9141ac", linewidth=2, label="ToMMV")
plt.plot(pH, tmv_charge, color="#e37817", linewidth=2, label="TMV")

# Axis labels
plt.xlabel("pH", fontsize=12)
plt.ylabel("Cumulative Acidic Charge / Subunit ($e$)", fontsize=12)

# Ticks
plt.minorticks_on()
plt.tick_params(axis='both', which='major', length=6, width=1)
plt.tick_params(axis='both', which='minor', length=3, width=0.8)

plt.grid(False)
plt.tight_layout()

# Save
plt.savefig("acidic_charge_titration_marked.tiff", dpi=600, format="tiff")
plt.show()
```

**Supplementary Text S3: Python script for hydrophobic patch calculation**

```
import numpy as np
from scipy.spatial.distance import pdist, squareform

# =============================================================
# INPUT FILES
# =============================================================
pdb_file = "cgmmv_local_assembly.pdb"

sasa_monomer_file = "cgmmv_monomer_sasa.txt"
sasa_assembly_file = "cgmmv_assembly_sasa.txt"

output_file = "cgmmv_delta_sasa_patches_central_chain_c.txt"

# =============================================================
# 9-MER ARCHITECTURE & PARAMETERS
# =============================================================
# Layer 1: L, M, N
# Layer 2: b, c (CENTRAL), d
# Layer 3: r, s, t
CENTRAL_CHAIN = "c"
NEIGHBOR_CHAINS = {"L", "M", "N", "b", "d", "r", "s", "t"}
ALL_9MER_CHAINS = NEIGHBOR_CHAINS | {CENTRAL_CHAIN}

DELTA_SASA_CUTOFF = 25.0   # Å²
DIST_CUTOFF = 7.0          # Å
MIN_PATCH_SIZE = 2

hydrophobic_residues = {
    "ALA", "VAL", "LEU", "ILE", "MET", "PHE", "TRP", "TYR", "PRO"
}

# =============================================================
# READ FREESASA FILE (SEQ FORMAT)
# =============================================================
def read_sasa(filename):
    data = {}
    with open(filename) as f:
        for line in f:
            if not line.startswith("SEQ"):
                continue
            parts = line.split()
            chain = parts[1]
            resnum = int(parts[2])
            resname = parts[3]
            sasa = float(parts[5])
            data[(chain, resnum)] = (resname, sasa)
    return data

# =============================================================
# READ PDB CA COORDINATES
# =============================================================
def read_coords(pdb_file):
    coords = {}
    with open(pdb_file) as f:
        for line in f:
```

```
            if line.startswith("ATOM") and line[12:16].strip() == "CA":
                chain = line[21]
                resnum = int(line[22:26])
                x = float(line[30:38])
                y = float(line[38:46])
                z = float(line[46:54])
                coords[(chain, resnum)] = np.array([x, y, z])
    return coords

# ============================================================
# GET INTERFACE RESIDUES (TARGETED TO CHAIN C & ITS 8 NEIGHBORS)
# ============================================================
def get_interface_residues(mono, assembly, coords, central_chain, valid_chains):
    residues = []

    # Map monomer SASA by residue number (ignoring monomer chain ID differences)
    mono_by_resnum = {resnum: sasa for (c, resnum), (name, sasa) in mono.items()}

    for (chain_a, resnum_a), (resname_a, sasa_assembly) in assembly.items():

        # 1. Filter out chains outside our 9-mer definition
        if chain_a not in valid_chains:
            continue

        # 2. Skip residues lacking C-alpha coordinates
        if (chain_a, resnum_a) not in coords:
            continue

        # 3. Skip non-hydrophobic residues
        if resname_a not in hydrophobic_residues:
            continue

        # 4. Check against monomer SASA
        if resnum_a not in mono_by_resnum:
            continue

        sasa_mono = mono_by_resnum[resnum_a]
        delta = sasa_mono - sasa_assembly

        # 5. Retain if burial threshold is met
        if delta > DELTA_SASA_CUTOFF:
            residues.append({
                "chain": chain_a,
                "resnum": resnum_a,
                "resname": resname_a,
                "delta": delta,
                "coord": coords[(chain_a, resnum_a)],
                "is_central": (chain_a == central_chain)
            })

    return residues

# ============================================================
# CLUSTER PATCHES (FOCUSED ON CENTRAL CHAIN C INTERFACES)
# ============================================================
def cluster_patches(residues, central_chain):
```

```
    if len(residues) < 2:
        return []

    coords_arr = np.array([r["coord"] for r in residues])
    dist = squareform(pdist(coords_arr))
    adj = dist < DIST_CUTOFF

    visited = set()
    patches = []

    for i in range(len(residues)):
        if i in visited:
            continue

        stack = [i]
        comp = []

        while stack:
            j = stack.pop()
            if j in visited:
                continue

            visited.add(j)
            comp.append(j)

            neighbors = np.where(adj[j])[0]
            stack.extend(neighbors)

        if len(comp) < MIN_PATCH_SIZE:
            continue

        patch_res = [residues[k] for k in comp]

        # FILTER: Keep only clusters containing at least one residue from central Chain 'c'
        has_central = any(r["chain"] == central_chain for r in patch_res)
        if not has_central:
            continue

        area = sum(r["delta"] for r in patch_res)
        chains = set(r["chain"] for r in patch_res)

        patches.append({
            "size": len(patch_res),
            "area": area,
            "chains": chains,
            "type": "interface" if len(chains) > 1 else "single-chain",
            "residues": patch_res
        })

    return sorted(patches, key=lambda x: x["area"], reverse=True)

# ============================================================
# OUTPUT WRITER
# ============================================================
def write_output(residues, patches, central_chain):
    with open(output_file, "w") as f:
```

```
    f.write("DELTA SASA HYDROPHOBIC PATCH ANALYSIS (CENTRAL CHAIN C FOCUSED)\n")
    f.write("=" * 68 + "\n")
    f.write(f"Target Central Chain : Chain {central_chain} (Layer 2)\n")
    f.write("Layer 1 Neighbors    : Chains L, M, N\n")
    f.write("Layer 2 Neighbors    : Chains b, d\n")
    f.write("Layer 3 Neighbors    : Chains r, s, t\n")
    f.write("-" * 68 + "\n")
    f.write(f"Total Candidate Interface Residues (ΔSASA > {DELTA_SASA_CUTOFF} Å²): {len(residues)}\n")
    f.write(f"Unique Chain 'c' Patches Identified: {len(patches)}\n\n")

    interface = [p for p in patches if p["type"] == "interface"]
    single = [p for p in patches if p["type"] == "single-chain"]

    f.write(f"Inter-subunit Interface Patches: {len(interface)}\n")
    f.write(f"Intrachain Patches (Chain c only): {len(single)}\n\n")

    if patches:
        areas = [p["area"] for p in patches]
        f.write("SUMMARY STATISTICS:\n")
        f.write(f"Mean Patch ΔSASA    : {np.mean(areas):.2f} Å²\n")
        f.write(f"Largest Patch ΔSASA: {np.max(areas):.2f} Å²\n\n")

    for i, p in enumerate(patches):
        f.write(f"Patch {i+1}\n")
        f.write("-" * 40 + "\n")
        f.write(f"Type              : {p['type']}\n")
        f.write(f"Residue Count     : {p['size']}\n")
        f.write(f"Buried Area (Δ)   : {p['area']:.2f} Å²\n")
        f.write(f"Involved Chains   : {', '.join(sorted(p['chains']))}\n")
        f.write("Residues:\n")

        for r in p["residues"]:
            f.write(f"  {r['resname']:3s} Chain {r['chain']} Res {r['resnum']:3d}  (ΔSASA = {r['delta']:.2f} Å²)\n")

        f.write("\n")

# ============================================================
# MAIN EXECUTION
# ============================================================
mono = read_sasa(sasa_monomer_file)
assembly = read_sasa(sasa_assembly_file)
coords = read_coords(pdb_file)

residues = get_interface_residues(mono, assembly, coords, CENTRAL_CHAIN, ALL_9MER_CHAINS)
patches = cluster_patches(residues, CENTRAL_CHAIN)

write_output(residues, patches, CENTRAL_CHAIN)

print(f"\nAnalysis Complete for Central Chain '{CENTRAL_CHAIN}'")
print(f"Inter-subunit Patches: {len([p for p in patches if p['type'] == 'interface'])}")
print(f"Intrachain Patches: {len([p for p in patches if p['type'] == 'single-chain'])}")
```

**Supplementary Text S4: Maximum Likelihood phylogenetic tree for Tobamoviruses with branch length and bootstrap support values (Newick format)**

(((((((((RheMV:0.02674090,TMV:0.00975428)0.9920:0.07413647,TBRFV:0.02502625)0.9540:0.05033618,(ToMV:0.04918217,ToMMV:0.03910736)0.9540:0.04090258)0.9740:0.07063907,BPMV:0.06422625)0.6440:0.01795459,ORSV:0.19040340)0.6540:0.04119022,(PMMoV:0.09442070,TSAMV:0.06320899)0.9520:0.05733473)0.7680:0.05081481,(TMGMV:0.12472202,((BrMMV:0.20133629,YTMMV:0.24704646)0.3340:0.01816639,(ObPV:0.07333763,PaMMV:0.03489160)1.0000:0.13828692)0.5900:0.06710680)0.7320:0.07032120)0.9860:0.22420741,(SFBV:0.27424104,(RMV:0.01628214,(TVCV:0.03978811,(WMoV:0.04021994,YoMV:0.03994297)0.7640:0.04220097)0.4480:0.02607269)1.0000:0.20917566)0.9840:0.19211904)0.7560:0.19303351,(CYMV:0.48386952,((HLFPV:0.11715134,HLSV:0.15175890)0.9900:0.23740089,((FrMV:0.11066740,PluMV:0.02999638)1.0000:0.46878631,((CGMMV:0.35596122,CMoV:0.33400147)0.5560:0.10717725,(ZGMMV:0.09676103,(CFMMV:0.10504231,KGMMV:0.10136789)0.7420:0.05258734)1.0000:0.40029998)0.6720:0.17893094)0.1600:0.07385383)0.1460:0.05566137)0.3740:0.04893273)0.3500:0.11751652,(MarMV:0.10569816,PFMV:0.14232960)1.0000:0.46262780,(CMMoV:0.28503036,RCNaV:0.40545506)1.0000:0.64919631);
